# Individual heartbeats track distinct prediction processes during human probabilistic learning

**DOI:** 10.64898/2026.08.19.745688

**Authors:** Maria Azanova, Lina Skora, Alina Studenova, Esra Al, Vadim V. Nikulin, Arno Villringer

## Abstract

Heart rate continuously adjusts to accommodate perception and action, and these shifts are frequently explained through predictive processes. Yet direct evidence that interbeat intervals exhibit graded scaling with prediction remains limited. Here we investigate millisecond-resolved physiological signatures of prediction processing using a mechanistically constrained analysis of beat-to-beat cardiac dynamics. We analysed trial-by-trial electrocardiogram and electroencephalogram recordings from 34 participants performing a probabilistic learning task. We quantified stimulus-locked cardiac responses as changes between consecutive interbeat intervals and accounted for cardiac phase at feedback. This single-beat approach separated anticipatory slowing, stimulus-locked parasympathetic brake, and subsequent acceleration. Anticipatory deceleration and rebound acceleration scaled with model-derived expectations, whereas the second heartbeat after feedback tracked signed prediction errors and outcome valence, particularly when feedback occurred early in the cardiac cycle. Peak stimulus-locked cardiac deceleration covaried with parietal P3b rather than prediction features. Thus, individual cardiac cycles carry separable signatures of anticipation, orienting, and feedback-based updating. These findings demonstrate how predictive processing propagates into human autonomic physiology on a beat-to-beat timescale and provide an interpretable, mechanistically grounded framework for quantifying brain-body co-modulation during adaptive behaviour.

## Introduction

The heart slows down when a biathlete hits a target^1^, an infant masters a vocalisation^2^, or a pianist hits a wrong note^3^. Prominent theories posit that fluctuations in heartbeat intervals serve moment-to-moment demands of an organism: the heart slows down to enhance perception, then speeds up for action, underpinning successful behavioural adaptation^4–7^. Behavioural adaptation relies on the processing of expectations, outcomes, and resulting prediction errors^8^, but how fast, to what degree, and if at all moment-to-moment heart rate adjustments track prediction remains unclear.

Over the last 60 years, heart rate modulations have been observed in a range of contexts: threat^9^, anticipation^10,11^, error^12–16^, and in general detection of unexpected events^17–19^. Prediction is the common denominator of these phenomena^19^: unexpected or salient events elicit a pattern of cardiac deceleration followed by acceleration, mirroring phasic changes in the autonomic balance and momentary bodily regulation^9^. However, cardiac responses are often treated as relatively coarse changes in heart rate across broad time windows, or as binary acceleration–deceleration effects. This makes it difficult to determine whether the heart merely responds to salient events, or whether cardiac dynamics scale with the degree of prediction: how strongly an outcome deviates from expectation, and if this deviation is better or worse than expected.

In theory, the body should be adjusted in proportion to environmental demands^20^. Activity in regions involved in both performance monitoring and autonomic control, including the anterior cingulate cortex and insula, scales continuously with expectations and prediction errors^21–27^. Therefore, at least part of this processing should be reflected in bodily signals. The precision of brain-body co-modulation in adaptive behaviours could vary among individuals and be involved in maladaptive scenarios^28–32^. In fact, such co-modulations can be viewed as microstates in the brain-body state space, limited by larger, overarching meso- and macrostates^33^. For instance, in anxiety, uncertainty evokes increased autonomic arousal and inadequate employment of bodily resources, which are reflected, among others, in a disrupted heart-brain connection^31,34^. However, before we can view the extent of brain-body co-modulation as a biomarker^28^, it is necessary to establish a robust and interpretable approach to the underlying physiological phenomenon.

A clear test of this proposition has been limited by three methodological problems. First, prediction-related cardiac effects are expected to be subtle, especially in non-salient contexts, and thus difficult to detect. Second, the failure to distinguish ongoing and transient cardiac dynamics complicates the interpretation of evoked cardiac responses, i.e., both sympathetic and parasympathetic influences are always reflected in each interbeat interval along with individual physiological traits^35–37^. Third, the smoothing of effects over longer time intervals is problematic for studying phasic changes in parasympathetic activity, which occur on a half- to one-second scale (in contrast to three-second-long sympathetic influences)^38^ and also vary depending on the cardiac phase at stimulus onset^39,40^. An analysis approach that accounts for mechanistic constraints on vagally-driven cardiac changes and treats the interbeat interval as the unit of physiological change may improve inference and sensitivity to subtle effects.

Here, we quantify stimulus-locked cardiac responses as changes between consecutive interbeat intervals, while also accounting for cardiac phase at stimulus onset. This way, we avoid conflating slower tonic fluctuations with rapid phasic responses, tuning potential interpretations to transient parasympathetic changes mediated by central cholinergic activity and the vagus nerve on the periphery^38^.

This physiologically informed framework clearly separates the three components of the stimulus-locked cardiac response. First, lengthening of interbeat intervals before feedback can be interpreted as anticipatory parasympathetic activation. Second, peak cardiac slowing immediately after feedback reflects a stimulus-locked parasympathetic brake, often linked to orienting and enhanced perception as well as potential network segregation but not yet stimulus processing. Third, subsequent shortening of interbeat intervals reflects parasympathetic withdrawal, and with time likely also sympathetic influences, signifying preparation for action or updating associated with network integration. Through this lens, we can ask which component of the cardiac response is modulated, at which heartbeat, and by which computational feature of prediction.

We apply this framework to a probabilistic approach-avoidance learning task with concurrent electrocardiogram and electroencephalogram recordings from 34 participants. The task used mild positive and negative symbolic feedback rather than monetary incentives or threat, providing a conservative test of whether prediction-related transient cardiac dynamics can be detected in a relatively non-salient learning context in the young, healthy, moderately sized sample. We fitted computational learning models to estimate trial-wise expectation, signed and unsigned prediction error, and outcome valence. We then tested whether changes in four heartbeats surrounding feedback, the last one before and the first three after, scaled with these variables. By simultaneously considering expectations, outcomes, and prediction errors, we covered the full prediction-processing loop and asked how it propagates to the heart from beat to beat.

We additionally considered P300 (frontal P3a and parietal P3b components) in order to distinguish the influence of attentional orienting: studies show that peak cardiac deceleration at the moment of orienting to a meaningful stimulus does not reflect its motivational features but rather the degree of attention to it, often associated with P300 amplitude^12,41–43^.

As a result, we demonstrate a physiological signature of prediction processing in human beat-to-beat cardiac dynamics: individual cardiac cycles track distinct computational features of learning. The effects on interbeat interval changes manifested on a scale of several milliseconds and were supported by strong evidence. Anticipatory deceleration and rebound acceleration scaled with expectation: greater expected loss led to relative increases in interbeat intervals before feedback and decreases after. Cardiac acceleration at the second heartbeat post-feedback reflected features of outcome: the heart accelerated less after negative outcomes and gradually less with negative signed or larger unsigned prediction errors, with stronger effects when feedback was presented early in the cardiac phase. The unsigned error result provided weaker evidence. Moreover, peak stimulus-locked cardiac deceleration positively covaried with parietal P3b but not frontal P3a amplitudes. Thus, prediction processing is not only visible in choices and neural responses; it is also expressed in the timing of individual heartbeats.

## Results

### Beat-to-beat analysis resolves feedback-locked cardiac dynamics

We analysed a simultaneous electrocardiogram and electroencephalogram dataset from 34 participants performing a probabilistic approach–avoidance learning task (Fig. 1A). After preprocessing, 9808 trials entered the cardiac analyses, with an average of 288 trials per participant. We quantified cardiac dynamics as changes between consecutive interbeat intervals (IBI changes): positive values indicate relative slowing and negative values indicate relative speeding of the current heartbeat compared to the previous one (Fig. 1B).

**Figure 1.**
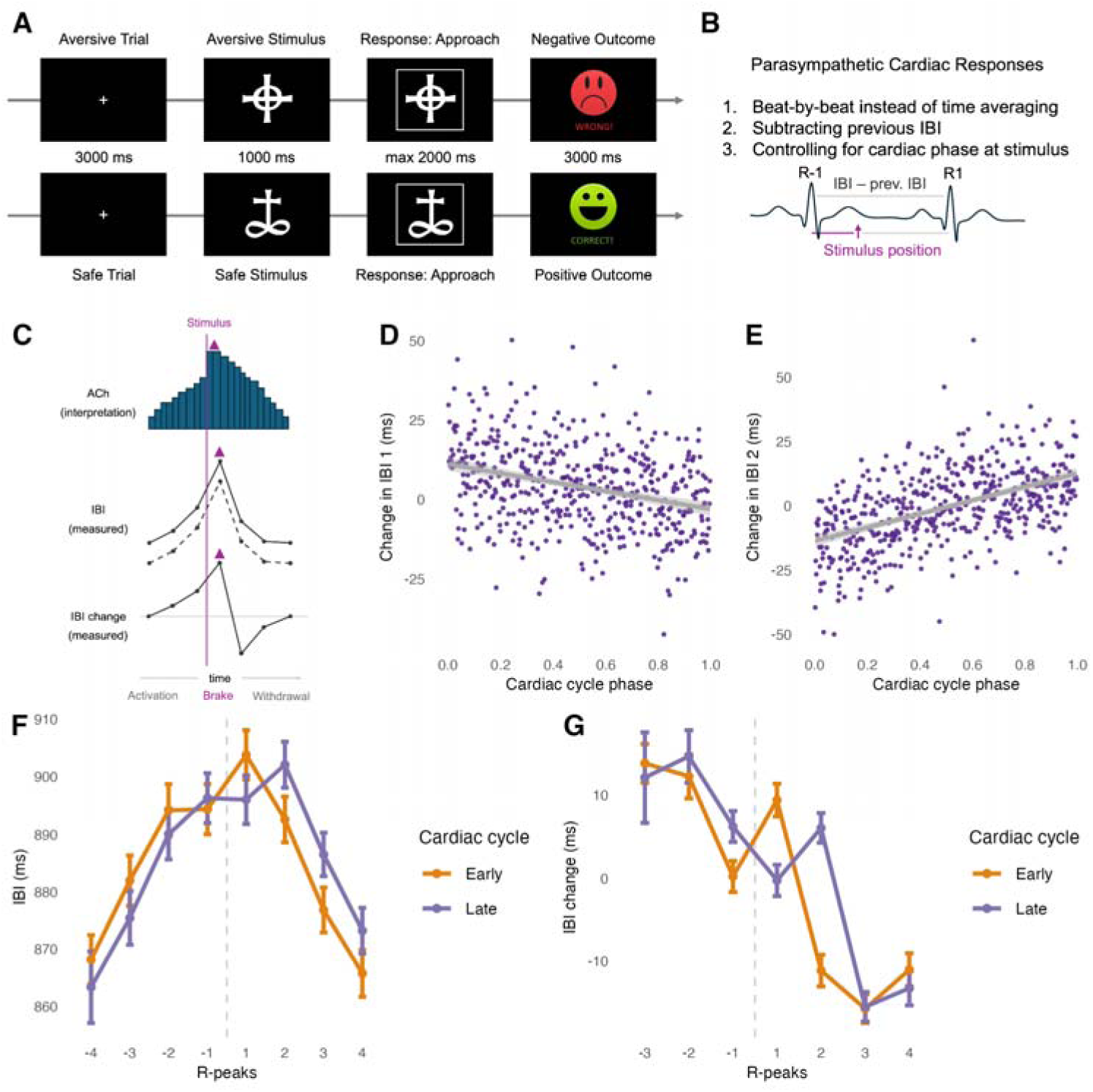
Experimental paradigm and beat-to-beat framework for resolving stimulus-locked cardiac dynamics. (A) Structure of the probabilistic approach-avoidance learning task. (B) Overview of the cardiac analysis framework - the approach aimed at increased precision in identifying immediate vagally-driven changes in heart rate. (C) Schematic interpretation of the triphasic cardiac response. Anticipatory parasympathetic activation produces progressive IBI lengthening before the stimulus; the stimulus-locked parasympathetic brake (i.e., pronounced increase in spiking) produces peak cardiac deceleration; and subsequent parasympathetic withdrawal produces IBI shortening and cardiac acceleration. Upper bars illustrate the hypothesised cholinergic activity underlying these dynamics, whereas the lower traces show corresponding measured IBI and consecutive IBI-change profiles. Magenta triangles mark the moment of stimulus-locked parasympathetic brake associated with pronounced short-term increase in ACh spiking (in the upper ACh panels) and the one heartbeat that reflects this brake (in the two lower IBI panels) (D, E) Empirical relationship between cardiac-cycle phase and the change in the first and second post-feedback IBI, respectively. Cardiac phase is expressed from 0 to 1, with lower values indicating feedback presented earlier in the cardiac cycle. Points represent 500 binned observations averaged across participants and trials; grey lines show linear trends with 95% confidence intervals. (F) Mean IBI across the four heartbeats before and four heartbeats after feedback, shown separately for feedback presented early or late in the cardiac cycle. R1 is the IBI containing feedback presentation. (G) Mean change in IBI relative to the preceding heartbeat for the three heartbeats before and four heartbeats after feedback, shown separately for early- and late-cycle trials. Positive values indicate relative cardiac slowing and negative values indicate relative acceleration. In F and G, points show means and error bars show 95% confidence intervals; the vertical dashed line marks feedback presentation. Early- and late-cycle trials correspond to feedback presented in the first and last third of the cardiac cycle, respectively. Abbreviations: ACh, acetylcholine; IBI, interbeat interval; ms, milliseconds.

We focused on four heartbeats surrounding feedback: the last heartbeat before feedback, termed R-1, in the prestimulus window, and the first three heartbeats after feedback, termed R1, R2, and R3 and corresponding to the feedback presentation window. R1 interbeat interval contains feedback presentations. Using single-trial mixed models, we tested the effects of outcome valence, model-derived expectation, signed prediction error (PE), unsigned PE, and P3b/P3a amplitudes at Pz and FCz channels, respectively. Evidence was quantified using Bayes factors (BF), with regions of practical equivalence (ROPE) defined so that effects below 1 ms across the full predictor range were treated as practically negligible. We also corrected for false discovery rate (FDR) based on posterior-ROPE overlaps. Here, we present only findings that survived it. In Supplementary Information, we also report on the additional three heartbeats that were more distant from feedback: R-3, R-2, and R4, covering the full length of a trial. These sensitivity analyses complemented the main analyses for R-1, R1, R2, and R3 by showing temporal specificity of observed effects.

First, we verified that the average cardiac response showed a triphasic beat-to-beat structure (Fig. 1CFG). Before feedback, R-3 and R-2 slowed down relative to previous intervals, lengthening on average by 11.7 ms and 11.8 ms, respectively. After feedback, R3 and R4 sped up, shortening by 13.3 ms and 10.2 ms, respectively. By contrast, the beats of interest R-1, R1 and R2 showed no clear slowing or speeding when averaged across all trials. This was explained by strong dependence on the cardiac phase at feedback (Fig. 1DE). When feedback occurred early in the cardiac cycle, in the first third, R1 showed peak deceleration of 10.6 ms and R2 showed acceleration of 10.9 ms. When feedback occurred late in the cardiac cycle, in the last third, R1 showed no clear change, whereas R2 showed peak deceleration of 8.7 ms followed by a 14.5 ms acceleration at R3. R-1 did not change strongly compared to R-2 but showed sustained deceleration. Feedback presentations were evenly distributed across the cardiac cycle and did not depend on trial number (BF in ROPE 3268807061), indicating that the phase-dependent effects were not caused by systematic entrainments of cardiac timing. Full heartbeat-wise estimates are reported in Tables S1-3.

Therefore, cardiac phase at feedback shaped the expression of immediate stimulus-locked dynamics for R1 and R2, as predicted by the literature^5,17,44^. The phase dependence was accounted for in subsequent analyses: we modelled prediction-related effects on IBI changes and included cardiac phase as a mechanistic moderator. The trial-wise cardiac dynamics provide the background against which prediction-related modulations should be interpreted: a heartbeat may slow down on average but slow down more for one kind of feedback, or speed up on average but speed up less for another kind of feedback.

### Expectations shape anticipatory and rebound cardiac dynamics

We tested whether IBI changes were gradually modulated by ongoing model-derived expectations: a continuous probabilistic value associated with the chosen option before feedback presentation. If cardiac dynamics reflect preparation for expected outcomes, R-1 should already contain information about upcoming feedback. Conversely, if expectation also shapes subsequent feedback integration, its influence may extend into later post-feedback dynamics.

R-1 and R3 showed pronounced but opposite scaling with expectation. R-1 lengthened more when more negative outcomes were expected, indicating stronger anticipatory slowing before worse expected feedback (BF 67). The coefficient converted back to milliseconds was -3.3 ms per unit of expectation, with a 95% highest-density interval (HDI) from -4.3 to - 1.3 ms. Because expectation ranged from -1 to 1, the estimated full-range modulation was on the order of -8.6 to -2.6 ms. R3 showed the opposite effect: it shortened more when more negative outcomes were expected prior to feedback, with a coefficient of 2.0 ms per unit of expectation and a 95% HDI from 1.4 to 4.3 ms (BF 108). Thus, expectation shaped both the anticipatory component of the cardiac response and the later rebound acceleration (Fig. 2A- D, Table S4, Fig. S1).

**Figure 2.**
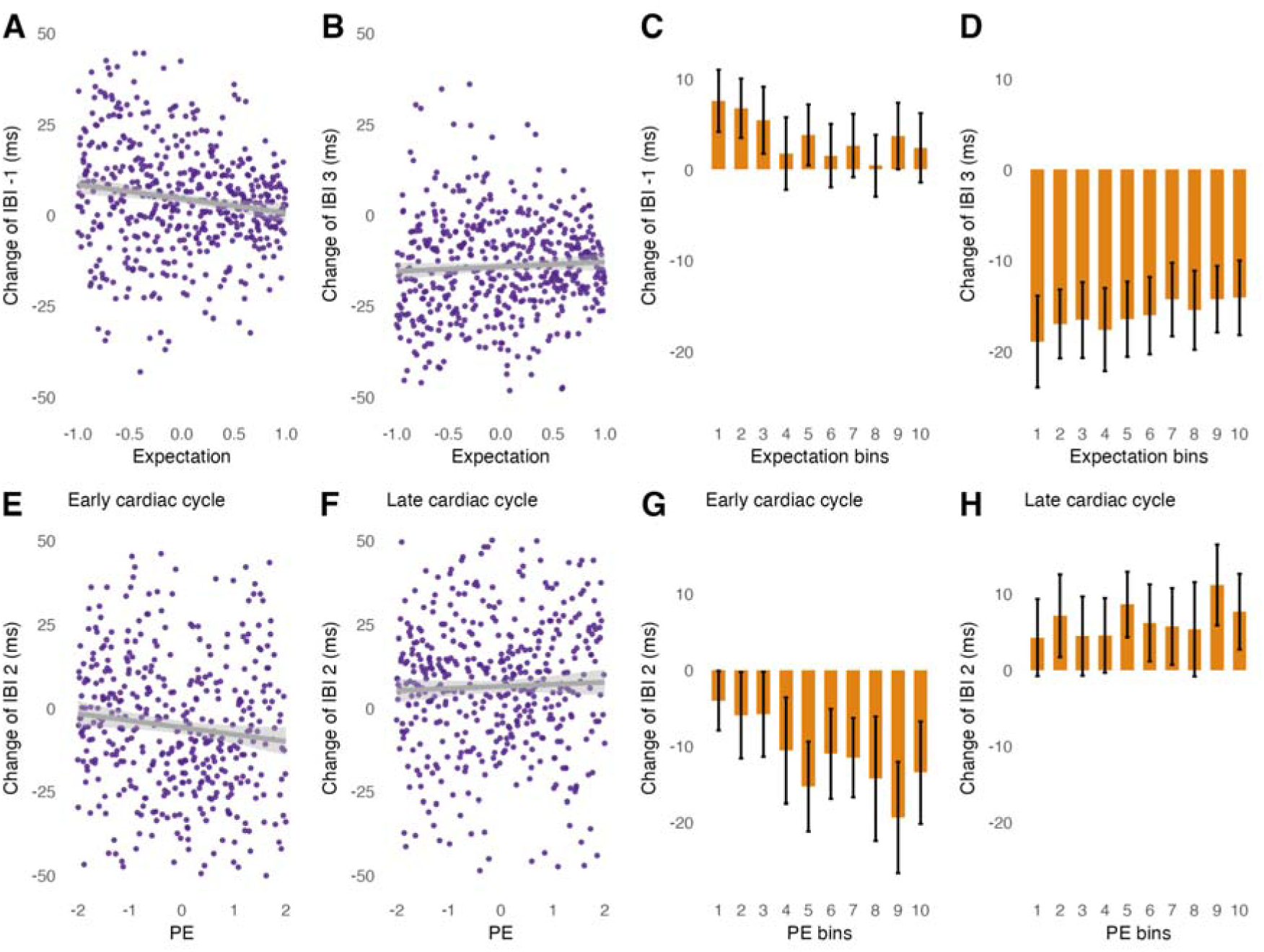
Expectations shape anticipatory and rebound cardiac dynamics, whereas prediction errors modulate the second heartbeat after feedback, especially when feedback appears early in the cardiac cycle. (A, B) Average change in interbeat interval (IBI), relative to the preceding heartbeat, as a function of model-derived expectation for the last heartbeat before feedback, R-1 (A), and the third heartbeat after feedback, R3 (B). Dots represent 500 binned observations averaged across participants and trials. Grey lines show linear trends and shaded areas indicate their 95% confidence intervals. More negative expectations were associated with stronger anticipatory slowing at R-1 and stronger rebound acceleration at R3. (C, D) Mean change in IBI at R-1 (C) and R3 (D) across ten expectation bins obtained by within-participant quantile splits. Bars show means and error bars indicate 95% confidence intervals. (E, F) Average change in IBI at the second heartbeat after feedback, R2, as a function of signed prediction error (PE), shown separately for trials in which feedback occurred early (E) or late (F) in the cardiac cycle. Dots represent 500 binned observations averaged across participants and trials; grey lines show linear trends with 95% confidence intervals. More negative PE was associated with a longer R2, particularly when feedback occurred early in the cardiac cycle. (G, H) Mean change in IBI at R2 across ten PE bins obtained by within-participant quantile splits, shown separately for systole (G) and diastole (H) trials. Bars show means and error bars indicate 95% confidence intervals. Positive IBI-change values indicate relative cardiac slowing, whereas negative values indicate relative acceleration. Abbreviations: IBI, interbeat interval; ms, milliseconds; PE, signed prediction error; R-1, last heartbeat before feedback; R2 and R3, second and third heartbeats after feedback.

These effects were temporally specific. R1 showed only moderate evidence for expectation (BF 3.8), and R2 did not show meaningful expectation effects (BF in ROPE 7.8). No interactions between cardiac phase and expectation were observed for any heartbeat.

These effects were reliable despite the modest participant sample and simple symbolic feedback. Posterior-predictive post hoc evidential power estimated from the current data indicated 99% power to detect strong evidence, BF > 10, for the R-1 expectation effect and >99.9% power for the R3 expectation effect in the present 34-participant sample. These simulations indicate that the observed expectation effects were detectable under the specified model and sampling assumptions.

### Signed prediction errors are reflected in the second heartbeat after feedback

Next, we tested whether feedback-related cardiac dynamics reflected signed PE. Signed PE is a difference between an observed outcome in a trial and expectation accumulated over previous trials: negative values indicate outcomes worse than expected, whereas positive values indicate outcomes better than expected. It is employed in updating beliefs about choice options, guiding a constant loop of interactions with the world^19,27^. Especially for heartbeats following feedback, it is critical to test how the integration of expectation and outcome is reflected in transient changes of the autonomic balance.

More negative PE was associated with a longer R2 compared to R1 (BF 16; -1.2 ms per PE unit; 95% HDI [-2.1, -0.4] ms). Because signed PE ranged from -2 to 2, the implied full-range difference in R2 change was approximately within 1.6-8.4 ms. The PE effect was substantially stronger when feedback occurred early in the cardiac cycle (interaction BF 20). Only among these trials, the coefficient was -2.4 ms per PE unit (BF 99; 95% HDI [-4.0, -1.1] ms). Across the full PE range, this corresponds to approximately 4.4-16.0 ms modulation of R2. This effect should be interpreted relative to the average dynamics of R2: in early cardiac-cycle trials, R2 shortened by approximately 10.9 ms on average, meaning that worse-than-expected outcomes reduced this acceleration after the feedback-locked brake in proportion to the degree of PE (Fig. 2E-H, Tables S5-6, Fig. S2). The observed modulations were temporally specific: other heartbeats showed no comparable evidence for signed prediction-error scaling, indicating that the outcome-expectation integration is a transient autonomic state, not merely a tonic arousal modulation, at least within the non-salient context.

Therefore, when feedback occurred early in the cardiac cycle, the second post-feedback heartbeat was, on average, in the acceleration phase and worse-than-expected outcomes reduced this acceleration (BF 99). When feedback occurred late in the cardiac cycle, the second heartbeat was in the deceleration phase and the prediction-error scaling was not supported by evidence (inconclusive BF in ROPE 2.8). This demonstrates why cardiac phase must be modelled: the same-numbered heartbeat can reflect different physiological processes depending on when feedback occurs within the cardiac cycle. The specificity of the feedback-locked effects to R2 and not other heartbeats shows why beat-by-beat analysis rather than time averaging increases precision for detecting transient effects.

The signed PE effect was robust in the present sample. Posterior-predictive post hoc evidential power indicated 96% power to detect strong evidence, BF > 10, for the main PE effect in the 34-participant dataset. Power for the PE-phase interaction was lower, 62% for BF > 3 and 43% for BF > 10, consistent with the reduced precision of interaction estimates. When the analysis was restricted to early cardiac-cycle trials, power for the PE effect was 79% for BF > 3 and 62% for BF > 10, reflecting decreased power when analysing thrice fewer trials. Thus, the main PE effect was detectable under the specified simulation framework, whereas the phase-specific interaction was estimated less precisely.

### Outcome valence and unsigned prediction error provide convergent but less reliable evidence

We then asked whether the R2 effect reflected signed prediction-error coding specifically, or whether it could be explained by simpler feedback features. Outcome valence distinguished negative from positive feedback, while unsigned PE captured surprise independent of valence.

Outcome valence also modulated R2. Negative outcomes lengthened R2 relative to positive outcomes by on average 1.8 ms (BF 11; 95% HDI [0.7, 3.9] ms). Exclusively in early cardiac-cycle trials, this effect was larger: negative outcomes lengthened R2 by 5.8 ms (BF 19; 95% HDI [1.0, 6.4] ms). Because R2 shortened by approximately 10.9 ms on average in these trials, negative outcomes reduced nearly half of the average acceleration. However, the outcome effect was less reliably powered than the signed PE effect. In the current 34- participant sample, post hoc evidential power was 59% for BF > 3 and 36% for BF > 10 for the main outcome effect, with 63% and 44% power for the outcome-by-phase interaction. Thus, outcome valence provided convergent evidence that negative feedback reduces post-feedback acceleration, but it was less robust than signed PE in the current dataset (Tables S7-8) and would have required a sample of approximately 100 participants to reach 93% power for strong evidence BF > 10.

Unsigned PE showed a related pattern but should be treated as secondary. Larger unsigned PE was also associated with 1.6-6.4 ms longer R2 (BF 26), particularly in early cardiac-cycle trials (BF 13), where the estimated full-range modulation was approximately 2.0-10.6 ms (Tables S9-10). This suggests that unsigned surprise may also contribute to post-feedback cardiac dynamics. However, posterior-predictive power analyses indicated that the effect was substantially less reliable than the expectation and signed PE effects. In the present 34- participant sample, the power to detect even moderate evidence, BF > 3, for the main absolute PE effect was only 36%. Even when restricted to early cardiac-cycle trials, power for BF > 3 was 49%. We therefore present absolute PE as convergent evidence that R2 is sensitive to feedback informativeness, but not as a primary result. Moreover, even in the reasonably large sample of 100 participants, this effect would still not be sufficiently powered for strong evidence BF > 10: it would only achieve 67%. This suggests that the effect of unsigned surprise may be more variable than its signed counterpart, perhaps depending on individual strategies or reflecting later sympathetically modulated arousal rather than transient vagal responses, and is therefore not captured as robustly with the beat-by-beat approach.

Together, the outcome, signed PE and unsigned PE analyses suggest that R2 is not a generic heartbeat marker of arousal. It is sensitive to feedback features, but the most reliable effect in the present dataset was signed PE: worse-than-expected outcomes reduced post-feedback acceleration in proportion to their deviation from expectation.

### Stimulus-locked peak cardiac deceleration covaries with parietal P3b

We also assessed whether neural feedback processing was related to stimulus-locked cardiac deceleration, typically interpreted as an orienting-related parasympathetic brake. We focused on the heartbeats where this brake was expressed most clearly: R1 when feedback occurred early in the cardiac cycle, and R2 when feedback occurred late in the cardiac cycle. We then asked whether cardiac deceleration covaried with the feedback-evoked P300 components: parietal P3b and frontal P3a.

Because EEG responses sorted by cardiac events can be contaminated by cardiac field artefacts, we took a conservative approach. We analysed P300 responses separately for early and late cardiac-cycle trials (first and last third of cardiac cycle) based on pseudotrial- corrected participant-level averages to assess whether the effects are present beyond heartbeat-locked artefacts. If we observed stable differences between high and low deceleration average responses, we complemented them with trial-wise inference.

In analyses of pseudotrial-corrected participant-level averages, only parietal P3b showed meaningful differences associated with the degree of deceleration at R1 in early cardiac-cycle trials and R2 in late cardiac-cycle trials. These effects were detectable with moderate evidence for R1 (BF 4.2) and strong evidence for R2 (BF 30). By contrast, frontal P3a did not show comparable evidence (BF in ROPE 13 and 3, respectively), so any apparent single-trial association would need to be interpreted cautiously given the possibility of residual cardiac field artefacts.

Furthermore, parietal P3b amplitude predicted trial-wise cardiac deceleration. In early cardiac-cycle trials, larger P3b amplitudes were associated with stronger R1 deceleration: a 1 µV increase in P3b amplitude corresponded to a 0.27 ms increase in R1 slowing (BF 19; 95% HDI [0.13, 0.66] ms). For an average P3b amplitude of approximately 5 µV, this corresponds to an estimated modulation of roughly 1–3 ms. In late cardiac-cycle trials, the P3b association with R2 was stronger: a 1 µV increase in P3b amplitude corresponded to a 0.47 ms increase in R2 slowing (BF 412; 95% HDI [0.26, 0.79] ms), corresponding to approximately 1.5–4 ms across typical P3b amplitudes. Thus, the neural–cardiac association was specific to parietal P3b and to the heartbeats in which the stimulus-locked cardiac brake was expressed (Fig. 3, Table S11, Fig. S3). However, the µV measurement is dependent on the specific recording setup and should not be extrapolated directly to other settings. Rather, the estimates are provided to demonstrate the overall degree of effects in the current setup, in contrast to behavioural findings.

**Figure 3.**
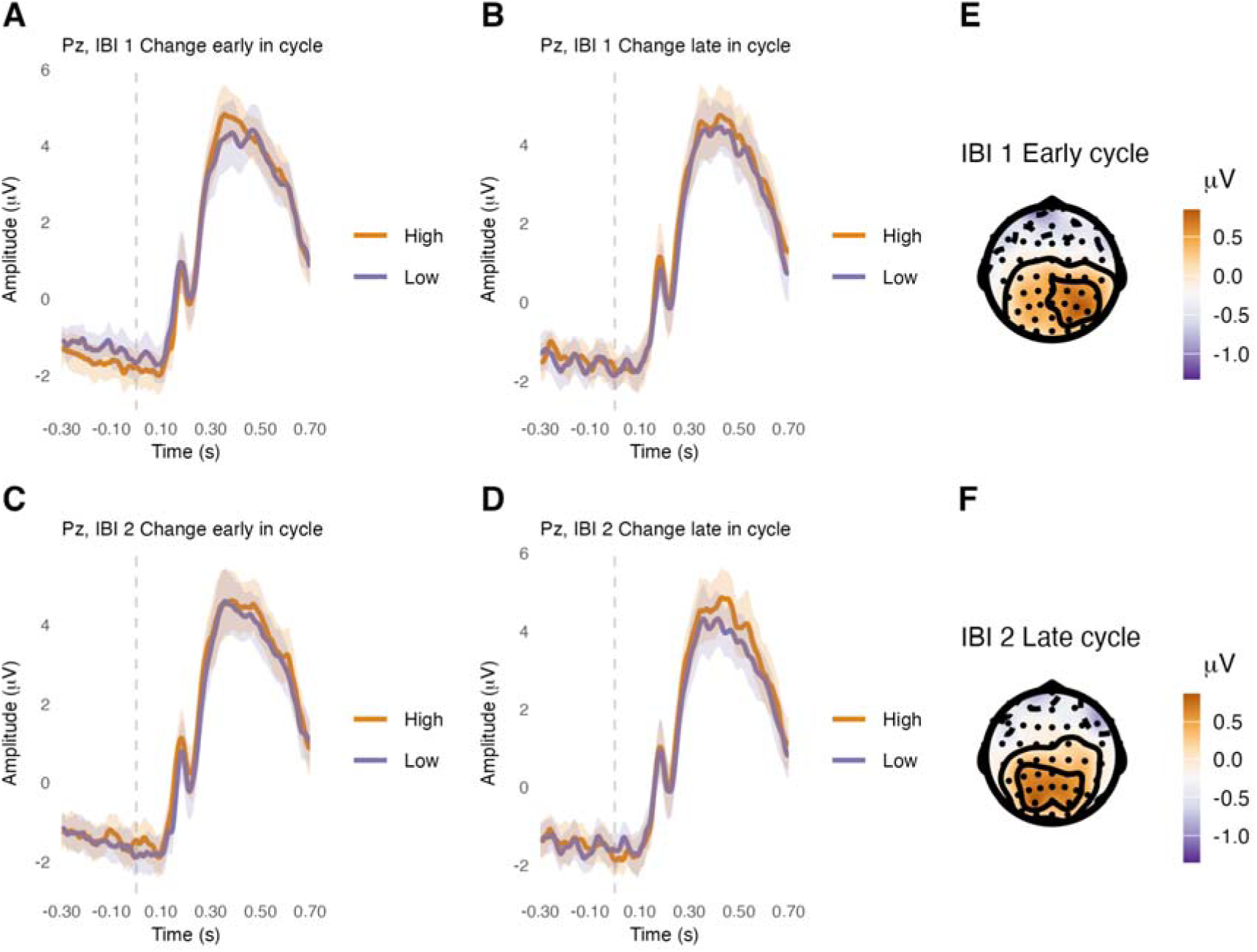
Pseudotrial-corrected parietal evoked responses associated with feedback-locked cardiac deceleration. (A–D) Grand-average feedback-evoked responses at Pz for trials with high versus low interbeat-interval (IBI) change, defined by a within-participant median split. Responses are shown for the first heartbeat after feedback, IBI 1, when feedback occurred early (A) or late (B) in the cardiac cycle, and for the second heartbeat after feedback, IBI 2, when feedback occurred early (C) or late (D) in the cardiac cycle. All participant-level averages were pseudotrial corrected before group averaging to reduce systematic differences caused by cardiac field artefacts associated with unequal R-peak timing across conditions. Lines show grand-average pseudotrial-corrected responses and shaded areas indicate 95% confidence intervals. The vertical dashed line marks feedback onset. (E, F) Scalp topographies of the pseudotrial-corrected difference between high- and low-IBI-change conditions for the cardiac-phase-specific heartbeats showing the expected stimulus-locked cardiac deceleration: IBI 1 when feedback occurred early in the cardiac cycle (E) and IBI 2 when feedback occurred late in the cardiac cycle (F). Topographies show the mean high-minus-low amplitude difference within the phase-specific parietal P300 50-millisecond time window centered on 360 (E) or 446 (F) milliseconds post-feedback. Time windows were determined based on group-level phase-specific grand average responses. Larger parietal P300 amplitudes were associated with stronger cardiac deceleration for IBI 1 in early-cycle trials and IBI 2 in late-cycle trials. Abbreviations: IBI, interbeat interval; µV, microvolts; s, seconds.

Posterior-predictive simulations indicated that the observed neural–cardiac associations were detectable under the specified model and sampling assumptions. For the P3b association with R1 in early cardiac-cycle trials, the dataset had 72% power to detect BF > 10 and 83% power to detect BF > 3. For the P3b association with R2 in late cardiac-cycle trials, power was 76% for BF > 10 and 92% for BF > 3. Thus, the neural–cardiac effects were detectable in the current sample despite the smaller number of trials available after splitting into early- or late-cardiac-cycle conditions and evoked response qualities that include heightened noise levels.

Larger P3b amplitudes therefore predict stronger deceleration at R1 in early cardiac-cycle trials and R2 in late cardiac-cycle trials. These are the cardiac-phase-specific heartbeats with the most strongly expressed effects of stimulus-locked parasympathetic brake. Cholinergic attention-related contributions within the multicomponent P300 response likely underlie the association, allowing better perception^19,42,45,46^. Additionally, we did not observe robust associations between prediction processing (signed and unsigned PE, outcome valence, expectation) and R2 in late cardiac-cycle trials or R1 modulations. These findings are consistent with the possibility that, even within the same heartbeat and especially across heartbeats, transient cardiac changes reflect diverse processes that include not only adjusting to external information and internal processing of it, but also, distinctly, orienting to a meaningful stimulus.

### Control analyses show physiological, computational, and neural specificity

We performed a broad set of control analyses to test whether the main effects could be explained by alternative physiological or behavioural variables or attributed to analysis settings. For the cardiac-prediction analyses, we controlled for baseline heart-rate variability (HRV) and average IBI during a 5-minute resting recording by quantifying HRV as the root mean square of successive differences (RMSSD). To contrast transient beat-by-beat dynamics with ongoing tonic cardiac states before and after feedback, we also assessed RMSSD and average IBI per trial. We used the trial number in a block (from 1 to 100) to control for learning effects. Moreover, the probabilistic task contained pre-feedback movement confounds in approach but not in avoid decision trials. We accounted for this distinction in three ways: by controlling for the decision type (approach/avoid), analysing them separately, and also controlling for reaction time in approach trials. To assess the sensitivity of statistical analyses, we reran the models with more or less informative priors, varied the HRV outlier removal, and compared the intercept-only models to the ones with random slopes. To show the advantages of the beat-to-beat analysis, we also reran the models on uncorrected IBI values, with or without controlling for the previous IBI length. For the cardiac-P3b analyses, the effects were additionally assessed against baseline activity (- 200 to 0 ms pre-feedback) and ECG amplitude in the same time window as P3b, as well as against feedback-related confounds (expectation, signed and unsigned PE, outcome).

The expectation, signed PE, outcome, and P3b effects across trials were robust to controlling for resting IBI and RMSSD, average IBI and RMSSD within the trial before and after feedback, trial number in the block, approach versus avoid decisions, prior sensitivity, HRV outlier removal, and presence of random slopes (Tables S12-26, S33-39, S48-51). Hence, these controls did not support an explanation in terms of baseline or trial-wise heart rate or HRV, learning or approach effects, as well as statistical setup. Rather, controlling for the ongoing heart rate and HRV revealed the specificity of beat-to-beat findings and even increased evidence in favour of some estimates. In the neural analyses, baseline, ECG amplitudes and feedback-related confounds did not explain the trial-wise P3b findings (Tables S31-32). Moreover, the analysis of pseudotrial-corrected averages between high and low IBI change conditions on the baseline time window at Pz, and at ECG channel showed specificity of P3b effects and effectiveness of pseudotrial correction (Tables S27- 30). The analysis of unadjusted IBI values, especially without accounting for the previous IBI, revealed that the beat-to-beat framework performs better in detecting the transient prediction-cardiac associations (Tables S40-47).

Separately analysing approach and avoid trials, where only the former contained a movement confound, led to generally decreased evidence in both conditions due to the lower number of available trials. Reaction-time controls further reduced evidence in approach-only analyses, which is expected given the shared variance among attention, response speed, and cardiac dynamics^47^. Expectation effects were affected the most but were not identified as negligible (BF in ROPE < 2.3). In avoid trials, where movement confounds were absent, there was still moderate to strong evidence that IBI changes scaled with expectation, signed and unsigned PE, outcome valence and P3b. This rules out a possibility that the effects merely reflect motor preparation or movement execution. Rather, the reaction speed seems to share variance with cardiac dynamics, especially with expectation-related effects. This observation is in line with the general view that prediction processing is an embodied process: cardiac and other bodily processes (e.g., reaction times, muscle tone, pupil dilation, skin conductance, etc.) are employed simultaneously to meet task demands. All of them are regulated by the autonomic nervous system, a shared overarching process.

## Discussion

We show that prediction processing is resolved in the timing of individual heartbeats. During probabilistic learning, interbeat intervals did not merely show a generic deceleration-acceleration response to feedback. Instead, distinct components of the prediction-processing loop were expressed in distinct heartbeats, evolving beat-by-beat. Model-derived expectation shaped anticipatory and rebound cardiac dynamics, signed prediction error was resolved in the second heartbeat after feedback, and stimulus-locked cardiac deceleration covaried with parietal P3b amplitude. The results therefore provide direct support for the view that autonomic physiology is continuously modulated during adaptive behaviour, rather than merely reflecting the salience or valence of events^6,20^.

The methodological advance is an interpretable, mechanistically grounded framework. Immediate changes in interbeat intervals unfold incrementally and are constrained by parasympathetic influences operating on a sub-second to second timescale^38^. By quantifying stimulus-locked cardiac responses as changes between consecutive interbeat intervals and accounting for cardiac phase^5^, we separated parasympathetically-mediated anticipatory activation, stimulus-locked brake, and later withdrawal. This made it possible to detect subtle cardiac effects that could be blurred by conventional heart-rate averaging or analysis of unadjusted interbeat intervals. The findings illustrate the precision gained by respecting physiological constraints on cardiac timing: prediction-related effects can be resolved in milliseconds in the moderately-sized sample of 34 young and healthy participants and in a task with simple symbolic feedback, even in the absence of salient reinforcement or threat.

Given the strength of effects, the present framework is a potentially useful and interpretable way to quantify how strongly bodily resources are employed during prediction processing. This could support future work on individual differences in brain-body coupling, including conditions in which autonomic regulation and adaptive behaviour are altered^28,33^. Rather than treating heart rate or its variability as a slow summary measure, beat-to-beat modelling can capture transient microstates of brain-body co-modulation during performance monitoring, as well as in other stimulus-locked contexts. The degree of parasympathetic response could then be studied in comparison to or complementary to conventional tonic cardiac parameters and macrostates. With further validation, it could present an effective intervention outcome: e.g., the clinical groups that overemploy bodily resources to threat or uncertainty could be trained to reduce the cardiac response. In contrast to neural evoked or oscillatory measures, it is possible to detect interbeat interval length online with little to no loss. The stimulus-evoked beat-to-beat framework may help distinguish which part of parasympathetic modulation - activation, brake, or withdrawal - requires intervention.

The study also clarifies the relationship between cardiac dynamics and neural feedback processing. Peak cardiac deceleration was coupled to parietal P3b amplitude in the cardiac-phase-specific heartbeats where the parasympathetic brake was most strongly expressed. This supports the idea that orienting to meaningful feedback is coordinated across neural and cardiac systems^19,42^. Moreover, we demonstrate that the processing and integration of stimulus features are reflected in later cardiac components: during acceleration, not at the moment of stimulus-locked deceleration.

Several limitations should be addressed in future studies. First, the present effects were observed in a relatively simple learning context, where feedback was symbolic rather than threatening or strongly incentivised. This makes the findings conservative. Stronger motivational contexts, such as monetary incentives, threat, or pain may elicit larger cardiac responses or alter their time course. The generality of the framework should now be tested across tasks that vary in salience, action demands, uncertainty, and clinical relevance. Second, here we focus on transient beat-to-beat cardiac adjustments that we interpret as vagally-mediated. However, we do not deny that prediction effects can also be related to slower fluctuations in sympathetic activity: e.g., average heart rate across several trials is related to learning effects and therefore likely to the average values of expectation and prediction error; or sympathetic activation after three seconds post-stimulus can show integration of evidence and preparation for future trials. In the future, it would make sense to compare different approaches to analysis and timescales of such co-modulations, but here we focus on the immediate cardiac response. Third, respiration was not recorded and should be included in future studies to separate respiratory frequency and phase from prediction-related cardiac dynamics. However, the observed effects were extremely short-lived, varied from heartbeat to heartbeat, and survived controls based on average interbeat interval and heart-rate variability, making a simple tonic cardiac or respiratory-rate explanation unlikely.

Together, these findings show that prediction processing is expressed in peripheral autonomic physiology with single-heartbeat precision. Thus, we provide direct evidence that heart-rate fluctuations reflect moment-to-moment predictive adjustments. Moreover, the beat-to-beat framework provides a way to estimate the precision and timescale with which bodily regulation accompanies adaptive behaviour. This opens methodological ground for studies of interindividual differences in brain-body coupling. Disturbed central-peripheral integration has been implicated in anxiety^14,31,34^, panic attacks^48^, attention-deficit disorders^49^, schizophrenia^50^, and ageing^51^, where bodily reactions to unexpected events may be up- or down-regulated. The present approach may therefore help quantify how individuals with differing degrees of central-peripheral integration modulate distinct components of the cardiac response in adaptive behaviour and performance monitoring.

## Methods

### Participants and ethics

The study was approved by the Ethics Committee of the Medical Faculty of Leipzig University (559/20-ek) and was conducted in accordance with the Declaration of Helsinki. Participants were recruited via the recruitment system of the Max Planck Institute for Human Cognitive and Brain Sciences, provided written informed consent before data collection, and received 9 euro per hour for participation. Volunteers had to be aged between 18 and 35 years and be fluent in German or English. 37 adults were recorded. Three were excluded after initial visual screening due to poor EEG data quality, leaving 34 participants (17 women; mean age = 26.2 years, SD = 3.5, range = 21–35 years). Exclusion criteria were neurological or psychiatric disorders, any occurrence of epileptic seizures, history of cardiac disorders, pregnancy or breastfeeding, and unwillingness to grant informed consent or be informed of incidental findings.

### Experimental procedure

After receiving instructions and signing a consent form, participants were fitted with EEG and ECG and seated in front of a computer in a soundproof, electrically shielded room. The session began with a five-minute eyes-open resting recording, which was used to derive participant-level measures of average IBI and resting HRV.

Participants then completed a probabilistic approach–avoidance learning task that lasted about 45-60 minutes. Stimuli were presented in MATLAB^52^ using Psychophysics Toolbox^53^ on a Sony CRT Trinitron Multiscan 200GS monitor (resolution 1280 x 1024). Eight visually neutral symbols from the Agathodaimon font were used: two during 10 practice trials and a different pair in each of three experimental blocks. All stimuli were 180x180 pixels in size and presented in white on a black background. Each experimental block contained 100 trials and was followed by a self-paced break.

Within each block, participants learned to approach or avoid one of two cues, by pressing a spacebar with the index finger of the right hand to approach or by withholding a response until a two-second response elapsed. Each trial began with a three-second fixation cross, followed by a one-second cue presentation and the two-second decision window. Feedback was then displayed for three seconds. Positive and negative outcomes were represented by green smiling and red sad emoji faces, respectively (Fig. 1). In the first 50 trials of each block, one cue resulted in a positive outcome 70% of the time when approached and 30% of the time when avoided. The other cue, when approached, resulted in a positive outcome 30% of the time, and when avoided, 70% of the time. After 50 trials, the contingencies associated with the two cues were symmetrically reversed without warning, requiring participants to update which cue should be approached and which should be avoided. This intentionally complex task structure resulted in, on average, 66% of correct choices and 55% of positive outcomes across trials and participants, generating a broad range of prediction errors and keeping the task engaging.

### Electrophysiological recording

Continuous EEG was recorded with a 64-channel cap (ant-neuro Wave-Guard), using a TMSi (EJ Oldenzaal, The Netherlands) DC-amplifier and a sampling rate of 500 Hz. We kept electrode impedances below 5 kOhm for all channels. The linked mastoid electrodes were used as a reference. An electrode placed on the sternum served as ground. Bipolar EOG channels were placed at the outer canthi and above and below the right eye. ECG was recorded from two bipolar electrodes placed on the left and right clavicles (lead I).

### Computational modelling

Trial-wise expectations and prediction errors were estimated by fitting three reinforcement-learning models separately to each participant^8,54–57^. Because the two cues had complementary contingencies, responses were represented as selection of one of two strategies: a “correct” strategy (approaching the advantageous cue or avoiding the disadvantageous cue) and the complementary “incorrect” strategy.

### Choice rule

LetQ_c_andQ_inc_be the learned Q-values associated with each of the strategies, ‘correct’ and ‘incorrect’, before trial . Both values were initially set to zero. The probability of selecting the strategy coded as correct was given by a softmax function:

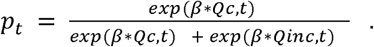

The inverse-temperature parameter quantified sensitivity to value differences (i.e., exploration/exploitation tradeoff). The Q-value of the selected strategy immediately before feedback was used as the trial-wise expectation estimate.

### Learning rules

Outcomesr_t_were coded as +1 and -1. Signed prediction error was calculated as the difference between the outcome and the selected strategy value:

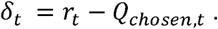

Unsigned prediction error was |δ_t_|. Three learning models were fitted. In the standard Rescorla–Wagner model, only the selected value was updated:

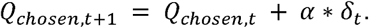

In the anticorrelated Rescorla–Wagner model, both strategy values were updated with a common learning rate:

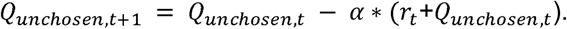

The confirmation-bias model used the same anticorrelated updating structure but estimated separate learning rates for positive and negative outcomes (α_pos_andα_neg_). The reinforcement-learning models were compared with a noisy win-stay-lose-shift model^57^ and a random-response model.

### Parameter estimation and model selection

Models were fitted in Python by maximum-likelihood estimation using L-BFGS-B optimisation with 25 stochastic starting values^58^. Inverse temperature was bounded between 0.01 and 10; learning-rate and the win-stay-lose-shift parameter were bounded between 0.01 and 0.99. Model fit was quantified as

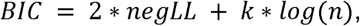

where negLL is the negative log-likelihood, k is the number of fitted parameters (2 or 3), and n is the number of trials. The reinforcement-learning models first had to outperform the heuristic controls in cumulative BIC. The best reinforcement-learning model was then identified for each participant, and the model selected for the largest number of participants was used to derive trial-wise variables for the complete sample. The confirmation-bias model was selected by this procedure because it explained best the data of 14 participants and also had the lowest cumulative BIC (Fig. S4).

### ECG processing and cardiac measures

#### R-peak detection

ECG data were processed in MATLAB^52^. Where necessary, the signal was inverted so that R-peak deflections were positive. Initial R-peak timings were detected using HEPLAB^59^ for EEGLAB^60^, which implements the Pan-Tompkins detection algorithm^61^. A custom code then moved an event to the largest local deflection when a different maximum occurred within a 20-millisecond window. Detected R peaks were inspected visually, with particular attention to the four cycles preceding and four cycles following feedback.

### Interbeat intervals and successive changes

For each trial, IBI were calculated for four cardiac cycles before feedback (R-4 to R-1) and four cycles after feedback (R1 to R4). R1 denotes the interval containing feedback onset. The primary cardiac measure was the difference between successive intervals:

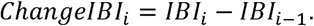

Positive values therefore indicate that the current interval was longer than the preceding interval (relative cardiac slowing), whereas negative values indicate relative shortening (cardiac acceleration). This differencing step was used to separate rapid incremental adjustments from the strong autocorrelation and slower tonic influences present in absolute IBI values^38^. The article focuses on R−1, R1, R2 and R3.

### Cardiac phase at feedback

Cardiac phase was expressed as the proportion of the ongoing R-1-to-R1 interval that had elapsed at feedback onset:

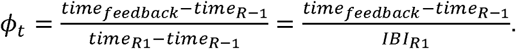

The continuous phase measure ranged from 0 immediately after R-1 to 1 immediately before R1 and was included in the prediction-related cardiac models because here we were looking to quantify how much of the cardiac cycle had elapsed before feedback could affect ongoing or subsequent interval. For categorical phase-specific analyses, trials in the first third of the cycle were classified as early-cycle trials and trials in the final third as late-cycle trials; the middle third was not included. These groups may be termed systole and diastole for comparability with prior work, but they do not constitute direct measurements of mechanical ventricular systole and diastole because T-wave offsets were not identified^62^.

### Tonic cardiac measures

Average IBI and the root mean square of successive differences (RMSSD) were calculated from the five-minute eyes-open resting recording. Average IBI and RMSSD were also computed separately from the four intervals before and four intervals after feedback within each trial. The short trial-level RMSSD estimates were used only as control variables contrasting global within-trial variability with heartbeat-specific IBI changes, not as standalone estimates of conventional HRV.

### EEG preprocessing and feedback-evoked responses

#### Preprocessing

EEG data were processed in MATLAB^52^ with EEGLAB^60^. Continuous data were high-pass filtered at 0.1 Hz and low-pass filtered at 40 Hz using bidirectionally applied zero-phase, non-causal second-order Butterworth IIR filters, yielding an effective fourth-order magnitude response (filtfilt function, -6dB at the nominal cutoff frequencies), followed by a 48–52-Hz notch filter (pop_eegfiltnew). Channels containing more than five seconds of flat signal or correlating with adjacent channels below 0.85 were marked as bad (clean_artifacts), reconstructed by spherical-spline interpolation (pop_interp). Then, the data was re-referenced to the common average. On average, 2.4 channels were interpolated per participant (max. 10), Pz was never among them.

Independent component analysis (ICA) was used to attenuate artefacts. FastICA was fitted to a copy of the recording additionally high-pass filtered at 1 Hz, with the rank set to the number of non-interpolated channels and recording breaks excluded. Artefactual components were identified by combining ICLabel^63^ and SASICA^64^. ICLabel probabilities above 0.90 for cardiac, ocular, muscular or “other” activity were considered evidence for artefact when the probability of brain activity was below 0.05. SASICA flagged components based on correlations with ECG exceeding two standard deviations from the mean, correlations with EOG exceeding four standard deviations, atypical autocorrelation exceeding two standard deviations or focal spatial concentration exceeding two standard deviations. Components identified by either procedure were removed only when their ICLabel probability of brain activity was below 0.05. On average, 9.8 components were identified as bad (max. 16), out of them, 0.53 were marked as cardiac (max. 2).

### Feedback-evoked responses

Feedback-locked epochs from -300 to 700 ms were generated using MNE-Python^65^. Epochs were flagged as artefactual when peak-to-peak activity at any channel exceeded 200 µV between 100 and 500 ms after feedback (6 epochs for one participant, two for another, one epoch for eight participants, and zero otherwise). Flagged epochs were removed when constructing participant-level evoked averages and visualisations. Trial-wise neural-cardiac models used single-trial mean amplitudes after the three-standard-deviation amplitude exclusion. Epochs were not baseline corrected because the -200-to-0-ms interval could contain anticipation-related activity and phase-dependent cardiac contamination; prestimulus amplitude was used as a control variable instead.

Parietal P3b was quantified at Pz. Participant-specific positive peaks were identified between 300 and 500 ms after feedback and then averaged to determine the group-level latency. Because peak latencies statistically differed between early- and late-cycle trials (paired t-test p-value < 0.1, and peaks are no closer than 25 ms), phase-specific 50-ms windows were used: 335–385 ms for early-cycle trials (centred at 360 ms) and 421–471 ms for late-cycle trials (centred at 446 ms). Frontal P3a was quantified at FCz from 353 to 403 ms (centred at 378 ms) and it did not differ for early- and late-cycle trials. Single-trial amplitudes were calculated as the mean voltage within the corresponding window.

### Pseudotrial correction of cardiac field artefacts

Sorting feedback-locked EEG by cardiac phase or IBI change also changes the distribution of R peaks around feedback and can therefore generate systematic cardiac field artefacts ^66–68^. We used pseudotrials to estimate this contamination^66,69^. For each participant, 300 artificial feedback triggers were positioned randomly within the task recording. Epochs from - 300 to 700 ms were generated around these triggers and classified using the real R-peak events and the same cardiac-phase and IBI-change definitions as the actual trials. Trigger sets that produced overlapping epochs were regenerated, with a maximum of 500 attempts. The procedure was repeated 10 times and the resulting pseudotrial averages were combined.

Actual trials were divided within participants into high- and low-IBI-change conditions by a median split, and the corresponding pseudotrial waveform was subtracted from each participant-level feedback-locked average. The full specificity screening covered R-1, R1, R2 and R3 in early- and late-cycle trials and the P3b, P3a, and prestimulus windows. Pseudotrial-corrected high-minus-low P3 differences were compared with zero using participant-level Bayesian models. Matching ECG-amplitude contrasts were examined to verify that residual differences were not attributable to cardiac activity, where ECG signal was processed in the same way as EEG. The median split was used only for participant-level corrected evoked responses; the trial-wise analyses treated P3b amplitude and IBI change as continuous variables and additionally controlled for ECG amplitude in the P3b time window or for the average EEG baseline amplitude.

### Statistical analysis

Analyses were performed in R^70^ using brms^71^ with the cmdstan^72^ backend. Trial-level models included a participant-specific random intercept. Random-slope versions were fitted as sensitivity analyses and compared with the random-intercept models using leave-one-out cross-validation. IBI-change outcomes were fitted with Student-t likelihoods after distributional diagnostics (Supplementary Information 6); the likelihood for contrast models was chosen from Gaussian, Student-t and skew-normal families using distributional analytics (Kolmogorov-Smirnov, Cramer-von-Mises, Shapiro-Wilk, Anderson-Darling tests; skewness and kurtosis values).

### Average cardiac dynamics

Mean IBI change at each cardiac cycle from R-3 through R4 was estimated with intercept-only multilevel models for all trials and separately for early- and late-cycle trials. IBI change remained in milliseconds for these descriptive models, and a narrow ROPE of -0.1 to 0.1 ms was used to distinguish slowing, acceleration and practical stability. The cardiac phase distribution (uniformity tests according to Kolmogorov-Smirnov, Cramer-von-Mises, skewness and kurtosis) and its stability over trial number (based on a mixed model with random participant-level intercepts) were assessed to exclude systematic appearance of feedback in a particular phase of the cardiac cycle.

### Prediction-related cardiac models

Separate models related IBI changes to prediction features: expectation, outcome valence, signed and unsigned PE. The models were:

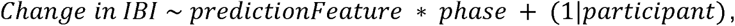

Expectation and PE were continuous variables, outcome valence was entered as a factor contrasting negative to positive feedback. Identical models were evaluated across R-3 to R4 to establish temporal specificity, with main focus on R-1 to R3. Phase-specific follow-up models were fitted separately to early- and late-cycle trials when the continuous phase interaction was supported and were not a part of main inference.

### Neural-cardiac models

First, participant-level pseudotrial-corrected contrasts were analysed separately with one-sample Bayesian models. The trial-wise models tested whether P3b or P3a amplitude predicted R1 IBI change in early-cycle trials and R2 IBI change in late-cycle trials:

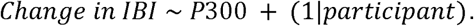

Equivalent component- and heartbeat-specific analyses involving the remaining cardiac cycles were used to evaluate specificity.

### Data transformations

Trial-wise outliers were defined as values at least three standard deviations from the pooled mean across participants and trials. A common exclusion mask was constructed from IBI changes at R-1, R1, R2 and R3 so that a trial excluded at one of these cycles was excluded consistently from the principal cardiac models. Neural-cardiac analyses additionally excluded extreme single-trial amplitudes. Continuous variables were standardised after outlier removal. Coefficients were subsequently transformed back to original units and reported as milliseconds of IBI change per predictor unit, as the negative-versus-positive outcome contrast, or as milliseconds per microvolt of P3 amplitude.

### Bayesian estimation and evidence

Models used four Markov chains with 4,000 iterations per chain, including 2,000 warm-up iterations, and Hamiltonian Monte Carlo sampling with *adapt_delta* = 0.95. For IBI change models, fixed-effect coefficients received Normal(0, 0.05) priors. Sensitivity analyses used Normal(0, 0.03) and Normal(0, 0.10) priors. In other models, Normal(0, X) for intercept was used, where X is the standard deviation of the dependent variable multiplied by 1.5. For every coefficient, we report the posterior density mode and 95% highest-density interval (HDI). Convergence was evaluated using R-hat < 1.01, absence of divergent transitions and posterior predictive checks, following Bayesian reporting recommendations^73^.

Evidence for a non-zero coefficient was quantified with Bayes factors obtained with bayestestR^74^ (describe_posterior) by exponentiating the coefficient-level log evidence ratio. BF > 10 was treated as strong evidence in the main analyses; BF > 3 was used as a moderate-evidence threshold.

Regions of practical equivalence (ROPE) were defined in original units so that a predictor spanning its observed range would have to change IBI by at least 1 ms to be considered practically meaningful. For a standardised continuous predictor, the upper ROPE bound was:

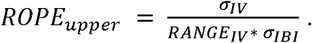

For outcome valence, the 1-ms criterion applied directly to the negative-versus-positive contrast. Evidence that an effect was practically negligible was quantified as the Bayes factor for the coefficient lying inside the ROPE.

A Bayesian FDR correction based on posterior ROPE overlap was applied across the families of heartbeat-wise (R-1 to R3) and component-wise (prediction features, neural) tests; we report only the comparisons that survive this correction. Coefficients were ranked from the smallest to the largest ROPE proportion, and the cumulative mean local FDR was calculated at each rank. The largest set of coefficients for which this cumulative value remained at or below (q = 0.05) was classified as discoveries.

### Control and sensitivity analyses

The principal cardiac models were rerun with the following controls added individually: resting average IBI and RMSSD; pre-feedback and post-feedback trial-level average IBI and RMSSD; trial number within block; approach versus avoid decision; and reaction time within approach trials. Approach and avoid trials were also analysed separately, providing an avoid-only analysis without a pre-feedback motor response. Additional sensitivity analyses varied the coefficient prior (Normal(0, 0.03) or Normal(0, 0.10)), excluded two resting RMSSD outliers, fitted participant-specific random slopes, and replaced successive IBI change with absolute IBI either with or without adjustment for the immediately preceding interval. For P3b models, further controls included expectation, outcome valence, signed and unsigned prediction error, mean Pz amplitude from -200 to 0 ms, and mean ECG amplitude in the P3b window.

### Posterior-predictive evidential power

Evidential power was estimated with 200 simulations of each fitted model. In each simulation, data were generated from one posterior draw (posterior_predict) and analysed with the same model, prior, sampling settings and Bayes-factor threshold. Simulations with R-hat > 1.01 or divergent transitions for any tested fixed effect were excluded using global gating. Power was the proportion of retained simulations in which the relevant coefficient exceeded BF > 3 or BF > 10. Simulations were conducted for the observed 34-participant sample and for a possible larger sample of 100 participants.

## Data availability statement

The raw data, model fits, and other data derivations, are available at https://osf.io/3ghyw/.

## Code availability statement

All codes used for analysis and visualisation are openly available at https://github.com/mariaazanova/HRPE.

## Acknowledgements

This project is supported by the Max Planck Society.

## Author contributions

M.A.: Conceptualization, Methodology, Software, Formal analysis, Investigation, Data Curation, Writing – original draft, Writing – review & editing, Visualization, Project administration. L.S.: Conceptualization, Investigation, Data Curation, Writing – review & editing. A.S.: Code Review, Writing – review & editing. E.A.: Data Curation. V.V.N.: Writing – review & editing, Supervision. A.V.: Writing – review & editing, Supervision, Funding acquisition.

