## Supplementary Information for "Individual heartbeats track distinct prediction processes during human probabilistic learning"

### Contents

1. Cardiac dynamics surrounding feedback	3

2. Prediction features and interbeat intervals	4

2.1. Expectation	4

2.2. Signed prediction error	6

2.3. Outcome valence	9

2.4. Unsigned prediction error	11

3. Neural responses and interbeat intervals	13

4. Control and sensitivity analyses	14

4.1. Expectation	14

4.1.1. Average IBI or RMSSD during rest	14

4.1.2. Average IBI or RMSSD within a trial, before or after feedback	16

4.1.3. Number of a trial in a block and approach/avoid decisions	18

4.1.4. Reaction times	20

4.1.5. Prior sensitivity	21

4.1.6. HRV outlier removal	22

4.2. Signed PE and outcome valence	23

4.2.1. Average IBI or RMSSD during rest	23

4.2.2. Average IBI or RMSSD within a trial, before or after feedback	25

4.2.3. Number of a trial in a block and approach/avoid decisions	28

4.2.4. Reaction times	29

4.2.5. Prior sensitivity	31

4.2.6. HRV outlier removal	32

4.3. P3b analyses	33

4.3.1. Pseudotrial-corrected average responses	33

4.3.2. Control for features of feedback	36

4.3.3. Baseline at Pz and average ECG amplitude	37

4.3.4. Average IBI or RMSSD during rest	38

4.3.5. Average IBI or RMSSD within a trial, before or after feedback	39

4.3.6. Number of a trial in a block and approach/avoid decisions	40

4.3.7. Reaction times	41

4.3.8. Prior sensitivity	42

4.3.9. HRV outlier removal	42

4.4. Results based on uncorrected IBI values	43

4.4.1. Controlling for previous IBI	43

4.4.2. Without controlling for previous IBI	51

4.5. Models with and without random slopes	57

4.5.1. Model results with random slopes	57

4.5.2. Model comparisons	60

5. Computational modelling results	61

5.1. Model fit	61

5.2. Recovery and confusion	62

6. Distributional diagnostics	63

### 1. Cardiac dynamics surrounding feedback

*Table S1.* The degree of change in milliseconds of the current interbeat interval (IBI) compared to the previous one, for three heartbeats before and four heartbeats after feedback. ROPE – region of practical equivalence, BF – Bayes Factor. Note that RHat for the model with the change of IBI 3 is >1.01, indicating mild MCMC convergence issues.

| Parameter | Dependent variable: Change of IBI -3 | Dependent variable: Change of IBI -2 | Dependent variable: Change of IBI -1 | Dependent variable: Change of IBI 1 | Dependent variable: Change of IBI 2 | Dependent variable: Change of IBI 3 | Dependent variable: Change of IBI 4 |
| --- | --- | --- | --- | --- | --- | --- | --- |
| Intercept | Mode: 11.681 95% CI: [8.848,13.922] BF: 59441 ROPE Bound: 0.1 In ROPE: 0 BF in ROPE: 0 RHat: 1.003 | Mode: 11.78 95% CI: [8.705,13.715] BF: 45306 ROPE Bound: 0.1 In ROPE: 0 BF in ROPE: 0 RHat: 1.002 | Mode: 1.89 95% CI: [0.937,6.892] BF: 0.465 ROPE Bound: 0.1 In ROPE: 0 BF in ROPE: 2.084 RHat: 1.005 | Mode: 1.558 95% CI: [0.655,6.63] BF: 0.293 ROPE Bound: 0.1 In ROPE: 0 BF in ROPE: 3.498 RHat: 1.002 | Mode: -1.269 95% CI: [-2.629,3.561] BF: 0.02 ROPE Bound: 0.1 In ROPE: 0.053 BF in ROPE: 53.405 RHat: 1.008 | Mode: -13.267 95% CI: [-19.038,-9.741] BF: 912 ROPE Bound: 0.1 In ROPE: 0 BF in ROPE: 0.001 RHat: 1.013 | Mode: -10.22 95% CI: [-15.927,-6.528] BF: 150 ROPE Bound: 0.1 In ROPE: 0 BF in ROPE: 0.007 RHat: 1.004 |
|  | Family: student Num. obs: 9808 Num. groups: 34 Bayes R2: 0.004 | Family: student Num. obs: 9808 Num. groups: 34 Bayes R2: 0.007 | Family: student Num. obs: 9808 Num. groups: 34 Bayes R2: 0.019 | Family: student Num. obs: 9808 Num. groups: 34 Bayes R2: 0.02 | Family: student Num. obs: 9808 Num. groups: 34 Bayes R2: 0.02 | Family: student Num. obs: 9808 Num. groups: 34 Bayes R2: 0.053 | Family: student Num. obs: 9808 Num. groups: 34 Bayes R2: 0.035 |

*Table S2*. The degree of change in milliseconds of the current interbeat interval (IBI) compared to the previous one only when the feedback was given in systole (first 1/3 of the cardiac cycle), for three heartbeats before and four heartbeats after feedback. ROPE – region of practical equivalence, BF – Bayes Factor.

| Parameter | Dependent variable: Change of IBI -3 | Dependent variable: Change of IBI -2 | Dependent variable: Change of IBI -1 | Dependent variable: Change of IBI 1 | Dependent variable: Change of IBI 2 | Dependent variable: Change of IBI 3 | Dependent variable: Change of IBI 4 |
| --- | --- | --- | --- | --- | --- | --- | --- |
| Intercept | Mode: 10.573 95% CI: [8.797,14.288] BF: 339750 ROPE Bound: 0.1 In ROPE: 0 BF in ROPE: 0 RHat: 1.001 | Mode: 10.92 95% CI: [6.451,12.491] BF: 785 ROPE Bound: 0.1 In ROPE: 0 BF in ROPE: 0.001 RHat: 1.001 | Mode: 1.117 95% CI: [-2.379,3.682] BF: 0.02 ROPE Bound: 0.1 In ROPE: 0.051 BF in ROPE: 53.51 RHat: 1 | Mode: 10.601 95% CI: [6.102,13.048] BF: 1760 ROPE Bound: 0.1 In ROPE: 0 BF in ROPE: 0.001 RHat: 1.001 | Mode: -10.872 95% CI: [-13.457,-3.728] BF: 5.414 ROPE Bound: 0.1 In ROPE: 0 BF in ROPE: 0.187 RHat: 1.002 | Mode: -13.017 95% CI: [-18.889,-10.028] BF: 4096 ROPE Bound: 0.1 In ROPE: 0 BF in ROPE: 0 RHat: 1.004 | Mode: -11.712 95% CI: [-14.379,-4.326] BF: 9.172 ROPE Bound: 0.1 In ROPE: 0 BF in ROPE: 0.11 RHat: 1.004 |
|  | Family: student Num. obs: 3260 Num. groups: 34 Bayes R2: 0.008 | Family: student Num. obs: 3260 Num. groups: 34 Bayes R2: 0.009 | Family: student Num. obs: 3260 Num. groups: 34 Bayes R2: 0.016 | Family: student Num. obs: 3260 Num. groups: 34 Bayes R2: 0.023 | Family: student Num. obs: 3260 Num. groups: 34 Bayes R2: 0.052 | Family: student Num. obs: 3260 Num. groups: 34 Bayes R2: 0.054 | Family: student Num. obs: 3260 Num. groups: 34 Bayes R2: 0.042 |

*Table S3*. The degree of change in milliseconds of the current interbeat interval (IBI) compared to the previous one only when the feedback was given in diastole (last 1/3 of the cardiac cycle), for three heartbeats before and four heartbeats after feedback. ROPE – region of practical equivalence, BF – Bayes Factor.

| Parameter | Dependent variable: Change of IBI -3 | Dependent variable: Change of IBI -2 | Dependent variable: Change of IBI -1 | Dependent variable: Change of IBI 1 | Dependent variable: Change of IBI 2 | Dependent variable: Change of IBI 3 | Dependent variable: Change of IBI 4 |
| --- | --- | --- | --- | --- | --- | --- | --- |
| Intercept | Mode: 11.914 95% CI: [8.243,13.881] BF: 30333 ROPE Bound: 0.1 In ROPE: 0 BF in ROPE: 0 RHat: 1.002 | Mode: 12.551 95% CI: [9.829,14.585] BF: 99460441 ROPE Bound: 0.1 In ROPE: 0 BF in ROPE: 0 RHat: 1.001 | Mode: 5.321 95% CI: [3.02,9.084] BF: 12.406 ROPE Bound: 0.1 In ROPE: 0 BF in ROPE: 0.082 RHat: 1.001 | Mode: -3.16 95% CI: [-4.645,2.912] BF: 0.025 ROPE Bound: 0.1 In ROPE: 0.036 BF in ROPE: 42.024 RHat: 1.004 | Mode: 8.748 95% CI: [5.027,10.045] BF: 713 ROPE Bound: 0.1 In ROPE: 0 BF in ROPE: 0.001 RHat: 1.001 | Mode: -14.526 95% CI: [-18.253,-8.267] BF: 2989 ROPE Bound: 0.1 In ROPE: 0 BF in ROPE: 0 RHat: 1.004 | Mode: -11.878 95% CI: [-17.265,-7.768] BF: 384 ROPE Bound: 0.1 In ROPE: 0 BF in ROPE: 0.003 RHat: 1.003 |
|  | Family: student Num. obs: 3265 Num. groups: 34 Bayes R2: 0.002 | Family: student Num. obs: 3265 Num. groups: 34 Bayes R2: 0.003 | Family: student Num. obs: 3265 Num. groups: 34 Bayes R2: 0.016 | Family: student Num. obs: 3265 Num. groups: 34 Bayes R2: 0.03 | Family: student Num. obs: 3265 Num. groups: 34 Bayes R2: 0.011 | Family: student Num. obs: 3265 Num. groups: 34 Bayes R2: 0.055 | Family: student Num. obs: 3265 Num. groups: 34 Bayes R2: 0.036 |

### 2. Prediction features and interbeat intervals

#### 2.1. Expectation

*Table S4.* The single-trial associations between the continuous measure of expectation and the degree of change of the current interbeat interval (IBI) compared to the previous one, for the three heartbeats before and the four heartbeats after feedback. ROPE – region of practical equivalence, BF – Bayes Factor. Cardiac phase is a continuous measure of distance from the last heartbeat before feedback to feedback, scaled by the distance from the last heartbeat before to the first heartbeat after feedback. All values are standardised.

| Parameter | Dependent variable: Change of IBI -3 | Dependent variable: Change of IBI -2 | Dependent variable: Change of IBI -1 | Dependent variable: Change of IBI 1 | Dependent variable: Change of IBI 2 | Dependent variable: Change of IBI 3 | Dependent variable: Change of IBI 4 |
| --- | --- | --- | --- | --- | --- | --- | --- |
| Intercept | Mode: -0.01 95% CI: [-0.037,0.014] BF: 0.007 ROPE Bound: 0.009 In ROPE: 0.402 BF in ROPE: 210 RHat: 1.006 | Mode: -0.029 95% CI: [-0.061,0.005] BF: 0.024 ROPE Bound: 0.013 In ROPE: 0.172 BF in ROPE: 55.46 RHat: 1.004 | Mode: 0.03 95% CI: [-0.044,0.067] BF: 0.011 ROPE Bound: 0.018 In ROPE: 0.508 BF in ROPE: 172 RHat: 1.003 | Mode: -0.019 95% CI: [-0.064,0.045] BF: 0.011 ROPE Bound: 0.018 In ROPE: 0.476 BF in ROPE: 157 RHat: 1.002 | Mode: 0.037 95% CI: [-0.027,0.084] BF: 0.018 ROPE Bound: 0.019 In ROPE: 0.331 BF in ROPE: 89.471 RHat: 1.003 | Mode: 0.016 95% CI: [-0.056,0.127] BF: 0.021 ROPE Bound: 0.02 In ROPE: 0.269 BF in ROPE: 60.991 RHat: 1.006 | Mode: 0.014 95% CI: [-0.054,0.101] BF: 0.016 ROPE Bound: 0.017 In ROPE: 0.305 BF in ROPE: 80.882 RHat: 1.004 |
| Expectation | Mode: 0.005 95% CI: [-0.006,0.01] BF: 0.092 ROPE Bound: 0.003 In ROPE: 0.497 BF in ROPE: 18.651 RHat: 1 | Mode: -0.002 95% CI: [-0.013,0.01] BF: 0.12 ROPE Bound: 0.004 In ROPE: 0.508 BF in ROPE: 14.165 RHat: 1 | Mode: -0.038 95% CI: [-0.048,-0.015] BF: 66.961 ROPE Bound: 0.006 In ROPE: 0 BF in ROPE: 0.023 RHat: 1 | Mode: -0.023 95% CI: [-0.037,-0.004] BF: 3.827 ROPE Bound: 0.005 In ROPE: 0.012 BF in ROPE: 0.4 RHat: 1 | Mode: 0.013 95% CI: [-0.011,0.021] BF: 0.198 ROPE Bound: 0.006 In ROPE: 0.438 BF in ROPE: 7.796 RHat: 1.001 | Mode: 0.024 95% CI: [0.017,0.051] BF: 108 ROPE Bound: 0.006 In ROPE: 0 BF in ROPE: 0.012 RHat: 1 | Mode: 0.011 95% CI: [-0.001,0.028] BF: 0.657 ROPE Bound: 0.005 In ROPE: 0.136 BF in ROPE: 2.005 RHat: 1.001 |
| Cardiac phase | Mode: -0.004 95% CI: [-0.011,0.004] BF: 0.121 ROPE Bound: 0.003 In ROPE: 0.369 BF in ROPE: 12.409 RHat: 1 | Mode: 0.012 95% CI: [0.004,0.025] BF: 3.708 ROPE Bound: 0.004 In ROPE: 0 BF in ROPE: 0.397 RHat: 1 | Mode: 0.029 95% CI: [0.024,0.056] BF: 844 ROPE Bound: 0.005 In ROPE: 0 BF in ROPE: 0.002 RHat: 1 | Mode: -0.074 95% CI: [-0.09,-0.059] BF: 459092653 ROPE Bound: 0.005 In ROPE: 0 BF in ROPE: 0 RHat: 1.001 | Mode: 0.116 95% CI: [0.099,0.131] BF: 34661997152952 ROPE Bound: 0.005 In ROPE: 0 BF in ROPE: 0 RHat: 1.001 | Mode: 0.016 95% CI: [-0.009,0.024] BF: 0.251 ROPE Bound: 0.006 In ROPE: 0.365 BF in ROPE: 5.595 RHat: 1.002 | Mode: -0.02 95% CI: [-0.028,0] BF: 1.227 ROPE Bound: 0.005 In ROPE: 0.076 BF in ROPE: 1.173 RHat: 1 |
| Expectation : Cardiac phase | Mode: 0.004 95% CI: [-0.004,0.012] BF: 0.148 ROPE Bound: 0.001 In ROPE: 0.092 BF in ROPE: 7.558 RHat: 1 | Mode: -0.006 95% CI: [-0.01,0.011] BF: 0.11 ROPE Bound: 0.001 In ROPE: 0.157 BF in ROPE: 10.442 RHat: 1 | Mode: -0.014 95% CI: [-0.019,0.013] BF: 0.17 ROPE Bound: 0.002 In ROPE: 0.151 BF in ROPE: 6.342 RHat: 1 | Mode: -0.013 95% CI: [-0.02,0.01] BF: 0.205 ROPE Bound: 0.002 In ROPE: 0.135 BF in ROPE: 5.476 RHat: 1 | Mode: -0.021 95% CI: [-0.025,0.006] BF: 0.315 ROPE Bound: 0.002 In ROPE: 0.081 BF in ROPE: 3.401 RHat: 1 | Mode: 0.005 95% CI: [-0.019,0.015] BF: 0.18 ROPE Bound: 0.002 In ROPE: 0.161 BF in ROPE: 6.622 RHat: 1.001 | Mode: 0.011 95% CI: [-0.012,0.016] BF: 0.155 ROPE Bound: 0.002 In ROPE: 0.158 BF in ROPE: 7.392 RHat: 1.002 |
|  | Family: student Num. obs: 9808 Num. groups: 34 Bayes R2: 0.004 | Family: student Num. obs: 9808 Num. groups: 34 Bayes R2: 0.007 | Family: student Num. obs: 9808 Num. groups: 34 Bayes R2: 0.022 | Family: student Num. obs: 9808 Num. groups: 34 Bayes R2: 0.026 | Family: student Num. obs: 9808 Num. groups: 34 Bayes R2: 0.034 | Family: student Num. obs: 9808 Num. groups: 34 Bayes R2: 0.054 | Family: student Num. obs: 9808 Num. groups: 34 Bayes R2: 0.035 |

*Figure S1*. Priors (light grey) and posteriors (dark grey) from the models linking single-trial RR interval changes to model-derived expectations, corresponding to *Hypothesis 1*. (A) For the last heartbeat before feedback, R-1. (B) For the third heartbeat after feedback, R3. The dashed lines mark the respective ROPE (region of practical equivalence). BF is Bayes Factor, here it is an evidence ratio for coefficients being outside of ROPE.

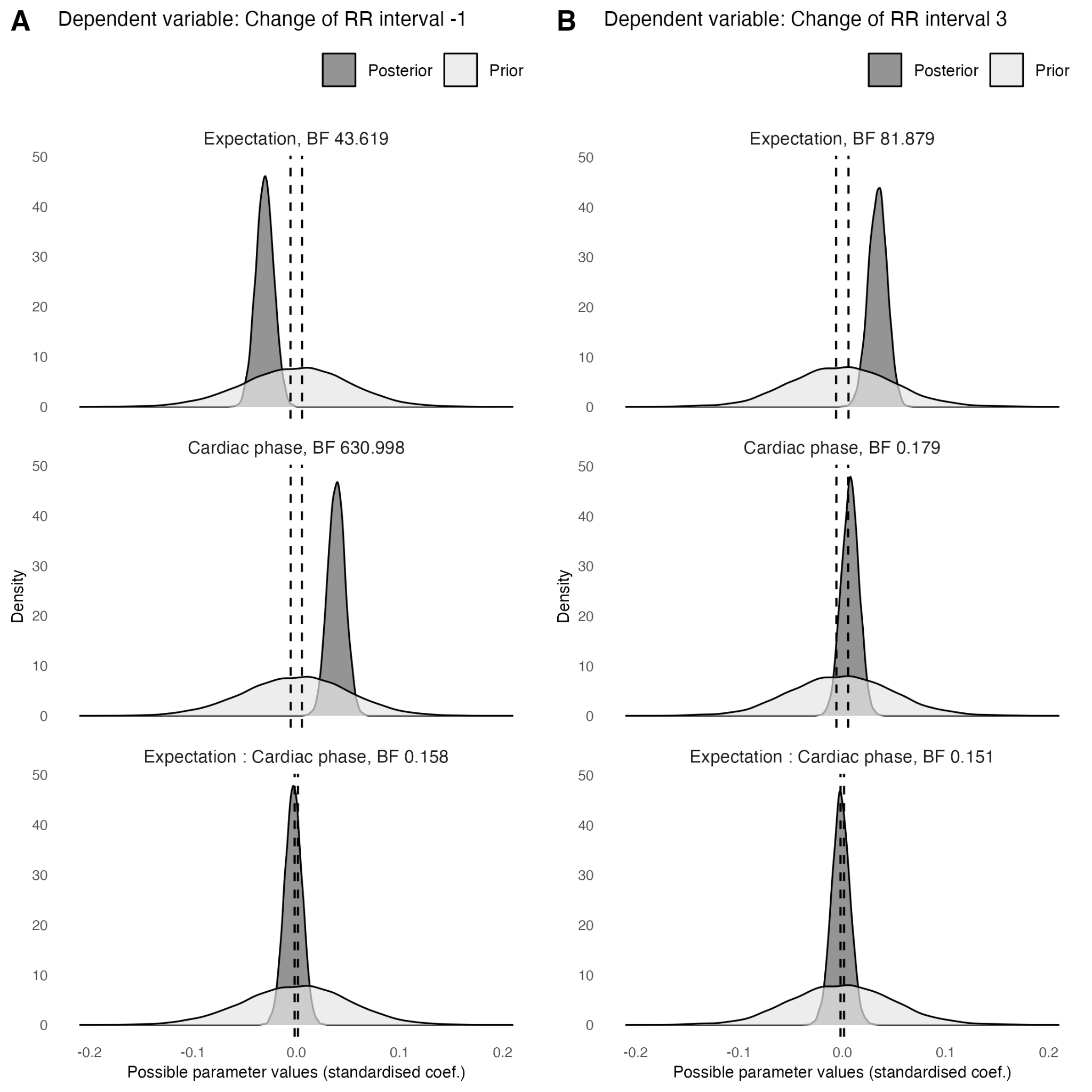

#### 2.2. Signed prediction error

*Table S5*. The single-trial associations between the continuous measure of prediction error (PE) and the degree of change of the current interbeat interval (IBI) compared to the previous one, for three heartbeats before and four heartbeats after feedback. ROPE – region of practical equivalence, BF – Bayes Factor. Cardiac phase is a continuous measure of distance from the last heartbeat before feedback to feedback, scaled by the distance from the last heartbeat before to the first heartbeat after feedback. All values are standardised.

| Parameter | Dependent variable: Change of IBI -3 | Dependent variable: Change of IBI -2 | Dependent variable: Change of IBI -1 | Dependent variable: Change of IBI 1 | Dependent variable: Change of IBI 2 | Dependent variable: Change of IBI 3 | Dependent variable: Change of IBI 4 |
| --- | --- | --- | --- | --- | --- | --- | --- |
| Intercept | Mode: -0.017 95% CI: [-0.035,0.014] BF: 0.007 ROPE Bound: 0.009 In ROPE: 0.42 BF in ROPE: 248 RHat: 1.003 | Mode: -0.039 95% CI: [-0.059,0.005] BF: 0.024 ROPE Bound: 0.013 In ROPE: 0.172 BF in ROPE: 59.515 RHat: 1.001 | Mode: 0.013 95% CI: [-0.05,0.061] BF: 0.01 ROPE Bound: 0.018 In ROPE: 0.51 BF in ROPE: 174 RHat: 1.001 | Mode: -0.026 95% CI: [-0.062,0.047] BF: 0.01 ROPE Bound: 0.018 In ROPE: 0.486 BF in ROPE: 159 RHat: 1.008 | Mode: 0.054 95% CI: [-0.032,0.085] BF: 0.018 ROPE Bound: 0.019 In ROPE: 0.322 BF in ROPE: 85.812 RHat: 1.003 | Mode: 0.132 95% CI: [-0.055,0.117] BF: 0.021 ROPE Bound: 0.02 In ROPE: 0.275 BF in ROPE: 63.556 RHat: 1.005 | Mode: 0.035 95% CI: [-0.049,0.101] BF: 0.015 ROPE Bound: 0.017 In ROPE: 0.329 BF in ROPE: 87.118 RHat: 1.005 |
| PE | Mode: -0.005 95% CI: [-0.011,0.005] BF: 0.103 ROPE Bound: 0.002 In ROPE: 0.381 BF in ROPE: 14.441 RHat: 1 | Mode: 0.01 95% CI: [-0.007,0.015] BF: 0.16 ROPE Bound: 0.003 In ROPE: 0.349 BF in ROPE: 8.845 RHat: 1 | Mode: 0.003 95% CI: [-0.008,0.025] BF: 0.271 ROPE Bound: 0.005 In ROPE: 0.284 BF in ROPE: 4.553 RHat: 1 | Mode: 0.013 95% CI: [0,0.031] BF: 1.045 ROPE Bound: 0.005 In ROPE: 0.062 BF in ROPE: 1.026 RHat: 1 | Mode: -0.024 95% CI: [-0.04,-0.009] BF: 16.369 ROPE Bound: 0.005 In ROPE: 0 BF in ROPE: 0.087 RHat: 1 | Mode: -0.012 95% CI: [-0.037,-0.003] BF: 2.59 ROPE Bound: 0.005 In ROPE: 0.019 BF in ROPE: 0.49 RHat: 1 | Mode: 0.017 95% CI: [0.001,0.029] BF: 1.315 ROPE Bound: 0.004 In ROPE: 0.049 BF in ROPE: 0.926 RHat: 1.001 |
| Cardiac phase | Mode: -0.003 95% CI: [-0.011,0.005] BF: 0.125 ROPE Bound: 0.003 In ROPE: 0.374 BF in ROPE: 12.236 RHat: 1.001 | Mode: 0.021 95% CI: [0.004,0.026] BF: 3.227 ROPE Bound: 0.004 In ROPE: 0.001 BF in ROPE: 0.425 RHat: 1 | Mode: 0.032 95% CI: [0.022,0.054] BF: 1615 ROPE Bound: 0.005 In ROPE: 0 BF in ROPE: 0.001 RHat: 1.001 | Mode: -0.071 95% CI: [-0.09,-0.059] BF: 15921953 ROPE Bound: 0.005 In ROPE: 0 BF in ROPE: 0 RHat: 1.002 | Mode: 0.113 95% CI: [0.099,0.13] BF: 2834367433731490 ROPE Bound: 0.005 In ROPE: 0 BF in ROPE: 0 RHat: 1 | Mode: 0.013 95% CI: [-0.008,0.025] BF: 0.26 ROPE Bound: 0.006 In ROPE: 0.37 BF in ROPE: 5.323 RHat: 1.002 | Mode: -0.019 95% CI: [-0.027,-0.001] BF: 1.353 ROPE Bound: 0.005 In ROPE: 0.063 BF in ROPE: 1.05 RHat: 1 |
| PE : Cardiac phase | Mode: -0.003 95% CI: [-0.013,0.003] BF: 0.168 ROPE Bound: 0.001 In ROPE: 0.072 BF in ROPE: 6.295 RHat: 1 | Mode: -0.002 95% CI: [-0.01,0.012] BF: 0.108 ROPE Bound: 0.001 In ROPE: 0.151 BF in ROPE: 10.256 RHat: 1 | Mode: 0.015 95% CI: [-0.008,0.025] BF: 0.244 ROPE Bound: 0.001 In ROPE: 0.085 BF in ROPE: 4.31 RHat: 1.001 | Mode: 0.012 95% CI: [-0.018,0.012] BF: 0.176 ROPE Bound: 0.001 In ROPE: 0.134 BF in ROPE: 6.51 RHat: 1.002 | Mode: 0.03 95% CI: [0.009,0.041] BF: 20.159 ROPE Bound: 0.001 In ROPE: 0 BF in ROPE: 0.051 RHat: 1 | Mode: -0.011 95% CI: [-0.03,0.003] BF: 0.567 ROPE Bound: 0.002 In ROPE: 0.046 BF in ROPE: 1.795 RHat: 1.001 | Mode: 0.017 95% CI: [-0.005,0.022] BF: 0.295 ROPE Bound: 0.001 In ROPE: 0.076 BF in ROPE: 3.48 RHat: 1 |
|  | Family: student Num. obs: 9808 Num. groups: 34 Bayes R2: 0.004 | Family: student Num. obs: 9808 Num. groups: 34 Bayes R2: 0.007 | Family: student Num. obs: 9808 Num. groups: 34 Bayes R2: 0.021 | Family: student Num. obs: 9808 Num. groups: 34 Bayes R2: 0.026 | Family: student Num. obs: 9808 Num. groups: 34 Bayes R2: 0.035 | Family: student Num. obs: 9808 Num. groups: 34 Bayes R2: 0.054 | Family: student Num. obs: 9808 Num. groups: 34 Bayes R2: 0.035 |

*Table S6*. Separately only within systole and diastole trials (first and last 1/3 of the cardiac cycle), the single-trial associations between the continuous measure of prediction error (PE) and the degree of change of the current interbeat interval (IBI) compared to the previous one, for the second and third heartbeats after feedback. ROPE – region of practical equivalence, BF – Bayes Factor. All values are standardised.

| Parameter | Dependent variable: Change of IBI 2 only in systole | Dependent variable: Change of IBI 2 only in diastole |
| --- | --- | --- |
| Intercept | Mode: -0.124 95% CI: [-0.234,-0.048] BF: 1.326 ROPE Bound: 0.019 In ROPE: 0 BF in ROPE: 0.892 RHat: 1.007 | Mode: 0.145 95% CI: [0.113,0.206] BF: 1558 ROPE Bound: 0.019 In ROPE: 0 BF in ROPE: 0.001 RHat: 1 |
| PE | Mode: -0.046 95% CI: [-0.078,-0.022] BF: 99.474 ROPE Bound: 0.005 In ROPE: 0 BF in ROPE: 0.011 RHat: 1.001 | Mode: 0.017 95% CI: [-0.014,0.039] BF: 0.417 ROPE Bound: 0.005 In ROPE: 0.2 BF in ROPE: 2.811 RHat: 1 |
|  | Family: student Num. obs: 3260 Num. groups: 34 Bayes R2: 0.055 | Family: student Num. obs: 3265 Num. groups: 34 Bayes R2: 0.012 |

*Figure S2.* Priors (light grey) and posteriors (dark grey) from the models linking single-trial RR interval changes to (A) outcomes and (B) model-derived prediction errors (PE), corresponding to *Hypothesis 2*, for the second heartbeat after feedback, R2. The dashed lines mark the respective ROPE (region of practical equivalence). BF is Bayes Factor, here it is an evidence ratio for coefficients being outside of ROPE.

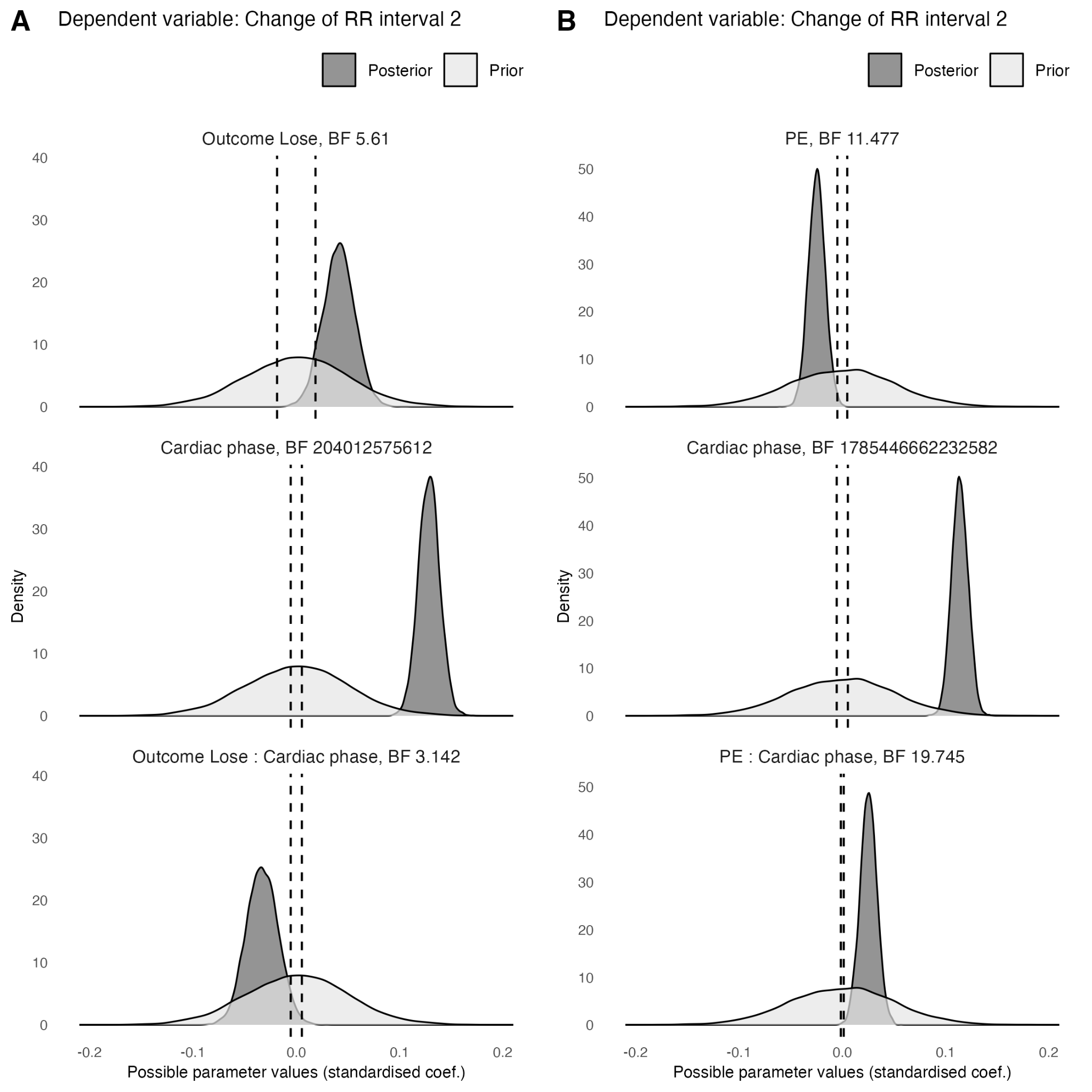

#### 2.3. Outcome valence

*Table S7.* The single-trial associations between the valence of outcome (contrast: lose compared to win) and the degree of change of the current interbeat interval (IBI) compared to the previous one, for the three heartbeats before and the four heartbeats after feedback. ROPE – region of practical equivalence, BF – Bayes Factor. Cardiac phase is a continuous measure of distance from the last heartbeat before feedback to feedback, scaled by the distance from the last heartbeat before to the first heartbeat after feedback. All values are standardised.

| Parameter | Dependent variable: Change of IBI -3 | Dependent variable: Change of IBI -2 | Dependent variable: Change of IBI -1 | Dependent variable: Change of IBI 1 | Dependent variable: Change of IBI 2 | Dependent variable: Change of IBI 3 | Dependent variable: Change of IBI 4 |
| --- | --- | --- | --- | --- | --- | --- | --- |
| Intercept | Mode: -0.011 95% CI: [-0.038,0.012] BF: 0.008 ROPE Bound: 0.009 In ROPE: 0.354 BF in ROPE: 184 RHat: 1.002 | Mode: -0.014 95% CI: [-0.057,0.01] BF: 0.016 ROPE Bound: 0.013 In ROPE: 0.249 BF in ROPE: 86.725 RHat: 1.004 | Mode: 0.01 95% CI: [-0.049,0.06] BF: 0.01 ROPE Bound: 0.018 In ROPE: 0.523 BF in ROPE: 177 RHat: 1.001 | Mode: -0.021 95% CI: [-0.063,0.053] BF: 0.011 ROPE Bound: 0.018 In ROPE: 0.481 BF in ROPE: 155 RHat: 1.002 | Mode: 0.037 95% CI: [-0.047,0.069] BF: 0.011 ROPE Bound: 0.019 In ROPE: 0.485 BF in ROPE: 148 RHat: 1.002 | Mode: 0.07 95% CI: [-0.053,0.127] BF: 0.022 ROPE Bound: 0.02 In ROPE: 0.285 BF in ROPE: 64.128 RHat: 1.004 | Mode: 0.015 95% CI: [-0.044,0.116] BF: 0.024 ROPE Bound: 0.017 In ROPE: 0.216 BF in ROPE: 50.454 RHat: 1.007 |
| Outcome Lose | Mode: 0.018 95% CI: [-0.012,0.019] BF: 0.182 ROPE Bound: 0.009 In ROPE: 0.725 BF in ROPE: 13.278 RHat: 1 | Mode: -0.011 95% CI: [-0.03,0.014] BF: 0.265 ROPE Bound: 0.013 In ROPE: 0.68 BF in ROPE: 6.727 RHat: 1.001 | Mode: 0.014 95% CI: [-0.018,0.045] BF: 0.499 ROPE Bound: 0.018 In ROPE: 0.571 BF in ROPE: 3.021 RHat: 1.001 | Mode: -0.019 95% CI: [-0.039,0.02] BF: 0.35 ROPE Bound: 0.018 In ROPE: 0.731 BF in ROPE: 5.783 RHat: 1 | Mode: 0.033 95% CI: [0.013,0.072] BF: 10.805 ROPE Bound: 0.019 In ROPE: 0.048 BF in ROPE: 0.178 RHat: 1.001 | Mode: 0.011 95% CI: [-0.029,0.035] BF: 0.334 ROPE Bound: 0.02 In ROPE: 0.796 BF in ROPE: 7.01 RHat: 1 | Mode: -0.037 95% CI: [-0.07,-0.015] BF: 25.648 ROPE Bound: 0.017 In ROPE: 0.013 BF in ROPE: 0.1 RHat: 1.001 |
| Cardiac phase | Mode: -0.01 95% CI: [-0.015,0.005] BF: 0.18 ROPE Bound: 0.003 In ROPE: 0.249 BF in ROPE: 7.173 RHat: 1 | Mode: 0.019 95% CI: [0.001,0.029] BF: 1.433 ROPE Bound: 0.004 In ROPE: 0.029 BF in ROPE: 0.81 RHat: 1.001 | Mode: 0.033 95% CI: [0.022,0.064] BF: 241 ROPE Bound: 0.005 In ROPE: 0 BF in ROPE: 0.005 RHat: 1.002 | Mode: -0.081 95% CI: [-0.098,-0.058] BF: 1588650 ROPE Bound: 0.005 In ROPE: 0 BF in ROPE: 0 RHat: 1.001 | Mode: 0.127 95% CI: [0.109,0.149] BF: 240794975575 ROPE Bound: 0.005 In ROPE: 0 BF in ROPE: 0 RHat: 1 | Mode: -0.018 95% CI: [-0.025,0.018] BF: 0.234 ROPE Bound: 0.006 In ROPE: 0.407 BF in ROPE: 6.093 RHat: 1.001 | Mode: -0.012 95% CI: [-0.024,0.012] BF: 0.217 ROPE Bound: 0.005 In ROPE: 0.366 BF in ROPE: 6.115 RHat: 1 |
| Outcome Lose : Cardiac phase | Mode: 0.014 95% CI: [-0.01,0.021] BF: 0.192 ROPE Bound: 0.003 In ROPE: 0.237 BF in ROPE: 6.569 RHat: 1 | Mode: 0.011 95% CI: [-0.023,0.02] BF: 0.229 ROPE Bound: 0.004 In ROPE: 0.267 BF in ROPE: 5.5 RHat: 1.001 | Mode: -0.033 95% CI: [-0.043,0.019] BF: 0.383 ROPE Bound: 0.005 In ROPE: 0.228 BF in ROPE: 3.018 RHat: 1 | Mode: 0.013 95% CI: [-0.023,0.038] BF: 0.344 ROPE Bound: 0.005 In ROPE: 0.255 BF in ROPE: 3.52 RHat: 1 | Mode: -0.032 95% CI: [-0.062,-0.002] BF: 3.256 ROPE Bound: 0.005 In ROPE: 0.011 BF in ROPE: 0.318 RHat: 1.002 | Mode: 0.044 95% CI: [-0.002,0.061] BF: 1.557 ROPE Bound: 0.006 In ROPE: 0.055 BF in ROPE: 0.661 RHat: 1.001 | Mode: -0.017 95% CI: [-0.047,0.005] BF: 0.74 ROPE Bound: 0.005 In ROPE: 0.113 BF in ROPE: 1.378 RHat: 1 |
|  | Family: student Num. obs: 9808 Num. groups: 34 Bayes R2: 0.004 | Family: student Num. obs: 9808 Num. groups: 34 Bayes R2: 0.007 | Family: student Num. obs: 9808 Num. groups: 34 Bayes R2: 0.021 | Family: student Num. obs: 9808 Num. groups: 34 Bayes R2: 0.026 | Family: student Num. obs: 9808 Num. groups: 34 Bayes R2: 0.035 | Family: student Num. obs: 9808 Num. groups: 34 Bayes R2: 0.054 | Family: student Num. obs: 9808 Num. groups: 34 Bayes R2: 0.036 |

*Table S8.* Separately only within systole and diastole trials (first and last 1/3 of the cardiac cycle), the single-trial associations between the valence of outcome (contrast: lose compared to win) and the degree of change of the current interbeat interval (IBI) compared to the previous one, for the second heartbeat after feedback. ROPE – region of practical equivalence, BF – Bayes Factor. All values are standardised.

| Parameter | Dependent variable: Change of IBI 2 only in systole | Dependent variable: Change of IBI 2 only in diastole |
| --- | --- | --- |
| Intercept | Mode: -0.221 95% CI: [-0.26,-0.08] BF: 10.028 ROPE Bound: 0.019 In ROPE: 0 BF in ROPE: 0.123 RHat: 1.001 | Mode: 0.16 95% CI: [0.109,0.215] BF: 554 ROPE Bound: 0.019 In ROPE: 0 BF in ROPE: 0.003 RHat: 1.001 |
| Outcome Lose | Mode: 0.107 95% CI: [0.019,0.118] BF: 19.521 ROPE Bound: 0.019 In ROPE: 0 BF in ROPE: 0.057 RHat: 1 | Mode: -0.034 95% CI: [-0.057,0.041] BF: 0.529 ROPE Bound: 0.019 In ROPE: 0.553 BF in ROPE: 2.604 RHat: 1 |
|  | Family: student Num. obs: 3260 Num. groups: 34 Bayes R2: 0.053 | Family: student Num. obs: 3265 Num. groups: 34 Bayes R2: 0.012 |

#### 2.4. Unsigned prediction error

*Table S9*. The single-trial associations between the continuous measure of unsigned prediction error (PE) and the degree of change of the current interbeat interval (IBI) compared to the previous one, for three heartbeats before and four heartbeats after feedback. ROPE – region of practical equivalence, BF – Bayes Factor. Cardiac phase is a continuous measure of distance from the last heartbeat before feedback to feedback, scaled by the distance from the last heartbeat before to the first heartbeat after feedback. All values are standardised.

| Parameter | Dependent variable: Change of IBI -3 | Dependent variable: Change of IBI -2 | Dependent variable: Change of IBI -1 | Dependent variable: Change of IBI 1 | Dependent variable: Change of IBI 2 | Dependent variable: Change of IBI 3 | Dependent variable: Change of IBI 4 |
| --- | --- | --- | --- | --- | --- | --- | --- |
| Intercept | Mode: -0.014 95% CI: [-0.036,0.014] BF: 0.007 ROPE Bound: 0.009 In ROPE: 0.401 BF in ROPE: 236 RHat: 1.003 | Mode: -0.019 95% CI: [-0.059,0.006] BF: 0.023 ROPE Bound: 0.013 In ROPE: 0.181 BF in ROPE: 61.745 RHat: 1.004 | Mode: 0.014 95% CI: [-0.044,0.064] BF: 0.011 ROPE Bound: 0.018 In ROPE: 0.505 BF in ROPE: 170 RHat: 1.001 | Mode: -0.027 95% CI: [-0.067,0.044] BF: 0.01 ROPE Bound: 0.018 In ROPE: 0.501 BF in ROPE: 170 RHat: 1.002 | Mode: 0.018 95% CI: [-0.027,0.085] BF: 0.018 ROPE Bound: 0.019 In ROPE: 0.319 BF in ROPE: 81.196 RHat: 1.001 | Mode: 0.129 95% CI: [-0.057,0.123] BF: 0.023 ROPE Bound: 0.02 In ROPE: 0.278 BF in ROPE: 63.375 RHat: 1.006 | Mode: 0.022 95% CI: [-0.058,0.099] BF: 0.016 ROPE Bound: 0.017 In ROPE: 0.32 BF in ROPE: 87.143 RHat: 1.004 |
| Unsigned PE | Mode: -0.002 95% CI: [-0.01,0.006] BF: 0.098 ROPE Bound: 0.003 In ROPE: 0.533 BF in ROPE: 18.622 RHat: 1 | Mode: 0.003 95% CI: [-0.009,0.014] BF: 0.127 ROPE Bound: 0.004 In ROPE: 0.554 BF in ROPE: 14.188 RHat: 1 | Mode: 0.012 95% CI: [0,0.032] BF: 1.127 ROPE Bound: 0.006 In ROPE: 0.1 BF in ROPE: 1.133 RHat: 1.001 | Mode: 0.012 95% CI: [-0.011,0.019] BF: 0.187 ROPE Bound: 0.006 In ROPE: 0.526 BF in ROPE: 9.334 RHat: 1.001 | Mode: 0.024 95% CI: [0.01,0.042] BF: 25.752 ROPE Bound: 0.007 In ROPE: 0 BF in ROPE: 0.072 RHat: 1.001 | Mode: 0.011 95% CI: [-0.001,0.033] BF: 1.041 ROPE Bound: 0.007 In ROPE: 0.12 BF in ROPE: 1.214 RHat: 1 | Mode: -0.016 95% CI: [-0.032,-0.003] BF: 2.993 ROPE Bound: 0.006 In ROPE: 0.028 BF in ROPE: 0.489 RHat: 1.001 |
| Cardiac phase | Mode: -0.005 95% CI: [-0.012,0.004] BF: 0.111 ROPE Bound: 0.003 In ROPE: 0.381 BF in ROPE: 12.659 RHat: 1 | Mode: 0.013 95% CI: [0.003,0.025] BF: 3.701 ROPE Bound: 0.004 In ROPE: 0 BF in ROPE: 0.373 RHat: 1.002 | Mode: 0.048 95% CI: [0.022,0.054] BF: 1453 ROPE Bound: 0.005 In ROPE: 0 BF in ROPE: 0.001 RHat: 1.001 | Mode: -0.076 95% CI: [-0.09,-0.059] BF: 62744005 ROPE Bound: 0.005 In ROPE: 0 BF in ROPE: 0 RHat: 1.001 | Mode: 0.112 95% CI: [0.098,0.129] BF: 58687441147092 ROPE Bound: 0.005 In ROPE: 0 BF in ROPE: 0 RHat: 1 | Mode: 0.011 95% CI: [-0.009,0.025] BF: 0.252 ROPE Bound: 0.006 In ROPE: 0.357 BF in ROPE: 5.016 RHat: 1 | Mode: -0.023 95% CI: [-0.028,0] BF: 1.06 ROPE Bound: 0.005 In ROPE: 0.08 BF in ROPE: 1.18 RHat: 1.001 |
| Unsigned PE : Cardiac phase | Mode: -0.002 95% CI: [-0.014,0.002] BF: 0.214 ROPE Bound: 0.001 In ROPE: 0.085 BF in ROPE: 5.292 RHat: 1 | Mode: -0.011 95% CI: [-0.016,0.006] BF: 0.186 ROPE Bound: 0.001 In ROPE: 0.119 BF in ROPE: 5.855 RHat: 1.001 | Mode: 0.007 95% CI: [-0.017,0.015] BF: 0.166 ROPE Bound: 0.002 In ROPE: 0.187 BF in ROPE: 7.184 RHat: 1.001 | Mode: 0.019 95% CI: [-0.01,0.021] BF: 0.211 ROPE Bound: 0.002 In ROPE: 0.142 BF in ROPE: 5.329 RHat: 1 | Mode: -0.022 95% CI: [-0.029,0.003] BF: 0.675 ROPE Bound: 0.002 In ROPE: 0.05 BF in ROPE: 1.535 RHat: 1 | Mode: 0.02 95% CI: [0.006,0.039] BF: 4.252 ROPE Bound: 0.002 In ROPE: 0 BF in ROPE: 0.229 RHat: 1 | Mode: -0.014 95% CI: [-0.02,0.007] BF: 0.206 ROPE Bound: 0.002 In ROPE: 0.139 BF in ROPE: 5.53 RHat: 1.002 |
|  | Family: student Num. obs: 9808 Num. groups: 34 Bayes R2: 0.004 | Family: student Num. obs: 9808 Num. groups: 34 Bayes R2: 0.007 | Family: student Num. obs: 9808 Num. groups: 34 Bayes R2: 0.021 | Family: student Num. obs: 9808 Num. groups: 34 Bayes R2: 0.026 | Family: student Num. obs: 9808 Num. groups: 34 Bayes R2: 0.035 | Family: student Num. obs: 9808 Num. groups: 34 Bayes R2: 0.054 | Family: student Num. obs: 9808 Num. groups: 34 Bayes R2: 0.036 |

*Table S10*. Separately only within systole and diastole trials (first and last 1/3 of the cardiac cycle), the single-trial associations between the continuous measure of unsigned prediction error (PE) and the degree of change of the current interbeat interval (IBI) compared to the previous one, for the second and third heartbeats after feedback. ROPE – region of practical equivalence, BF – Bayes Factor. All values are standardised.

| Parameter | Dependent variable: Change of IBI 2 only in systole | Dependent variable: Change of IBI 3 only in systole | Dependent variable: Change of IBI 2 only in diastole | Dependent variable: Change of IBI 3 only in diastole |
| --- | --- | --- | --- | --- |
| Intercept | Mode: -0.157 95% CI: [-0.23,-0.046] BF: 1.089 ROPE Bound: 0.019 In ROPE: 0 BF in ROPE: 0.992 RHat: 1.002 | Mode: 0.027 95% CI: [-0.059,0.116] BF: 0.021 ROPE Bound: 0.02 In ROPE: 0.285 BF in ROPE: 62.858 RHat: 1.003 | Mode: 0.158 95% CI: [0.111,0.207] BF: 693 ROPE Bound: 0.019 In ROPE: 0 BF in ROPE: 0.002 RHat: 1.001 | Mode: 0.067 95% CI: [-0.048,0.15] BF: 0.036 ROPE Bound: 0.02 In ROPE: 0.183 BF in ROPE: 36.955 RHat: 1.002 |
| Unsigned PE | Mode: 0.032 95% CI: [0.013,0.069] BF: 12.88 ROPE Bound: 0.007 In ROPE: 0 BF in ROPE: 0.085 RHat: 1 | Mode: -0.025 95% CI: [-0.048,0.009] BF: 0.668 ROPE Bound: 0.007 In ROPE: 0.181 BF in ROPE: 1.587 RHat: 1.001 | Mode: 0.02 95% CI: [-0.018,0.036] BF: 0.312 ROPE Bound: 0.007 In ROPE: 0.32 BF in ROPE: 3.834 RHat: 1.001 | Mode: 0.017 95% CI: [0,0.058] BF: 2.131 ROPE Bound: 0.007 In ROPE: 0.044 BF in ROPE: 0.484 RHat: 1 |
|  | Family: student Num. obs: 3260 Num. groups: 34 Bayes R2: 0.054 | Family: student Num. obs: 3260 Num. groups: 34 Bayes R2: 0.054 | Family: student Num. obs: 3265 Num. groups: 34 Bayes R2: 0.012 | Family: student Num. obs: 3265 Num. groups: 34 Bayes R2: 0.055 |

### 3. Neural responses and interbeat intervals

*Table S11.* Separately only within systole or diastole trials (first and last 1/3 of the cardiac cycle), the single-trial associations between the continuous measure of P3b amplitude and the degree of change in the current interbeat interval (IBI) compared to the previous one, for the first and second heartbeats after feedback. ROPE – region of practical equivalence, BF – Bayes Factor. All values are standardised.

| Parameter | Dependent variable: Change of IBI 1 only in systole | Dependent variable: Change of IBI 2 only in systole | Dependent variable: Change of IBI 1 only in diastole | Dependent variable: Change of IBI 2 only in diastole |
| --- | --- | --- | --- | --- |
| Intercept | Mode: 0.107 95% CI: [0.031,0.157] BF: 0.714 ROPE Bound: 0.018 In ROPE: 0 BF in ROPE: 1.731 RHat: 1.001 | Mode: -0.195 95% CI: [-0.241,-0.053] BF: 1.631 ROPE Bound: 0.019 In ROPE: 0 BF in ROPE: 0.735 RHat: 1.001 | Mode: -0.128 95% CI: [-0.167,-0.025] BF: 0.378 ROPE Bound: 0.018 In ROPE: 0 BF in ROPE: 3.292 RHat: 1.001 | Mode: 0.172 95% CI: [0.115,0.208] BF: 610 ROPE Bound: 0.019 In ROPE: 0 BF in ROPE: 0.002 RHat: 1.001 |
| P3b | Mode: 0.028 95% CI: [0.014,0.068] BF: 19.028 ROPE Bound: 0.003 In ROPE: 0 BF in ROPE: 0.052 RHat: 1 | Mode: -0.018 95% CI: [-0.025,0.032] BF: 0.301 ROPE Bound: 0.003 In ROPE: 0.158 BF in ROPE: 3.596 RHat: 1 | Mode: 0.019 95% CI: [-0.009,0.043] BF: 0.624 ROPE Bound: 0.003 In ROPE: 0.076 BF in ROPE: 1.695 RHat: 1 | Mode: 0.049 95% CI: [0.027,0.082] BF: 412 ROPE Bound: 0.003 In ROPE: 0 BF in ROPE: 0.002 RHat: 1 |
|  | Family: student Num. obs: 3212 Num. groups: 34 Bayes R2: 0.026 | Family: student Num. obs: 3212 Num. groups: 34 Bayes R2: 0.056 | Family: student Num. obs: 3229 Num. groups: 34 Bayes R2: 0.033 | Family: student Num. obs: 3229 Num. groups: 34 Bayes R2: 0.014 |

*Figure S3.* Priors (light grey) and posteriors (dark grey) from the models linking single-trial RR interval changes to P3b amplitudes, corresponding to *Hypothesis 3*, for (A) the first and (B) second heartbeats after feedback. The dashed lines mark the respective ROPE (region of practical equivalence). BF is Bayes Factor, here it is an evidence ratio for coefficients being outside of ROPE.

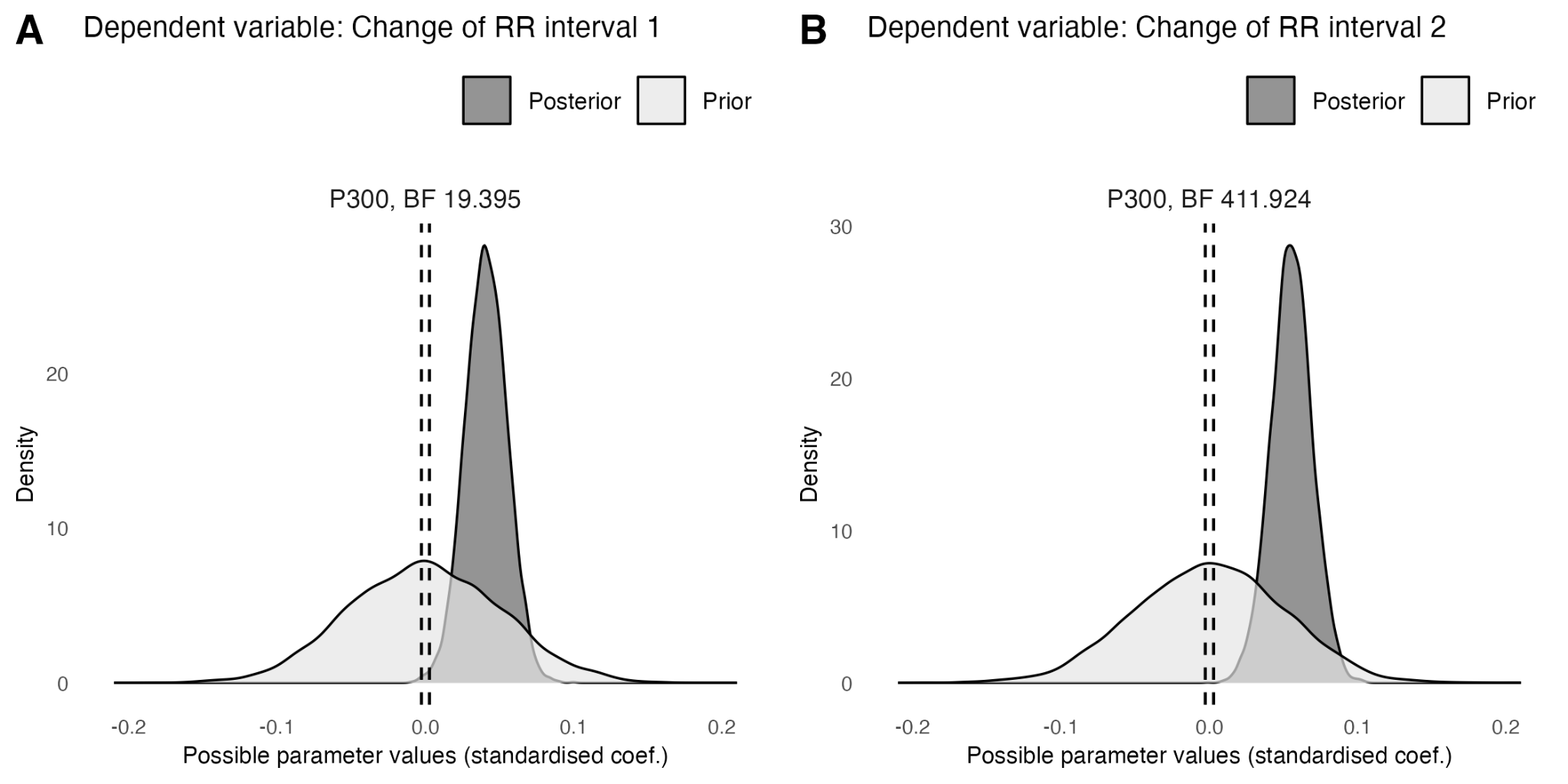

### 4. Control and sensitivity analyses

#### 4.1. Expectation

##### 4.1.1. Average IBI or RMSSD during rest

*Table S12.* The single-trial associations between the continuous measure of expectation and the degree of change of the current interbeat interval (IBI) compared to the previous one, for the last heartbeat before and the third one after feedback, controlling for the resting average IBI or resting RMSSD. ROPE – region of practical equivalence, BF – Bayes Factor. Cardiac phase is a continuous measure of distance from the last heartbeat before feedback to feedback, scaled by the distance from the last heartbeat before to the first heartbeat after feedback (IBI 1). All values are standardised.

| Parameter | Dependent variable: Change of IBI -1 | Dependent variable: Change of IBI 3 | Dependent variable: Change of IBI -1 | Dependent variable: Change of IBI 3 |
| --- | --- | --- | --- | --- |
| Intercept | Mode: 0.019 95% CI: [-0.043,0.066] BF: 0.011 ROPE Bound: 0.018 In ROPE: 0.498 BF in ROPE: 175 RHat: 1.004 | Mode: 0.05 95% CI: [-0.06,0.134] BF: 0.023 ROPE Bound: 0.02 In ROPE: 0.26 BF in ROPE: 61.189 RHat: 1.007 | Mode: 0.019 95% CI: [-0.043,0.066] BF: 0.011 ROPE Bound: 0.018 In ROPE: 0.489 BF in ROPE: 161 RHat: 1.009 | Mode: 0.102 95% CI: [-0.05,0.122] BF: 0.022 ROPE Bound: 0.02 In ROPE: 0.263 BF in ROPE: 59.521 RHat: 1.004 |
| Expectation | Mode: -0.033 95% CI: [-0.048,-0.014] BF: 69.049 ROPE Bound: 0.006 In ROPE: 0 BF in ROPE: 0.022 RHat: 1 | Mode: 0.035 95% CI: [0.018,0.053] BF: 250 ROPE Bound: 0.006 In ROPE: 0 BF in ROPE: 0.007 RHat: 1 | Mode: -0.041 95% CI: [-0.048,-0.015] BF: 164 ROPE Bound: 0.006 In ROPE: 0 BF in ROPE: 0.012 RHat: 1.001 | Mode: 0.047 95% CI: [0.017,0.052] BF: 275 ROPE Bound: 0.006 In ROPE: 0 BF in ROPE: 0.007 RHat: 1 |
| Cardiac phase | Mode: 0.039 95% CI: [0.023,0.055] BF: 1165 ROPE Bound: 0.005 In ROPE: 0 BF in ROPE: 0.001 RHat: 1 | Mode: 0.008 95% CI: [-0.013,0.021] BF: 0.197 ROPE Bound: 0.006 In ROPE: 0.464 BF in ROPE: 7.88 RHat: 1 | Mode: 0.049 95% CI: [0.029,0.063] BF: 5845 ROPE Bound: 0.005 In ROPE: 0 BF in ROPE: 0 RHat: 1.002 | Mode: 0.014 95% CI: [-0.014,0.021] BF: 0.189 ROPE Bound: 0.006 In ROPE: 0.478 BF in ROPE: 8.491 RHat: 1 |
| Rest Average IBI | Mode: -0.011 95% CI: [-0.059,0.035] BF: 0.527 ROPE Bound: 0.004 In ROPE: 0.127 BF in ROPE: 1.945 RHat: 1.003 | Mode: -0.044 95% CI: [-0.077,0.056] BF: 0.761 ROPE Bound: 0.004 In ROPE: 0.094 BF in ROPE: 1.35 RHat: 1.002 |  |  |
| Expectation : Cardiac phase | Mode: -0.016 95% CI: [-0.019,0.015] BF: 0.177 ROPE Bound: 0.002 In ROPE: 0.154 BF in ROPE: 6.266 RHat: 1.001 | Mode: 0.017 95% CI: [-0.015,0.019] BF: 0.166 ROPE Bound: 0.002 In ROPE: 0.16 BF in ROPE: 6.207 RHat: 1 | Mode: 0.003 95% CI: [-0.019,0.014] BF: 0.173 ROPE Bound: 0.002 In ROPE: 0.153 BF in ROPE: 6.55 RHat: 1.002 | Mode: -0.01 95% CI: [-0.019,0.015] BF: 0.171 ROPE Bound: 0.002 In ROPE: 0.162 BF in ROPE: 6.747 RHat: 1 |
| Cardiac phase : Rest Average IBI | Mode: -0.011 95% CI: [-0.02,0.016] BF: 0.185 ROPE Bound: 0.001 In ROPE: 0.113 BF in ROPE: 6.086 RHat: 1 | Mode: -0.031 95% CI: [-0.045,-0.008] BF: 10.204 ROPE Bound: 0.001 In ROPE: 0 BF in ROPE: 0.1 RHat: 1 |  |  |
| Rest RMSSD |  |  | Mode: -0.014 95% CI: [-0.057,0.035] BF: 0.518 ROPE Bound: 0.003 In ROPE: 0.103 BF in ROPE: 2.036 RHat: 1.001 | Mode: -0.039 95% CI: [-0.119,0.002] BF: 3.337 ROPE Bound: 0.003 In ROPE: 0.016 BF in ROPE: 0.287 RHat: 1.001 |
| Cardiac phase : Rest RMSSD |  |  | Mode: 0.018 95% CI: [0.006,0.051] BF: 4.601 ROPE Bound: 0.001 In ROPE: 0 BF in ROPE: 0.217 RHat: 1 | Mode: -0.023 95% CI: [-0.047,0.002] BF: 1.274 ROPE Bound: 0.001 In ROPE: 0.015 BF in ROPE: 0.765 RHat: 1 |
|  | Family: student Num. obs: 9808 Num. groups: 34 Bayes R2: 0.022 | Family: student Num. obs: 9808 Num. groups: 34 Bayes R2: 0.055 | Family: student Num. obs: 9808 Num. groups: 34 Bayes R2: 0.023 | Family: student Num. obs: 9808 Num. groups: 34 Bayes R2: 0.055 |

##### 4.1.2. Average IBI or RMSSD within a trial, before or after feedback

*Table S13.* The single-trial associations between the continuous measure of expectation and the degree of change of the current interbeat interval (IBI) compared to the previous one, for the last heartbeat before and the third one after feedback, controlling for the average IBI or RMSSD before feedback. ROPE – region of practical equivalence, BF – Bayes Factor. Cardiac phase is a continuous measure of distance from the last heartbeat before feedback to feedback, scaled by the distance from the last heartbeat before to the first heartbeat after feedback (IBI 1). All values are standardised.

| Parameter | Dependent variable: Change of IBI -1 | Dependent variable: Change of IBI 3 | Dependent variable: Change of IBI -1 | Dependent variable: Change of IBI 3 |
| --- | --- | --- | --- | --- |
| Intercept | Mode: -0.011 95% CI: [-0.075,0.087] BF: 0.014 ROPE Bound: 0.018 In ROPE: 0.373 BF in ROPE: 102 RHat: 1.002 | Mode: 0.032 95% CI: [-0.062,0.124] BF: 0.02 ROPE Bound: 0.02 In ROPE: 0.273 BF in ROPE: 61.985 RHat: 1.006 | Mode: 0.021 95% CI: [-0.055,0.051] BF: 0.01 ROPE Bound: 0.018 In ROPE: 0.542 BF in ROPE: 190 RHat: 1.003 | Mode: 0.052 95% CI: [-0.05,0.121] BF: 0.022 ROPE Bound: 0.02 In ROPE: 0.278 BF in ROPE: 62.805 RHat: 1.003 |
| Expectation | Mode: -0.033 95% CI: [-0.046,-0.012] BF: 533 ROPE Bound: 0.006 In ROPE: 0 BF in ROPE: 0.021 RHat: 1.001 | Mode: 0.042 95% CI: [0.017,0.052] BF: 150 ROPE Bound: 0.006 In ROPE: 0 BF in ROPE: 0.01 RHat: 1 | Mode: -0.025 95% CI: [-0.046,-0.013] BF: 112 ROPE Bound: 0.006 In ROPE: 0 BF in ROPE: 0.015 RHat: 1 | Mode: 0.033 95% CI: [0.017,0.051] BF: 222 ROPE Bound: 0.006 In ROPE: 0 BF in ROPE: 0.007 RHat: 1 |
| Cardiac phase | Mode: 0.039 95% CI: [0.025,0.057] BF: 1930 ROPE Bound: 0.005 In ROPE: 0 BF in ROPE: 0.001 RHat: 1 | Mode: 0.011 95% CI: [-0.013,0.02] BF: 0.185 ROPE Bound: 0.006 In ROPE: 0.466 BF in ROPE: 8.282 RHat: 1.001 | Mode: 0.053 95% CI: [0.037,0.072] BF: 715106 ROPE Bound: 0.005 In ROPE: 0 BF in ROPE: 0 RHat: 1.001 | Mode: 0.006 95% CI: [-0.011,0.023] BF: 0.217 ROPE Bound: 0.006 In ROPE: 0.433 BF in ROPE: 7.03 RHat: 1.001 |
| Average IBI before feedback | Mode: -0.248 95% CI: [-0.27,-0.207] BF: 2586943281618798 ROPE Bound: 0.001 In ROPE: 0 BF in ROPE: 0 RHat: 1.001 | Mode: -0.151 95% CI: [-0.184,-0.124] BF: 185510070 ROPE Bound: 0.001 In ROPE: 0 BF in ROPE: 0 RHat: 1 |  |  |
| Expectation : Cardiac phase | Mode: 0.001 95% CI: [-0.021,0.012] BF: 0.203 ROPE Bound: 0.002 In ROPE: 0.142 BF in ROPE: 5.561 RHat: 1 | Mode: -0.003 95% CI: [-0.017,0.017] BF: 0.169 ROPE Bound: 0.002 In ROPE: 0.167 BF in ROPE: 6.617 RHat: 1 | Mode: -0.011 95% CI: [-0.02,0.013] BF: 0.177 ROPE Bound: 0.002 In ROPE: 0.147 BF in ROPE: 6.108 RHat: 1 | Mode: 0.003 95% CI: [-0.018,0.016] BF: 0.167 ROPE Bound: 0.002 In ROPE: 0.169 BF in ROPE: 6.59 RHat: 1.001 |
| Cardiac phase : Average IBI before feedback | Mode: 0.018 95% CI: [-0.001,0.035] BF: 0.907 ROPE Bound: 0 In ROPE: 0.004 BF in ROPE: 1.068 RHat: 1 | Mode: -0.026 95% CI: [-0.046,-0.008] BF: 11.655 ROPE Bound: 0 In ROPE: 0 BF in ROPE: 0.089 RHat: 1 |  |  |
| RMSSD before feedback |  |  | Mode: -0.075 95% CI: [-0.095,-0.042] BF: 2455 ROPE Bound: 0 In ROPE: 0 BF in ROPE: 0 RHat: 1.001 | Mode: -0.168 95% CI: [-0.193,-0.12] BF: 35498439 ROPE Bound: 0 In ROPE: 0 BF in ROPE: 0 RHat: 1 |
| Cardiac phase : RMSSD before feedback |  |  | Mode: 0.103 95% CI: [0.044,0.101] BF: 10188 ROPE Bound: 0 In ROPE: 0 BF in ROPE: 0 RHat: 1.001 | Mode: -0.031 95% CI: [-0.057,0.003] BF: 1.437 ROPE Bound: 0 In ROPE: 0.001 BF in ROPE: 0.685 RHat: 1.001 |
|  | Family: student Num. obs: 9808 Num. groups: 34 Bayes R2: 0.047 | Family: student Num. obs: 9808 Num. groups: 34 Bayes R2: 0.065 | Family: student Num. obs: 9808 Num. groups: 34 Bayes R2: 0.027 | Family: student Num. obs: 9808 Num. groups: 34 Bayes R2: 0.076 |

*Table S14.* The single-trial associations between the continuous measure of expectation and the degree of change of the current interbeat interval (IBI) compared to the previous one, for the last heartbeat before and the third one after feedback, controlling for the average IBI or RMSSD after feedback. ROPE – region of practical equivalence, BF – Bayes Factor. Cardiac phase is a continuous measure of distance from the last heartbeat before feedback to feedback, scaled by the distance from the last heartbeat before to the first heartbeat after feedback (IBI 1). All values are standardised.

| Parameter | Dependent variable: Change of IBI -1 | Dependent variable: Change of IBI 3 | Dependent variable: Change of IBI -1 | Dependent variable: Change of IBI 3 |
| --- | --- | --- | --- | --- |
| Intercept | Mode: 0.064 95% CI: [-0.088,0.137] BF: 0.024 ROPE Bound: 0.018 In ROPE: 0.262 BF in ROPE: 58.119 RHat: 1.005 | Mode: 0.081 95% CI: [-0.058,0.12] BF: 0.02 ROPE Bound: 0.02 In ROPE: 0.291 BF in ROPE: 63.403 RHat: 1.007 | Mode: 0.026 95% CI: [-0.041,0.077] BF: 0.013 ROPE Bound: 0.018 In ROPE: 0.427 BF in ROPE: 118 RHat: 1.005 | Mode: -0.101 95% CI: [-0.173,0.004] BF: 0.093 ROPE Bound: 0.02 In ROPE: 0.058 BF in ROPE: 12.158 RHat: 1.009 |
| Expectation | Mode: -0.035 95% CI: [-0.048,-0.015] BF: 117 ROPE Bound: 0.006 In ROPE: 0 BF in ROPE: 0.012 RHat: 1 | Mode: 0.039 95% CI: [0.016,0.05] BF: 144 ROPE Bound: 0.006 In ROPE: 0 BF in ROPE: 0.011 RHat: 1.001 | Mode: -0.023 95% CI: [-0.048,-0.014] BF: 63.223 ROPE Bound: 0.006 In ROPE: 0 BF in ROPE: 0.023 RHat: 1.001 | Mode: 0.042 95% CI: [0.018,0.049] BF: 631 ROPE Bound: 0.006 In ROPE: 0 BF in ROPE: 0.003 RHat: 1 |
| Cardiac phase | Mode: 0.04 95% CI: [0.018,0.049] BF: 377209 ROPE Bound: 0.005 In ROPE: 0 BF in ROPE: 0 RHat: 1 | Mode: 0.012 95% CI: [-0.014,0.019] BF: 0.191 ROPE Bound: 0.006 In ROPE: 0.476 BF in ROPE: 8.088 RHat: 1 | Mode: 0.024 95% CI: [0.01,0.042] BF: 23.953 ROPE Bound: 0.005 In ROPE: 0 BF in ROPE: 0.06 RHat: 1 | Mode: -0.016 95% CI: [-0.022,0.012] BF: 0.192 ROPE Bound: 0.006 In ROPE: 0.452 BF in ROPE: 7.521 RHat: 1.001 |
| Average IBI after feedback | Mode: 0.334 95% CI: [0.281,0.343] BF: 1.81099813166838e+24 ROPE Bound: 0.002 In ROPE: 0 BF in ROPE: 0 RHat: 1 | Mode: -0.117 95% CI: [-0.138,-0.078] BF: 205685 ROPE Bound: 0.002 In ROPE: 0 BF in ROPE: 0 RHat: 1 |  |  |
| Expectation : Cardiac phase | Mode: -0.012 95% CI: [-0.017,0.015] BF: 0.171 ROPE Bound: 0.002 In ROPE: 0.159 BF in ROPE: 6.58 RHat: 1 | Mode: -0.012 95% CI: [-0.017,0.017] BF: 0.172 ROPE Bound: 0.002 In ROPE: 0.163 BF in ROPE: 6.564 RHat: 1 | Mode: 0.002 95% CI: [-0.02,0.013] BF: 0.168 ROPE Bound: 0.002 In ROPE: 0.153 BF in ROPE: 6.638 RHat: 1.001 | Mode: -0.011 95% CI: [-0.018,0.012] BF: 0.175 ROPE Bound: 0.002 In ROPE: 0.165 BF in ROPE: 6.482 RHat: 1.001 |
| Cardiac phase : Average IBI after feedback | Mode: 0.015 95% CI: [-0.002,0.031] BF: 0.642 ROPE Bound: 0 In ROPE: 0.014 BF in ROPE: 1.493 RHat: 1 | Mode: -0.046 95% CI: [-0.059,-0.023] BF: 525 ROPE Bound: 0.001 In ROPE: 0 BF in ROPE: 0.002 RHat: 1 |  |  |
| RMSSD after feedback |  |  | Mode: 0.105 95% CI: [0.074,0.122] BF: 5382073 ROPE Bound: 0.001 In ROPE: 0 BF in ROPE: 0 RHat: 1.001 | Mode: -0.494 95% CI: [-0.523,-0.461] BF: 1.78707155953808e+32 ROPE Bound: 0.002 In ROPE: 0 BF in ROPE: 0 RHat: 1.001 |
| Cardiac phase : RMSSD after feedback |  |  | Mode: -0.049 95% CI: [-0.071,-0.032] BF: 2222 ROPE Bound: 0 In ROPE: 0 BF in ROPE: 0 RHat: 1.001 | Mode: -0.049 95% CI: [-0.068,-0.02] BF: 112 ROPE Bound: 0 In ROPE: 0 BF in ROPE: 0.009 RHat: 1.001 |
|  | Family: student Num. obs: 9808 Num. groups: 34 Bayes R2: 0.062 | Family: student Num. obs: 9808 Num. groups: 34 Bayes R2: 0.062 | Family: student Num. obs: 9808 Num. groups: 34 Bayes R2: 0.032 | Family: student Num. obs: 9808 Num. groups: 34 Bayes R2: 0.195 |

##### 4.1.3. Number of a trial in a block and approach/avoid decisions

*Table S15.* The single-trial associations between the continuous measure of expectation and the degree of change of the current interbeat interval (IBI) compared to the previous one, for the last heartbeat before and the third one after feedback, controlling for the number of trial in a block (learning effects) or approach/avoid decision trials. ROPE – region of practical equivalence, BF – Bayes Factor. Cardiac phase is a continuous measure of distance from the last heartbeat before feedback to feedback, scaled by the distance from the last heartbeat before to the first heartbeat after feedback (IBI 1). All values are standardised.

| Parameter | Dependent variable: Change of IBI -1 | Dependent variable: Change of IBI 3 | Dependent variable: Change of IBI -1 | Dependent variable: Change of IBI 3 |
| --- | --- | --- | --- | --- |
| Intercept | Mode: 0.034 95% CI: [-0.045,0.064] BF: 0.011 ROPE Bound: 0.018 In ROPE: 0.493 BF in ROPE: 162 RHat: 1.003 | Mode: 0.058 95% CI: [-0.058,0.122] BF: 0.02 ROPE Bound: 0.02 In ROPE: 0.285 BF in ROPE: 61.494 RHat: 1.006 | Mode: 0.046 95% CI: [-0.06,0.053] BF: 0.01 ROPE Bound: 0.018 In ROPE: 0.53 BF in ROPE: 177 RHat: 1.002 | Mode: 0.018 95% CI: [-0.058,0.131] BF: 0.024 ROPE Bound: 0.02 In ROPE: 0.26 BF in ROPE: 57.258 RHat: 1.005 |
| Expectation | Mode: -0.033 95% CI: [-0.049,-0.015] BF: 50.107 ROPE Bound: 0.006 In ROPE: 0 BF in ROPE: 0.025 RHat: 1.001 | Mode: 0.033 95% CI: [0.017,0.052] BF: 134 ROPE Bound: 0.006 In ROPE: 0 BF in ROPE: 0.011 RHat: 1.001 | Mode: -0.037 95% CI: [-0.05,-0.015] BF: 68.335 ROPE Bound: 0.006 In ROPE: 0 BF in ROPE: 0.022 RHat: 1 | Mode: 0.043 95% CI: [0.018,0.053] BF: 150 ROPE Bound: 0.006 In ROPE: 0 BF in ROPE: 0.009 RHat: 1.001 |
| Cardiac phase | Mode: 0.038 95% CI: [0.023,0.056] BF: 2739 ROPE Bound: 0.005 In ROPE: 0 BF in ROPE: 0.001 RHat: 1 | Mode: 0.011 95% CI: [-0.009,0.024] BF: 0.255 ROPE Bound: 0.006 In ROPE: 0.376 BF in ROPE: 5.612 RHat: 1.001 | Mode: 0.064 95% CI: [0.037,0.079] BF: 39630 ROPE Bound: 0.005 In ROPE: 0 BF in ROPE: 0 RHat: 1.001 | Mode: -0.019 95% CI: [-0.03,0.014] BF: 0.284 ROPE Bound: 0.006 In ROPE: 0.336 BF in ROPE: 4.678 RHat: 1.001 |
| Trial in Block | Mode: -0.028 95% CI: [-0.042,-0.01] BF: 17.223 ROPE Bound: 0.005 In ROPE: 0 BF in ROPE: 0.077 RHat: 1 | Mode: 0.015 95% CI: [0.003,0.037] BF: 3.023 ROPE Bound: 0.006 In ROPE: 0.017 BF in ROPE: 0.414 RHat: 1.001 |  |  |
| Expectation : Cardiac phase | Mode: -0.012 95% CI: [-0.019,0.015] BF: 0.177 ROPE Bound: 0.002 In ROPE: 0.153 BF in ROPE: 6.132 RHat: 1 | Mode: -0.014 95% CI: [-0.017,0.016] BF: 0.181 ROPE Bound: 0.002 In ROPE: 0.165 BF in ROPE: 6.498 RHat: 1.002 | Mode: -0.005 95% CI: [-0.018,0.015] BF: 0.164 ROPE Bound: 0.002 In ROPE: 0.154 BF in ROPE: 6.618 RHat: 1.001 | Mode: -0.008 95% CI: [-0.02,0.015] BF: 0.174 ROPE Bound: 0.002 In ROPE: 0.156 BF in ROPE: 6.486 RHat: 1.001 |
| Cardiac phase : Trial in Block | Mode: 0.022 95% CI: [-0.002,0.03] BF: 0.614 ROPE Bound: 0.002 In ROPE: 0.046 BF in ROPE: 1.745 RHat: 1 | Mode: 0.014 95% CI: [-0.003,0.03] BF: 0.611 ROPE Bound: 0.002 In ROPE: 0.046 BF in ROPE: 1.743 RHat: 1 |  |  |
| Decision Avoid |  |  | Mode: 0.021 95% CI: [-0.005,0.059] BF: 1.356 ROPE Bound: 0.018 In ROPE: 0.286 BF in ROPE: 1.095 RHat: 1.001 | Mode: 0.014 95% CI: [-0.032,0.031] BF: 0.327 ROPE Bound: 0.02 In ROPE: 0.815 BF in ROPE: 7.663 RHat: 1 |
| Cardiac phase : Decision Avoid |  |  | Mode: -0.043 95% CI: [-0.077,-0.016] BF: 24.366 ROPE Bound: 0.005 In ROPE: 0 BF in ROPE: 0.041 RHat: 1 | Mode: 0.028 95% CI: [0.002,0.067] BF: 3.319 ROPE Bound: 0.006 In ROPE: 0.014 BF in ROPE: 0.308 RHat: 1 |
|  | Family: student Num. obs: 9808 Num. groups: 34 Bayes R2: 0.023 | Family: student Num. obs: 9808 Num. groups: 34 Bayes R2: 0.055 | Family: student Num. obs: 9808 Num. groups: 34 Bayes R2: 0.022 | Family: student Num. obs: 9808 Num. groups: 34 Bayes R2: 0.055 |

*Table S16.* Only in approach or avoid decision trials, the single-trial associations between the continuous measure of expectation and the degree of change of the current interbeat interval (IBI) compared to the previous one, for the last heartbeat before and the third one after feedback. Avoid trials did not have a movement confound. ROPE – region of practical equivalence, BF – Bayes Factor. Cardiac phase is a continuous measure of distance from the last heartbeat before feedback to feedback, scaled by the distance from the last heartbeat before to the first heartbeat after feedback (IBI 1). All values are standardised.

| Parameter | Dependent variable: Change of IBI -1 only in approach trials | Dependent variable: Change of IBI 3 only in approach trials | Dependent variable: Change of IBI -1 only in avoid trials | Dependent variable: Change of IBI 3 only in avoid trials |
| --- | --- | --- | --- | --- |
| Intercept | Mode: -0.022 95% CI: [-0.058,0.072] BF: 0.012 ROPE Bound: 0.019 In ROPE: 0.454 BF in ROPE: 136 RHat: 1.004 | Mode: 0.046 95% CI: [-0.053,0.129] BF: 0.028 ROPE Bound: 0.02 In ROPE: 0.235 BF in ROPE: 49.181 RHat: 1.002 | Mode: 0.066 95% CI: [-0.035,0.073] BF: 0.014 ROPE Bound: 0.018 In ROPE: 0.407 BF in ROPE: 119 RHat: 1.001 | Mode: 0.025 95% CI: [-0.08,0.11] BF: 0.017 ROPE Bound: 0.019 In ROPE: 0.331 BF in ROPE: 80.198 RHat: 1.002 |
| Expectation | Mode: -0.028 95% CI: [-0.05,-0.006] BF: 6.748 ROPE Bound: 0.006 In ROPE: 0 BF in ROPE: 0.179 RHat: 1.002 | Mode: 0.022 95% CI: [-0.002,0.042] BF: 1.124 ROPE Bound: 0.006 In ROPE: 0.086 BF in ROPE: 0.977 RHat: 1 | Mode: -0.04 95% CI: [-0.058,-0.006] BF: 6.311 ROPE Bound: 0.005 In ROPE: 0 BF in ROPE: 0.17 RHat: 1 | Mode: 0.057 95% CI: [0.023,0.077] BF: 185 ROPE Bound: 0.006 In ROPE: 0 BF in ROPE: 0.006 RHat: 1 |
| Cardiac phase | Mode: 0.072 95% CI: [0.041,0.083] BF: 18006 ROPE Bound: 0.005 In ROPE: 0 BF in ROPE: 0 RHat: 1 | Mode: -0.015 95% CI: [-0.03,0.013] BF: 0.289 ROPE Bound: 0.006 In ROPE: 0.317 BF in ROPE: 4.524 RHat: 1 | Mode: 0.013 95% CI: [-0.016,0.032] BF: 0.315 ROPE Bound: 0.005 In ROPE: 0.272 BF in ROPE: 3.848 RHat: 1 | Mode: 0.012 95% CI: [0.002,0.052] BF: 2.78 ROPE Bound: 0.006 In ROPE: 0.019 BF in ROPE: 0.402 RHat: 1 |
| Expectation : Cardiac phase | Mode: 0.012 95% CI: [-0.016,0.026] BF: 0.247 ROPE Bound: 0.002 In ROPE: 0.118 BF in ROPE: 4.562 RHat: 1 | Mode: 0.01 95% CI: [-0.016,0.028] BF: 0.248 ROPE Bound: 0.002 In ROPE: 0.129 BF in ROPE: 4.353 RHat: 1 | Mode: -0.013 95% CI: [-0.034,0.017] BF: 0.337 ROPE Bound: 0.002 In ROPE: 0.087 BF in ROPE: 3.187 RHat: 1.001 | Mode: -0.022 95% CI: [-0.042,0.01] BF: 0.521 ROPE Bound: 0.002 In ROPE: 0.046 BF in ROPE: 1.866 RHat: 1 |
|  | Family: student Num. obs: 5370 Num. groups: 34 Bayes R2: 0.033 | Family: student Num. obs: 5370 Num. groups: 34 Bayes R2: 0.056 | Family: student Num. obs: 4438 Num. groups: 34 Bayes R2: 0.016 | Family: student Num. obs: 4438 Num. groups: 34 Bayes R2: 0.057 |

##### 4.1.4. Reaction times

*Table S17.* Only in approach decision trials, the single-trial associations between the continuous measure of expectation and the degree of change of the current interbeat interval (IBI) compared to the previous one, for the last heartbeat before and the third one after feedback, after controlling for reaction times (RT). ROPE – region of practical equivalence, BF – Bayes Factor. Cardiac phase is a continuous measure of distance from the last heartbeat before feedback to feedback, scaled by the distance from the last heartbeat before to the first heartbeat after feedback (IBI 1). All values are standardised.

| Parameter | Dependent variable: Change of IBI -1 only in approach trials | Dependent variable: Change of IBI 3 only in approach trials |
| --- | --- | --- |
| Intercept | Mode: 0.029 95% CI: [-0.053,0.068] BF: 0.011 ROPE Bound: 0.019 In ROPE: 0.486 BF in ROPE: 153 RHat: 1.001 | Mode: 0.107 95% CI: [-0.054,0.126] BF: 0.025 ROPE Bound: 0.02 In ROPE: 0.247 BF in ROPE: 49.841 RHat: 1.003 |
| Expectation | Mode: -0.02 95% CI: [-0.039,0.005] BF: 0.647 ROPE Bound: 0.006 In ROPE: 0.165 BF in ROPE: 1.784 RHat: 1 | Mode: 0.02 95% CI: [-0.008,0.037] BF: 0.547 ROPE Bound: 0.006 In ROPE: 0.209 BF in ROPE: 2.205 RHat: 1 |
| Cardiac phase | Mode: 0.053 95% CI: [0.041,0.082] BF: 8880 ROPE Bound: 0.005 In ROPE: 0 BF in ROPE: 0 RHat: 1 | Mode: -0.011 95% CI: [-0.029,0.015] BF: 0.273 ROPE Bound: 0.006 In ROPE: 0.332 BF in ROPE: 4.658 RHat: 1.001 |
| RT | Mode: 0.124 95% CI: [0.109,0.154] BF: 1491420935863 ROPE Bound: 0.004 In ROPE: 0 BF in ROPE: 0 RHat: 1 | Mode: -0.074 95% CI: [-0.076,-0.031] BF: 1456 ROPE Bound: 0.004 In ROPE: 0 BF in ROPE: 0.001 RHat: 1.001 |
| Expectation : Cardiac phase | Mode: 0.011 95% CI: [-0.014,0.028] BF: 0.244 ROPE Bound: 0.002 In ROPE: 0.115 BF in ROPE: 4.297 RHat: 1 | Mode: 0.01 95% CI: [-0.015,0.03] BF: 0.262 ROPE Bound: 0.002 In ROPE: 0.116 BF in ROPE: 4.103 RHat: 1.001 |
| Cardiac phase : RT | Mode: -0.019 95% CI: [-0.033,0.01] BF: 0.344 ROPE Bound: 0.001 In ROPE: 0.052 BF in ROPE: 3.184 RHat: 1.001 | Mode: 0.02 95% CI: [-0.011,0.034] BF: 0.407 ROPE Bound: 0.001 In ROPE: 0.047 BF in ROPE: 2.544 RHat: 1.001 |
|  | Family: student Num. obs: 5370 Num. groups: 34 Bayes R2: 0.049 | Family: student Num. obs: 5370 Num. groups: 34 Bayes R2: 0.059 |

##### 4.1.5. Prior sensitivity

*Table S18.* The single-trial associations between the continuous measure of expectation and the degree of change of the current interbeat interval (IBI) compared to the previous one, for the last heartbeat before and the third one after feedback, with twice less (normal(0,0.1)) or twice more (normal(0,0.03)) informative priors. ROPE – region of practical equivalence, BF – Bayes Factor. Cardiac phase is a continuous measure of distance from the last heartbeat before feedback to feedback, scaled by the distance from the last heartbeat before to the first heartbeat after feedback (IBI 1). All values are standardised.

| Parameter | Dependent variable: Change of IBI -1  Prior: normal(0,0.1) | Dependent variable: Change of IBI 3  Prior: normal(0,0.1) | Dependent variable: Change of IBI -1  Prior: normal(0,0.03) | Dependent variable: Change of IBI 3  Prior: normal(0,0.03) |
| --- | --- | --- | --- | --- |
| Intercept | Mode: 0.05 95% CI: [-0.048,0.062] BF: 0.011 ROPE Bound: 0.018 In ROPE: 0.512 BF in ROPE: 174 RHat: 1.001 | Mode: 0.038 95% CI: [-0.049,0.128] BF: 0.021 ROPE Bound: 0.02 In ROPE: 0.278 BF in ROPE: 63.925 RHat: 1.005 | Mode: -0.01 95% CI: [-0.047,0.063] BF: 0.01 ROPE Bound: 0.018 In ROPE: 0.507 BF in ROPE: 161 RHat: 1.002 | Mode: 0.022 95% CI: [-0.059,0.127] BF: 0.022 ROPE Bound: 0.02 In ROPE: 0.255 BF in ROPE: 59.951 RHat: 1.004 |
| Expectation | Mode: -0.039 95% CI: [-0.049,-0.015] BF: 31.377 ROPE Bound: 0.006 In ROPE: 0 BF in ROPE: 0.047 RHat: 1.001 | Mode: 0.039 95% CI: [0.017,0.053] BF: 59.68 ROPE Bound: 0.006 In ROPE: 0 BF in ROPE: 0.025 RHat: 1.001 | Mode: -0.023 95% CI: [-0.046,-0.013] BF: 174 ROPE Bound: 0.006 In ROPE: 0 BF in ROPE: 0.012 RHat: 1.001 | Mode: 0.039 95% CI: [0.014,0.049] BF: 145 ROPE Bound: 0.006 In ROPE: 0 BF in ROPE: 0.01 RHat: 1.001 |
| Cardiac phase | Mode: 0.046 95% CI: [0.024,0.057] BF: 439 ROPE Bound: 0.005 In ROPE: 0 BF in ROPE: 0.003 RHat: 1.001 | Mode: 0.012 95% CI: [-0.008,0.025] BF: 0.129 ROPE Bound: 0.006 In ROPE: 0.361 BF in ROPE: 10.969 RHat: 1.001 | Mode: 0.041 95% CI: [0.022,0.053] BF: 1402 ROPE Bound: 0.005 In ROPE: 0 BF in ROPE: 0.001 RHat: 1 | Mode: -0.013 95% CI: [-0.008,0.024] BF: 0.391 ROPE Bound: 0.006 In ROPE: 0.385 BF in ROPE: 3.312 RHat: 1.001 |
| Expectation : Cardiac phase | Mode: -0.012 95% CI: [-0.019,0.014] BF: 0.085 ROPE Bound: 0.002 In ROPE: 0.149 BF in ROPE: 13.789 RHat: 1 | Mode: 0.01 95% CI: [-0.019,0.016] BF: 0.089 ROPE Bound: 0.002 In ROPE: 0.157 BF in ROPE: 13.704 RHat: 1.001 | Mode: -0.005 95% CI: [-0.019,0.013] BF: 0.289 ROPE Bound: 0.002 In ROPE: 0.155 BF in ROPE: 3.688 RHat: 1 | Mode: -0.012 95% CI: [-0.018,0.014] BF: 0.285 ROPE Bound: 0.002 In ROPE: 0.177 BF in ROPE: 3.904 RHat: 1.001 |
|  | Family: student Num. obs: 9808 Num. groups: 34 Bayes R2: 0.022 | Family: student Num. obs: 9808 Num. groups: 34 Bayes R2: 0.055 | Family: student Num. obs: 9808 Num. groups: 34 Bayes R2: 0.021 | Family: student Num. obs: 9808 Num. groups: 34 Bayes R2: 0.054 |

##### 4.1.6. HRV outlier removal

*Table S19.* The single-trial associations between the continuous measure of expectation and the degree of change of the current interbeat interval (IBI) compared to the previous one, for the last heartbeat before and the third one after feedback, after removing RMSSD outliers. ROPE – region of practical equivalence, BF – Bayes Factor. Cardiac phase is a continuous measure of distance from the last heartbeat before feedback to feedback, scaled by the distance from the last heartbeat before to the first heartbeat after feedback (IBI 1). All values are standardised. 2 participants were removed due to high RMSSD (more than 3 SD away from the mean across participants).

| Parameter | Dependent variable: Change of IBI -1 | Dependent variable: Change of IBI 3 |
| --- | --- | --- |
| Intercept | Mode: 0.021 95% CI: [-0.061,0.058] BF: 0.01 ROPE Bound: 0.02 In ROPE: 0.547 BF in ROPE: 179 RHat: 1.004 | Mode: 0.048 95% CI: [-0.043,0.118] BF: 0.023 ROPE Bound: 0.022 In ROPE: 0.288 BF in ROPE: 61.064 RHat: 1.003 |
| Expectation | Mode: -0.027 95% CI: [-0.05,-0.014] BF: 40.347 ROPE Bound: 0.006 In ROPE: 0 BF in ROPE: 0.035 RHat: 1 | Mode: 0.041 95% CI: [0.02,0.057] BF: 115 ROPE Bound: 0.007 In ROPE: 0 BF in ROPE: 0.011 RHat: 1 |
| Cardiac phase | Mode: 0.034 95% CI: [0.024,0.058] BF: 1589 ROPE Bound: 0.006 In ROPE: 0 BF in ROPE: 0.001 RHat: 1 | Mode: 0.02 95% CI: [-0.008,0.029] BF: 0.351 ROPE Bound: 0.006 In ROPE: 0.3 BF in ROPE: 3.619 RHat: 1.001 |
| Expectation : Cardiac phase | Mode: -0.005 95% CI: [-0.02,0.014] BF: 0.192 ROPE Bound: 0.002 In ROPE: 0.152 BF in ROPE: 5.856 RHat: 1.001 | Mode: -0.013 95% CI: [-0.016,0.02] BF: 0.188 ROPE Bound: 0.002 In ROPE: 0.166 BF in ROPE: 5.88 RHat: 1 |
|  | Family: student Num. obs: 9262 Num. groups: 32 Bayes R2: 0.024 | Family: student Num. obs: 9262 Num. groups: 32 Bayes R2: 0.046 |

#### 4.2. Signed PE and outcome valence

##### 4.2.1. Average IBI or RMSSD during rest

*Table S20.* The single-trial associations between the valence of outcome (contrast: lose compared to win) or prediction error (PE) and the degree of change of the current interbeat interval (IBI) compared to the previous one, for the second heartbeat after feedback, controlling for the resting average IBI or resting RMSSD. ROPE – region of practical equivalence, BF – Bayes Factor. Cardiac phase is a continuous measure of distance from the last heartbeat before feedback to feedback, scaled by the distance from the last heartbeat before to the first heartbeat after feedback (IBI 1). All values are standardised.

| Parameter | Dependent variable: Change of IBI 2 | Dependent variable: Change of IBI 2 | Dependent variable: Change of IBI 2 | Dependent variable: Change of IBI 2 |
| --- | --- | --- | --- | --- |
| Intercept | Mode: 0.018 95% CI: [-0.037,0.06] BF: 0.01 ROPE Bound: 0.019 In ROPE: 0.538 BF in ROPE: 184 RHat: 1.002 | Mode: 0.018 95% CI: [-0.038,0.06] BF: 0.01 ROPE Bound: 0.019 In ROPE: 0.542 BF in ROPE: 205 RHat: 1.002 | Mode: 0.026 95% CI: [-0.02,0.076] BF: 0.019 ROPE Bound: 0.019 In ROPE: 0.307 BF in ROPE: 77.167 RHat: 1.004 | Mode: 0.056 95% CI: [-0.018,0.08] BF: 0.02 ROPE Bound: 0.019 In ROPE: 0.304 BF in ROPE: 75.501 RHat: 1.005 |
| Outcome Lose | Mode: 0.05 95% CI: [0.01,0.07] BF: 10.956 ROPE Bound: 0.019 In ROPE: 0.054 BF in ROPE: 0.198 RHat: 1.001 | Mode: 0.032 95% CI: [0.013,0.074] BF: 12.821 ROPE Bound: 0.019 In ROPE: 0.048 BF in ROPE: 0.168 RHat: 1 |  |  |
| Cardiac phase | Mode: 0.132 95% CI: [0.111,0.153] BF: 6282241258076190 ROPE Bound: 0.005 In ROPE: 0 BF in ROPE: 0 RHat: 1.001 | Mode: 0.141 95% CI: [0.118,0.159] BF: 15532985500587458 ROPE Bound: 0.005 In ROPE: 0 BF in ROPE: 0 RHat: 1 | Mode: 0.113 95% CI: [0.101,0.133] BF: 89189465335911 ROPE Bound: 0.005 In ROPE: 0 BF in ROPE: 0 RHat: 1 | Mode: 0.125 95% CI: [0.107,0.139] BF: 26337975654513336 ROPE Bound: 0.005 In ROPE: 0 BF in ROPE: 0 RHat: 1 |
| Rest Average IBI | Mode: -0.075 95% CI: [-0.114,-0.027] BF: 28.466 ROPE Bound: 0.004 In ROPE: 0 BF in ROPE: 0.032 RHat: 1.001 |  | Mode: -0.064 95% CI: [-0.112,-0.026] BF: 57.317 ROPE Bound: 0.004 In ROPE: 0 BF in ROPE: 0.017 RHat: 1.001 |  |
| Outcome Lose : Cardiac phase | Mode: -0.041 95% CI: [-0.064,-0.004] BF: 3.259 ROPE Bound: 0.005 In ROPE: 0.01 BF in ROPE: 0.34 RHat: 1.001 | Mode: -0.032 95% CI: [-0.065,-0.005] BF: 4.29 ROPE Bound: 0.005 In ROPE: 0.003 BF in ROPE: 0.241 RHat: 1 |  |  |
| Cardiac phase : Rest Average IBI | Mode: 0.017 95% CI: [0.006,0.04] BF: 4.785 ROPE Bound: 0.001 In ROPE: 0 BF in ROPE: 0.206 RHat: 1.001 |  | Mode: 0.026 95% CI: [0.007,0.042] BF: 5.61 ROPE Bound: 0.001 In ROPE: 0 BF in ROPE: 0.177 RHat: 1 |  |
| Rest RMSSD |  | Mode: -0.104 95% CI: [-0.115,-0.029] BF: 135 ROPE Bound: 0.003 In ROPE: 0 BF in ROPE: 0.007 RHat: 1.002 |  | Mode: -0.105 95% CI: [-0.116,-0.029] BF: 51.502 ROPE Bound: 0.003 In ROPE: 0 BF in ROPE: 0.018 RHat: 1.001 |
| Cardiac phase : Rest RMSSD |  | Mode: 0.048 95% CI: [0.02,0.065] BF: 123 ROPE Bound: 0.001 In ROPE: 0 BF in ROPE: 0.009 RHat: 1 |  | Mode: 0.04 95% CI: [0.021,0.065] BF: 148 ROPE Bound: 0.001 In ROPE: 0 BF in ROPE: 0.007 RHat: 1 |
| PE |  |  | Mode: -0.017 95% CI: [-0.04,-0.009] BF: 19.282 ROPE Bound: 0.005 In ROPE: 0 BF in ROPE: 0.068 RHat: 1.001 | Mode: -0.021 95% CI: [-0.04,-0.009] BF: 12.791 ROPE Bound: 0.005 In ROPE: 0 BF in ROPE: 0.102 RHat: 1 |
| PE : Cardiac phase |  |  | Mode: 0.02 95% CI: [0.011,0.042] BF: 30.152 ROPE Bound: 0.001 In ROPE: 0 BF in ROPE: 0.033 RHat: 1 | Mode: 0.037 95% CI: [0.011,0.042] BF: 35.21 ROPE Bound: 0.001 In ROPE: 0 BF in ROPE: 0.03 RHat: 1.001 |
|  | Family: student Num. obs: 9808 Num. groups: 34 Bayes R2: 0.037 | Family: student Num. obs: 9808 Num. groups: 34 Bayes R2: 0.04 | Family: student Num. obs: 9808 Num. groups: 34 Bayes R2: 0.037 | Family: student Num. obs: 9808 Num. groups: 34 Bayes R2: 0.041 |

##### 4.2.2. Average IBI or RMSSD within a trial, before or after feedback

*Table S21.* The single-trial associations between the valence of outcome (contrast: lose compared to win) or prediction error (PE) and the degree of change of the current interbeat interval (IBI) compared to the previous one, for the second heartbeat after feedback, controlling for the average IBI or RMSSD before feedback. ROPE – region of practical equivalence, BF – Bayes Factor. Cardiac phase is a continuous measure of distance from the last heartbeat before feedback to feedback, scaled by the distance from the last heartbeat before to the first heartbeat after feedback (IBI 1). All values are standardised.

| Parameter | Dependent variable: Change of IBI 2 | Dependent variable: Change of IBI 2 | Dependent variable: Change of IBI 2 | Dependent variable: Change of IBI 2 |
| --- | --- | --- | --- | --- |
| Intercept | Mode: -0.015 95% CI: [-0.071,0.077] BF: 0.013 ROPE Bound: 0.019 In ROPE: 0.404 BF in ROPE: 105 RHat: 1.006 | Mode: 0.028 95% CI: [-0.044,0.057] BF: 0.01 ROPE Bound: 0.019 In ROPE: 0.543 BF in ROPE: 199 RHat: 1.003 | Mode: -0.039 95% CI: [-0.05,0.091] BF: 0.016 ROPE Bound: 0.019 In ROPE: 0.347 BF in ROPE: 93.835 RHat: 1.006 | Mode: -0.015 95% CI: [-0.023,0.071] BF: 0.013 ROPE Bound: 0.019 In ROPE: 0.368 BF in ROPE: 101 RHat: 1.003 |
| Outcome Lose | Mode: 0.039 95% CI: [0.017,0.076] BF: 31.58 ROPE Bound: 0.019 In ROPE: 0.003 BF in ROPE: 0.068 RHat: 1 | Mode: 0.058 95% CI: [0.012,0.072] BF: 11.981 ROPE Bound: 0.019 In ROPE: 0.049 BF in ROPE: 0.193 RHat: 1 |  |  |
| Cardiac phase | Mode: 0.129 95% CI: [0.109,0.148] BF: 22532636006735 ROPE Bound: 0.005 In ROPE: 0 BF in ROPE: 0 RHat: 1.001 | Mode: 0.138 95% CI: [0.118,0.159] BF: 1.02711404343304e+31 ROPE Bound: 0.005 In ROPE: 0 BF in ROPE: 0 RHat: 1 | Mode: 0.12 95% CI: [0.1,0.13] BF: 45894251164908 ROPE Bound: 0.005 In ROPE: 0 BF in ROPE: 0 RHat: 1.003 | Mode: 0.127 95% CI: [0.108,0.14] BF: 72202668450214944 ROPE Bound: 0.005 In ROPE: 0 BF in ROPE: 0 RHat: 1 |
| Average IBI before feedback | Mode: -0.311 95% CI: [-0.346,-0.288] BF: 1.53411009240211e+23 ROPE Bound: 0.001 In ROPE: 0 BF in ROPE: 0 RHat: 1.001 |  | Mode: -0.315 95% CI: [-0.348,-0.289] BF: 7.68912826207902e+21 ROPE Bound: 0.001 In ROPE: 0 BF in ROPE: 0 RHat: 1 |  |
| Outcome Lose : Cardiac phase | Mode: -0.028 95% CI: [-0.059,-0.001] BF: 2.344 ROPE Bound: 0.005 In ROPE: 0.025 BF in ROPE: 0.439 RHat: 1.001 | Mode: -0.034 95% CI: [-0.061,0] BF: 2.807 ROPE Bound: 0.005 In ROPE: 0.022 BF in ROPE: 0.362 RHat: 1 |  |  |
| Cardiac phase : Average IBI before feedback | Mode: 0.02 95% CI: [0.008,0.043] BF: 9.24 ROPE Bound: 0 In ROPE: 0 BF in ROPE: 0.106 RHat: 1 |  | Mode: 0.034 95% CI: [0.009,0.045] BF: 10.147 ROPE Bound: 0 In ROPE: 0 BF in ROPE: 0.096 RHat: 1 |  |
| RMSSD before feedback |  | Mode: -0.106 95% CI: [-0.137,-0.074] BF: 144284 ROPE Bound: 0 In ROPE: 0 BF in ROPE: 0 RHat: 1 |  | Mode: -0.101 95% CI: [-0.137,-0.075] BF: 95684 ROPE Bound: 0 In ROPE: 0 BF in ROPE: 0 RHat: 1 |
| Cardiac phase : RMSSD before feedback |  | Mode: 0.068 95% CI: [0.043,0.093] BF: 5571 ROPE Bound: 0 In ROPE: 0 BF in ROPE: 0 RHat: 1 |  | Mode: 0.06 95% CI: [0.044,0.095] BF: 23727 ROPE Bound: 0 In ROPE: 0 BF in ROPE: 0 RHat: 1 |
| PE |  |  | Mode: -0.026 95% CI: [-0.044,-0.013] BF: 108 ROPE Bound: 0.005 In ROPE: 0 BF in ROPE: 0.014 RHat: 1 | Mode: -0.016 95% CI: [-0.041,-0.01] BF: 18.221 ROPE Bound: 0.005 In ROPE: 0 BF in ROPE: 0.076 RHat: 1 |
| PE : Cardiac phase |  |  | Mode: 0.031 95% CI: [0.011,0.042] BF: 32.391 ROPE Bound: 0.001 In ROPE: 0 BF in ROPE: 0.032 RHat: 1 | Mode: 0.023 95% CI: [0.011,0.041] BF: 21.844 ROPE Bound: 0.001 In ROPE: 0 BF in ROPE: 0.047 RHat: 1 |
|  | Family: student Num. obs: 9808 Num. groups: 34 Bayes R2: 0.085 | Family: student Num. obs: 9808 Num. groups: 34 Bayes R2: 0.045 | Family: student Num. obs: 9808 Num. groups: 34 Bayes R2: 0.086 | Family: student Num. obs: 9808 Num. groups: 34 Bayes R2: 0.045 |

*Table S22.* The single-trial associations between the valence of outcome (contrast: lose compared to win) or prediction error (PE) and the degree of change of the current interbeat interval (IBI) compared to the previous one, for the second heartbeat after feedback, controlling for the average IBI or RMSSD after feedback. ROPE – region of practical equivalence, BF – Bayes Factor. Cardiac phase is a continuous measure of distance from the last heartbeat before feedback to feedback, scaled by the distance from the last heartbeat before to the first heartbeat after feedback (IBI 1). All values are standardised.

| Parameter | Dependent variable: Change of IBI 2 | Dependent variable: Change of IBI 2 | Dependent variable: Change of IBI 2 | Dependent variable: Change of IBI 2 |
| --- | --- | --- | --- | --- |
| Intercept | Mode: 0.021 95% CI: [-0.053,0.074] BF: 0.012 ROPE Bound: 0.019 In ROPE: 0.435 BF in ROPE: 129 RHat: 1.007 | Mode: 0.022 95% CI: [-0.066,0.04] BF: 0.011 ROPE Bound: 0.019 In ROPE: 0.485 BF in ROPE: 157 RHat: 1.002 | Mode: 0.014 95% CI: [-0.035,0.091] BF: 0.017 ROPE Bound: 0.019 In ROPE: 0.324 BF in ROPE: 81.65 RHat: 1.008 | Mode: -0.022 95% CI: [-0.047,0.06] BF: 0.01 ROPE Bound: 0.019 In ROPE: 0.531 BF in ROPE: 185 RHat: 1.001 |
| Outcome Lose | Mode: 0.044 95% CI: [0.009,0.069] BF: 8.423 ROPE Bound: 0.019 In ROPE: 0.066 BF in ROPE: 0.221 RHat: 1 | Mode: 0.031 95% CI: [0.014,0.073] BF: 19.013 ROPE Bound: 0.019 In ROPE: 0.027 BF in ROPE: 0.124 RHat: 1.001 |  |  |
| Cardiac phase | Mode: 0.135 95% CI: [0.111,0.153] BF: 10868614644768 ROPE Bound: 0.005 In ROPE: 0 BF in ROPE: 0 RHat: 1 | Mode: 0.209 95% CI: [0.19,0.232] BF: 3.04453636968741e+24 ROPE Bound: 0.005 In ROPE: 0 BF in ROPE: 0 RHat: 1.001 | Mode: 0.12 95% CI: [0.101,0.133] BF: 3707220484654 ROPE Bound: 0.005 In ROPE: 0 BF in ROPE: 0 RHat: 1 | Mode: 0.201 95% CI: [0.179,0.214] BF: 8.39202960540127e+24 ROPE Bound: 0.005 In ROPE: 0 BF in ROPE: 0 RHat: 1 |
| Average IBI after feedback | Mode: 0.024 95% CI: [0.001,0.057] BF: 2.32 ROPE Bound: 0.002 In ROPE: 0.005 BF in ROPE: 0.433 RHat: 1.001 |  | Mode: 0.03 95% CI: [-0.001,0.058] BF: 1.863 ROPE Bound: 0.002 In ROPE: 0.012 BF in ROPE: 0.52 RHat: 1.001 |  |
| Outcome Lose : Cardiac phase | Mode: -0.043 95% CI: [-0.065,-0.005] BF: 3.999 ROPE Bound: 0.005 In ROPE: 0.005 BF in ROPE: 0.252 RHat: 1 | Mode: -0.019 95% CI: [-0.06,-0.001] BF: 2.296 ROPE Bound: 0.005 In ROPE: 0.022 BF in ROPE: 0.4 RHat: 1 |  |  |
| Cardiac phase : Average IBI after feedback | Mode: 0.04 95% CI: [0.013,0.048] BF: 49.689 ROPE Bound: 0 In ROPE: 0 BF in ROPE: 0.018 RHat: 1 |  | Mode: 0.039 95% CI: [0.015,0.05] BF: 83.632 ROPE Bound: 0 In ROPE: 0 BF in ROPE: 0.012 RHat: 1 |  |
| RMSSD after feedback |  | Mode: -0.07 95% CI: [-0.093,-0.036] BF: 486 ROPE Bound: 0.001 In ROPE: 0 BF in ROPE: 0.002 RHat: 1 |  | Mode: -0.06 95% CI: [-0.092,-0.035] BF: 746 ROPE Bound: 0.001 In ROPE: 0 BF in ROPE: 0.001 RHat: 1.001 |
| Cardiac phase : RMSSD after feedback |  | Mode: 0.24 95% CI: [0.217,0.265] BF: 6.97263823322337e+22 ROPE Bound: 0 In ROPE: 0 BF in ROPE: 0 RHat: 1 |  | Mode: 0.234 95% CI: [0.218,0.265] BF: 8.4735137495242e+23 ROPE Bound: 0 In ROPE: 0 BF in ROPE: 0 RHat: 1 |
| PE |  |  | Mode: -0.029 95% CI: [-0.04,-0.008] BF: 15.051 ROPE Bound: 0.005 In ROPE: 0 BF in ROPE: 0.099 RHat: 1.001 | Mode: -0.027 95% CI: [-0.04,-0.009] BF: 17.102 ROPE Bound: 0.005 In ROPE: 0 BF in ROPE: 0.072 RHat: 1 |
| PE : Cardiac phase |  |  | Mode: 0.028 95% CI: [0.011,0.043] BF: 35.372 ROPE Bound: 0.001 In ROPE: 0 BF in ROPE: 0.028 RHat: 1.001 | Mode: 0.021 95% CI: [0.01,0.04] BF: 18.14 ROPE Bound: 0.001 In ROPE: 0 BF in ROPE: 0.054 RHat: 1 |
|  | Family: student Num. obs: 9808 Num. groups: 34 Bayes R2: 0.037 | Family: student Num. obs: 9808 Num. groups: 34 Bayes R2: 0.119 | Family: student Num. obs: 9808 Num. groups: 34 Bayes R2: 0.038 | Family: student Num. obs: 9808 Num. groups: 34 Bayes R2: 0.12 |

##### 4.2.3. Number of a trial in a block and approach/avoid decisions

*Table S23.* The single-trial associations between the valence of outcome (contrast: lose compared to win) or prediction error (PE) and the degree of change of the current interbeat interval (IBI) compared to the previous one, for the second heartbeat after feedback, controlling for the number of trial in a block (learning effects) or approach/avoid decision trials. ROPE – region of practical equivalence, BF – Bayes Factor. Cardiac phase is a continuous measure of distance from the last heartbeat before feedback to feedback, scaled by the distance from the last heartbeat before to the first heartbeat after feedback (IBI 1). All values are standardised.

| Parameter | Dependent variable: Change of IBI 2 | Dependent variable: Change of IBI 2 | Dependent variable: Change of IBI 2 | Dependent variable: Change of IBI 2 |
| --- | --- | --- | --- | --- |
| Intercept | Mode: 0.045 95% CI: [-0.048,0.069] BF: 0.01 ROPE Bound: 0.019 In ROPE: 0.482 BF in ROPE: 158 RHat: 1.001 | Mode: 0.02 95% CI: [-0.026,0.084] BF: 0.018 ROPE Bound: 0.019 In ROPE: 0.334 BF in ROPE: 84.587 RHat: 1.004 | Mode: -0.023 95% CI: [-0.078,0.039] BF: 0.013 ROPE Bound: 0.019 In ROPE: 0.391 BF in ROPE: 113 RHat: 1.003 | Mode: -0.019 95% CI: [-0.06,0.057] BF: 0.011 ROPE Bound: 0.019 In ROPE: 0.496 BF in ROPE: 165 RHat: 1.002 |
| Outcome Lose | Mode: 0.037 95% CI: [0.011,0.072] BF: 8.767 ROPE Bound: 0.019 In ROPE: 0.059 BF in ROPE: 0.218 RHat: 1.002 |  | Mode: 0.033 95% CI: [0.01,0.071] BF: 9.833 ROPE Bound: 0.019 In ROPE: 0.051 BF in ROPE: 0.192 RHat: 1.001 |  |
| Cardiac phase | Mode: 0.128 95% CI: [0.108,0.149] BF: 439953568723 ROPE Bound: 0.005 In ROPE: 0 BF in ROPE: 0 RHat: 1 | Mode: 0.12 95% CI: [0.099,0.131] BF: 19786950357955200 ROPE Bound: 0.005 In ROPE: 0 BF in ROPE: 0 RHat: 1.001 | Mode: 0.127 95% CI: [0.107,0.155] BF: 71333287124 ROPE Bound: 0.005 In ROPE: 0 BF in ROPE: 0 RHat: 1.001 | Mode: 0.121 95% CI: [0.098,0.139] BF: 115871196074 ROPE Bound: 0.005 In ROPE: 0 BF in ROPE: 0 RHat: 1.001 |
| Trial in Block | Mode: 0.02 95% CI: [0.018,0.049] BF: 656 ROPE Bound: 0.005 In ROPE: 0 BF in ROPE: 0.003 RHat: 1 | Mode: 0.041 95% CI: [0.017,0.049] BF: 287 ROPE Bound: 0.005 In ROPE: 0 BF in ROPE: 0.005 RHat: 1 |  |  |
| Outcome Lose : Cardiac phase | Mode: -0.049 95% CI: [-0.063,-0.002] BF: 2.574 ROPE Bound: 0.005 In ROPE: 0.019 BF in ROPE: 0.383 RHat: 1 |  | Mode: -0.032 95% CI: [-0.063,-0.002] BF: 3.036 ROPE Bound: 0.005 In ROPE: 0.018 BF in ROPE: 0.354 RHat: 1 |  |
| Cardiac phase : Trial in Block | Mode: -0.015 95% CI: [-0.021,0.01] BF: 0.2 ROPE Bound: 0.002 In ROPE: 0.138 BF in ROPE: 5.497 RHat: 1.002 | Mode: -0.009 95% CI: [-0.021,0.01] BF: 0.2 ROPE Bound: 0.002 In ROPE: 0.132 BF in ROPE: 5.509 RHat: 1.001 |  |  |
| PE |  | Mode: -0.02 95% CI: [-0.04,-0.008] BF: 13.109 ROPE Bound: 0.005 In ROPE: 0 BF in ROPE: 0.094 RHat: 1 |  | Mode: -0.015 95% CI: [-0.04,-0.008] BF: 12.875 ROPE Bound: 0.005 In ROPE: 0 BF in ROPE: 0.104 RHat: 1.002 |
| PE : Cardiac phase |  | Mode: 0.027 95% CI: [0.009,0.042] BF: 12.313 ROPE Bound: 0.001 In ROPE: 0 BF in ROPE: 0.078 RHat: 1 |  | Mode: 0.018 95% CI: [0.009,0.04] BF: 16.646 ROPE Bound: 0.001 In ROPE: 0 BF in ROPE: 0.073 RHat: 1.001 |
| Decision Avoid |  |  | Mode: 0.041 95% CI: [0.038,0.099] BF: 2733 ROPE Bound: 0.019 In ROPE: 0 BF in ROPE: 0.002 RHat: 1 | Mode: 0.066 95% CI: [0.037,0.097] BF: 1611 ROPE Bound: 0.019 In ROPE: 0 BF in ROPE: 0.002 RHat: 1.001 |
| Cardiac phase : Decision Avoid |  |  | Mode: -0.016 95% CI: [-0.04,0.02] BF: 0.347 ROPE Bound: 0.005 In ROPE: 0.251 BF in ROPE: 3.553 RHat: 1.001 | Mode: -0.019 95% CI: [-0.039,0.02] BF: 0.353 ROPE Bound: 0.005 In ROPE: 0.253 BF in ROPE: 3.284 RHat: 1.001 |
|  | Family: student Num. obs: 9808 Num. groups: 34 Bayes R2: 0.036 | Family: student Num. obs: 9808 Num. groups: 34 Bayes R2: 0.037 | Family: student Num. obs: 9808 Num. groups: 34 Bayes R2: 0.036 | Family: student Num. obs: 9808 Num. groups: 34 Bayes R2: 0.036 |

##### 4.2.4. Reaction times

*Table S24.* Only in approach decision trials, the single-trial associations between the valence of outcome (contrast: lose compared to win), prediction error (PE) or unsigned PE and the degree of change of the current interbeat interval (IBI) compared to the previous one, for the second heartbeat after feedback, controlling for the reaction time (RT). ROPE – region of practical equivalence, BF – Bayes Factor. Cardiac phase is a continuous measure of distance from the last heartbeat before feedback to feedback, scaled by the distance from the last heartbeat before to the first heartbeat after feedback (IBI 1). All values are standardised.

| Parameter | Dependent variable: Change of IBI 2 only in approach trials | Dependent variable: Change of IBI 2 only in approach trials | Dependent variable: Change of IBI 2 only in approach trials |
| --- | --- | --- | --- |
| Intercept | Mode: -0.016 95% CI: [-0.094,0.046] BF: 0.015 ROPE Bound: 0.018 In ROPE: 0.346 BF in ROPE: 90.795 RHat: 1.002 | Mode: -0.037 95% CI: [-0.066,0.067] BF: 0.012 ROPE Bound: 0.018 In ROPE: 0.438 BF in ROPE: 141 RHat: 1.002 | Mode: -0.024 95% CI: [-0.067,0.065] BF: 0.012 ROPE Bound: 0.018 In ROPE: 0.436 BF in ROPE: 129 RHat: 1.012 |
| Outcome Lose | Mode: 0.04 95% CI: [0,0.079] BF: 2.886 ROPE Bound: 0.018 In ROPE: 0.129 BF in ROPE: 0.398 RHat: 1 |  |  |
| Cardiac phase | Mode: 0.138 95% CI: [0.1,0.155] BF: 681106648 ROPE Bound: 0.005 In ROPE: 0 BF in ROPE: 0 RHat: 1 | Mode: 0.118 95% CI: [0.097,0.14] BF: 24534660706 ROPE Bound: 0.005 In ROPE: 0 BF in ROPE: 0 RHat: 1 | Mode: 0.119 95% CI: [0.1,0.141] BF: 279397636531 ROPE Bound: 0.005 In ROPE: 0 BF in ROPE: 0 RHat: 1 |
| RT | Mode: -0.071 95% CI: [-0.097,-0.053] BF: 103962 ROPE Bound: 0.004 In ROPE: 0 BF in ROPE: 0 RHat: 1.001 | Mode: -0.077 95% CI: [-0.096,-0.052] BF: 253367 ROPE Bound: 0.004 In ROPE: 0 BF in ROPE: 0 RHat: 1.001 | Mode: -0.076 95% CI: [-0.095,-0.052] BF: 3768625 ROPE Bound: 0.004 In ROPE: 0 BF in ROPE: 0 RHat: 1 |
| Outcome Lose : Cardiac phase | Mode: -0.022 95% CI: [-0.058,0.021] BF: 0.606 ROPE Bound: 0.005 In ROPE: 0.149 BF in ROPE: 1.702 RHat: 1 |  |  |
| Cardiac phase : RT | Mode: -0.017 95% CI: [-0.033,0.009] BF: 0.422 ROPE Bound: 0.001 In ROPE: 0.039 BF in ROPE: 2.482 RHat: 1 | Mode: -0.011 95% CI: [-0.035,0.007] BF: 0.459 ROPE Bound: 0.001 In ROPE: 0.038 BF in ROPE: 2.18 RHat: 1.001 | Mode: -0.013 95% CI: [-0.035,0.008] BF: 0.461 ROPE Bound: 0.001 In ROPE: 0.035 BF in ROPE: 2.295 RHat: 1.001 |
| PE |  | Mode: -0.014 95% CI: [-0.044,-0.002] BF: 2.138 ROPE Bound: 0.005 In ROPE: 0.024 BF in ROPE: 0.501 RHat: 1 |  |
| PE : Cardiac phase |  | Mode: 0.028 95% CI: [0.008,0.051] BF: 7.185 ROPE Bound: 0.001 In ROPE: 0 BF in ROPE: 0.134 RHat: 1.002 |  |
| Unsigned PE |  |  | Mode: 0.027 95% CI: [0.008,0.05] BF: 9.075 ROPE Bound: 0.006 In ROPE: 0 BF in ROPE: 0.138 RHat: 1.001 |
| Unsigned PE : Cardiac phase |  |  | Mode: -0.027 95% CI: [-0.047,-0.004] BF: 3.19 ROPE Bound: 0.002 In ROPE: 0 BF in ROPE: 0.315 RHat: 1.001 |
|  | Family: student Num. obs: 5370 Num. groups: 34 Bayes R2: 0.046 | Family: student Num. obs: 5370 Num. groups: 34 Bayes R2: 0.047 | Family: student Num. obs: 5370 Num. groups: 34 Bayes R2: 0.047 |

##### 4.2.5. Prior sensitivity

*Table S25.* The single-trial associations between the valence of outcome (contrast: lose compared to win) or prediction error (PE) and the degree of change of the current interbeat interval (IBI) compared to the previous one, for the second heartbeat after feedback, with twice less (normal(0,0.1)) or twice more (normal(0,0.03)) informative priors. ROPE – region of practical equivalence, BF – Bayes Factor. Cardiac phase is a continuous measure of distance from the last heartbeat before feedback to feedback, scaled by the distance from the last heartbeat before to the first heartbeat after feedback (IBI 1). All values are standardised.

| Parameter | Dependent variable: Change of IBI 2  Prior: normal(0,0.1) | Dependent variable: Change of IBI 2  Prior: normal(0,0.1) | Dependent variable: Change of IBI 2  Prior: normal(0,0.03) | Dependent variable: Change of IBI 2  Prior: normal(0,0.03) |
| --- | --- | --- | --- | --- |
| Intercept | Mode: 0.018 95% CI: [-0.053,0.064] BF: 0.011 ROPE Bound: 0.019 In ROPE: 0.48 BF in ROPE: 147 RHat: 1.003 | Mode: 0.045 95% CI: [-0.03,0.084] BF: 0.018 ROPE Bound: 0.019 In ROPE: 0.322 BF in ROPE: 80.736 RHat: 1.002 | Mode: 0.015 95% CI: [-0.047,0.069] BF: 0.012 ROPE Bound: 0.019 In ROPE: 0.457 BF in ROPE: 148 RHat: 1.004 | Mode: 0.072 95% CI: [-0.023,0.09] BF: 0.02 ROPE Bound: 0.019 In ROPE: 0.311 BF in ROPE: 75.077 RHat: 1.003 |
| Outcome Lose | Mode: 0.05 95% CI: [0.014,0.078] BF: 6.93 ROPE Bound: 0.019 In ROPE: 0.036 BF in ROPE: 0.343 RHat: 1 |  | Mode: 0.038 95% CI: [0.006,0.063] BF: 9.488 ROPE Bound: 0.019 In ROPE: 0.106 BF in ROPE: 0.169 RHat: 1 |  |
| Cardiac phase | Mode: 0.124 95% CI: [0.112,0.155] BF: 284566289092785 ROPE Bound: 0.005 In ROPE: 0 BF in ROPE: 0 RHat: 1.001 | Mode: 0.119 95% CI: [0.101,0.132] BF: 3642794137591 ROPE Bound: 0.005 In ROPE: 0 BF in ROPE: 0 RHat: 1 | Mode: 0.129 95% CI: [0.098,0.137] BF: 4900983897166 ROPE Bound: 0.005 In ROPE: 0 BF in ROPE: 0 RHat: 1.001 | Mode: 0.11 95% CI: [0.094,0.124] BF: 146260456626443232 ROPE Bound: 0.005 In ROPE: 0 BF in ROPE: 0 RHat: 1 |
| Outcome Lose : Cardiac phase | Mode: -0.046 95% CI: [-0.073,-0.009] BF: 3.285 ROPE Bound: 0.005 In ROPE: 0 BF in ROPE: 0.336 RHat: 1.001 |  | Mode: -0.021 95% CI: [-0.049,0.007] BF: 1.345 ROPE Bound: 0.005 In ROPE: 0.116 BF in ROPE: 0.739 RHat: 1.001 |  |
| PE |  | Mode: -0.018 95% CI: [-0.041,-0.01] BF: 12.669 ROPE Bound: 0.005 In ROPE: 0 BF in ROPE: 0.132 RHat: 1.001 |  | Mode: -0.028 95% CI: [-0.039,-0.008] BF: 21.665 ROPE Bound: 0.005 In ROPE: 0 BF in ROPE: 0.058 RHat: 1 |
| PE : Cardiac phase |  | Mode: 0.029 95% CI: [0.01,0.041] BF: 19.788 ROPE Bound: 0.001 In ROPE: 0 BF in ROPE: 0.05 RHat: 1 |  | Mode: 0.027 95% CI: [0.009,0.039] BF: 27.997 ROPE Bound: 0.001 In ROPE: 0 BF in ROPE: 0.034 RHat: 1.001 |
|  | Family: student Num. obs: 9808 Num. groups: 34 Bayes R2: 0.035 | Family: student Num. obs: 9808 Num. groups: 34 Bayes R2: 0.036 | Family: student Num. obs: 9808 Num. groups: 34 Bayes R2: 0.033 | Family: student Num. obs: 9808 Num. groups: 34 Bayes R2: 0.034 |

##### 4.2.6. HRV outlier removal

*Table S26.* The single-trial associations between the valence of outcome (contrast: lose compared to win) or prediction error (PE) and the degree of change of the current interbeat interval (IBI) compared to the previous one, for the second heartbeat after feedback, after removing RMSSD outliers. ROPE – region of practical equivalence, BF – Bayes Factor. Cardiac phase is a continuous measure of distance from the last heartbeat before feedback to feedback, scaled by the distance from the last heartbeat before to the first heartbeat after feedback (IBI 1). All values are standardised.

| Parameter | Dependent variable: Change of IBI 2 | Dependent variable: Change of IBI 2 |
| --- | --- | --- |
| Intercept | Mode: 0.015 95% CI: [-0.048,0.067] BF: 0.01 ROPE Bound: 0.021 In ROPE: 0.537 BF in ROPE: 171 RHat: 1.002 | Mode: 0.015 95% CI: [-0.025,0.081] BF: 0.019 ROPE Bound: 0.021 In ROPE: 0.354 BF in ROPE: 86.086 RHat: 1.002 |
| Outcome Lose | Mode: 0.059 95% CI: [0.012,0.079] BF: 15.476 ROPE Bound: 0.021 In ROPE: 0.044 BF in ROPE: 0.143 RHat: 1 |  |
| Cardiac phase | Mode: 0.149 95% CI: [0.117,0.162] BF: 132754215820 ROPE Bound: 0.006 In ROPE: 0 BF in ROPE: 0 RHat: 1 | Mode: 0.133 95% CI: [0.108,0.143] BF: 38779703599495984 ROPE Bound: 0.006 In ROPE: 0 BF in ROPE: 0 RHat: 1 |
| Outcome Lose : Cardiac phase | Mode: -0.041 95% CI: [-0.063,0.003] BF: 1.519 ROPE Bound: 0.006 In ROPE: 0.055 BF in ROPE: 0.638 RHat: 1 |  |
| PE |  | Mode: -0.024 95% CI: [-0.044,-0.009] BF: 9.757 ROPE Bound: 0.005 In ROPE: 0 BF in ROPE: 0.118 RHat: 1 |
| PE : Cardiac phase |  | Mode: 0.027 95% CI: [0.008,0.043] BF: 8.255 ROPE Bound: 0.002 In ROPE: 0 BF in ROPE: 0.122 RHat: 1 |
|  | Family: student Num. obs: 9262 Num. groups: 32 Bayes R2: 0.035 | Family: student Num. obs: 9262 Num. groups: 32 Bayes R2: 0.036 |

#### 4.3. P3b analyses

##### 4.3.1. Pseudotrial-corrected average responses

*Table S27.* For the first heartbeat after feedback (R1) in systole trials (first 1/3 of the cardiac cycle) the difference between pseudotrial-corrected averaged evoked responses of participants for high and low IBI conditions obtained with a median split within participants for the following channels and time windows: Pz (P3b) from 335 to 385 ms, FCz (p3a) from 353 to 403 ms, FCz (FRN) from 241 to 291 ms, Pz and FCz baseline from -200 to 0 ms. Average ECG amplitudes in the same time windows are supposed to be close to 0 showing effectiveness of pseudotrial control. ROPE – region of practical equivalence, BF – Bayes Factor. P3a and P3b were the main analyses, the rest are sensitivity analyses to show specificity.

| P3b | Pz baseline | ECG P3b | ECG baseline | ECG p3a | ECG FRN | FCz baseline | FCz p3a | FCz FRN |
| --- | --- | --- | --- | --- | --- | --- | --- | --- |
| Mode: 0.679 95% CI: [0.131,0.912] **BF: 4.215** ROPE Bound: 0.1 In ROPE: 0 BF in ROPE: 0.302 RHat: 1.001 | Mode: -0.365 95% CI: [-0.614,-0.033] BF: 1.301 ROPE Bound: 0.1 In ROPE: 0.038 BF in ROPE: 0.916 RHat: 1.001 | Mode: 1.771 95% CI: [-1.131,1.246] BF: 0.114 ROPE Bound: 0.1 In ROPE: 0.126 BF in ROPE: 10.196 RHat: 1 | Mode: -2.957 95% CI: [-4.72,0.081] BF: 0.751 ROPE Bound: 0.1 In ROPE: 0.009 BF in ROPE: 1.283 RHat: 1.001 | Mode: 0.013 95% CI: [-1.1,1.355] BF: 0.104 ROPE Bound: 0.1 In ROPE: 0.125 BF in ROPE: 10.739 RHat: 1.001 | Mode: 4.623 95% CI: [-1.123,9.049] BF: 0.352 ROPE Bound: 0.1 In ROPE: 0.012 BF in ROPE: 2.921 RHat: 1.001 | Mode: -0.484 95% CI: [-0.769,-0.032] BF: 1.096 ROPE Bound: 0.1 In ROPE: 0.038 BF in ROPE: 1.065 RHat: 1 | Mode: -0.019 95% CI: [-0.387,0.438] BF: 0.114 ROPE Bound: 0.1 In ROPE: 0.392 BF in ROPE: 13.129 RHat: 1 | Mode: -0.231 95% CI: [-0.558,0.308] BF: 0.128 ROPE Bound: 0.1 In ROPE: 0.33 BF in ROPE: 11.06 RHat: 1 |
| Family: gaussian Num. obs: 34 | Family: skew normal Num. obs: 34 | Family: student Num. obs: 34 | Family: student Num. obs: 34 | Family: student Num. obs: 34 | Family: gaussian Num. obs: 34 | Family: gaussian Num. obs: 34 | Family: gaussian Num. obs: 34 | Family: gaussian Num. obs: 34 |

*Table S28.* For the first heartbeat after feedback (R1) in diastole trials (last 1/3 of the cardiac cycle) the difference between pseudotrial-corrected averaged evoked responses of participants for high and low IBI conditions obtained with a median split within participants for the following channels and time windows: Pz (P3b) from 421 to 471 ms, FCz (p3a) from 353 to 403 ms, FCz (FRN) from 241 to 291 ms, Pz and FCz baseline from -200 to 0 ms. Average ECG amplitudes in the same time windows are supposed to be close to 0 showing effectiveness of pseudotrial control. ROPE – region of practical equivalence, BF – Bayes Factor. P3a and P3b were the main analyses, the rest are sensitivity analyses to show specificity.

| P3b | Pz baseline | ECG P3b | ECG baseline | ECG p3a | ECG FRN | FCz baseline | FCz p3a | FCz FRN |
| --- | --- | --- | --- | --- | --- | --- | --- | --- |
| Mode: 0.831 95% CI: [-0.143,0.792] BF: 0.258 ROPE Bound: 0.1 In ROPE: 0.156 BF in ROPE: 4.472 RHat: 1.001 | Mode: 0.117 95% CI: [-0.259,0.443] BF: 0.129 ROPE Bound: 0.1 In ROPE: 0.4 BF in ROPE: 11.199 RHat: 1.001 | Mode: 1.283 95% CI: [-5.271,6.103] BF: 0.111 ROPE Bound: 0.1 In ROPE: 0.032 BF in ROPE: 9.408 RHat: 1 | Mode: 0.83 95% CI: [-0.609,1.745] BF: 0.113 ROPE Bound: 0.1 In ROPE: 0.098 BF in ROPE: 9.687 RHat: 1.001 | Mode: -1.832 95% CI: [-7.934,2.939] BF: 0.184 ROPE Bound: 0.1 In ROPE: 0.02 BF in ROPE: 5.855 RHat: 1 | Mode: 1.115 95% CI: [-5.669,4.225] BF: 0.128 ROPE Bound: 0.1 In ROPE: 0.03 BF in ROPE: 8.545 RHat: 1.001 | Mode: -0.171 95% CI: [-0.568,0.132] BF: 0.242 ROPE Bound: 0.1 In ROPE: 0.232 BF in ROPE: 5.218 RHat: 1 | Mode: 0.197 95% CI: [-0.14,0.565] BF: 0.221 ROPE Bound: 0.1 In ROPE: 0.244 BF in ROPE: 5.798 RHat: 1 | Mode: 0.455 95% CI: [-0.72,0.317] BF: 0.161 ROPE Bound: 0.1 In ROPE: 0.243 BF in ROPE: 7.704 RHat: 1 |
| Family: gaussian Num. obs: 34 | Family: gaussian Num. obs: 34 | Family: gaussian Num. obs: 34 | Family: student Num. obs: 34 | Family: gaussian Num. obs: 34 | Family: gaussian Num. obs: 34 | Family: gaussian Num. obs: 34 | Family: gaussian Num. obs: 34 | Family: gaussian Num. obs: 34 |

*Table S29.* For the second heartbeat after feedback (R2) in systole trials (first 1/3 of the cardiac cycle) the difference between pseudotrial-corrected averaged evoked responses of participants for high and low IBI conditions obtained with a median split within participants for the following channels and time windows: Pz (P3b) from 335 to 385 ms, FCz (p3a) from 353 to 403 ms, FCz (FRN) from 241 to 291 ms, Pz and FCz baseline from -200 to 0 ms. Average ECG amplitudes in the same time windows are supposed to be close to 0 showing effectiveness of pseudotrial control. ROPE – region of practical equivalence, BF – Bayes Factor. P3a and P3b were the main analyses, the rest are sensitivity analyses to show specificity.

| P3b | Pz baseline | ECG P3b | ECG baseline | ECG p3a | ECG FRN | FCz baseline | FCz p3a | FCz FRN |
| --- | --- | --- | --- | --- | --- | --- | --- | --- |
| Mode: -0.079 95% CI: [-0.336,0.347] BF: 0.114 ROPE Bound: 0.1 In ROPE: 0.475 BF in ROPE: 14.097 RHat: 1.002 | Mode: 0.384 95% CI: [-0.193,0.445] BF: 0.154 ROPE Bound: 0.1 In ROPE: 0.386 BF in ROPE: 9.325 RHat: 1 | Mode: 1.065 95% CI: [-1.269,1.863] BF: 0.096 ROPE Bound: 0.1 In ROPE: 0.107 BF in ROPE: 11.494 RHat: 1.001 | Mode: 1.16 95% CI: [-2.143,2.342] BF: 0.107 ROPE Bound: 0.1 In ROPE: 0.08 BF in ROPE: 9.749 RHat: 1 | Mode: 1.177 95% CI: [-0.886,2.394] BF: 0.163 ROPE Bound: 0.1 In ROPE: 0.071 BF in ROPE: 6.617 RHat: 1.002 | Mode: -1.829 95% CI: [-5.12,3.128] BF: 0.125 ROPE Bound: 0.1 In ROPE: 0.033 BF in ROPE: 8.495 RHat: 1 | Mode: -0.142 95% CI: [-0.412,0.314] BF: 0.123 ROPE Bound: 0.1 In ROPE: 0.428 BF in ROPE: 12.672 RHat: 1.001 | Mode: -0.382 95% CI: [-1.07,0.089] BF: 0.532 ROPE Bound: 0.1 In ROPE: 0.066 BF in ROPE: 2.014 RHat: 1 | Mode: -0.585 95% CI: [-0.921,0.007] BF: 0.743 ROPE Bound: 0.1 In ROPE: 0.05 BF in ROPE: 1.511 RHat: 1.001 |
| Family: gaussian Num. obs: 34 | Family: gaussian Num. obs: 34 | Family: student Num. obs: 34 | Family: gaussian Num. obs: 34 | Family: student Num. obs: 34 | Family: gaussian Num. obs: 34 | Family: gaussian Num. obs: 34 | Family: gaussian Num. obs: 34 | Family: gaussian Num. obs: 34 |

*Table S30.* For the second heartbeat after feedback (R2) in diastole trials (last 1/3 of the cardiac cycle) the difference between pseudotrial-corrected averaged evoked responses of participants for high and low IBI conditions obtained with a median split within participants for the following channels and time windows: Pz (P3b) from 421 to 471 ms, FCz (p3a) from 353 to 403 ms, FCz (FRN) from 241 to 291 ms, Pz and FCz baseline from -200 to 0 ms. Average ECG amplitudes in the same time windows are supposed to be close to 0 showing effectiveness of pseudotrial control. ROPE – region of practical equivalence, BF – Bayes Factor. P3a and P3b were the main analyses, the rest are sensitivity analyses to show specificity.

| P3b | Pz baseline | ECG P3b | ECG baseline | ECG p3a | ECG FRN | FCz baseline | FCz p3a | FCz FRN |
| --- | --- | --- | --- | --- | --- | --- | --- | --- |
| Mode: 1.057 95% CI: [0.337,1.164] **BF: 29.791** ROPE Bound: 0.1 In ROPE: 0 BF in ROPE: 0.039 RHat: 1 | Mode: 0.134 95% CI: [-0.286,0.44] BF: 0.123 ROPE Bound: 0.1 In ROPE: 0.417 BF in ROPE: 12.174 RHat: 1 | Mode: -1.818 95% CI: [-6.753,4.291] BF: 0.127 ROPE Bound: 0.1 In ROPE: 0.025 BF in ROPE: 8.053 RHat: 1 | Mode: -0.289 95% CI: [-1.43,0.849] BF: 0.09 ROPE Bound: 0.1 In ROPE: 0.132 BF in ROPE: 12.824 RHat: 1 | Mode: -2.106 95% CI: [-7.217,3.771] BF: 0.131 ROPE Bound: 0.1 In ROPE: 0.025 BF in ROPE: 7.894 RHat: 1.001 | Mode: -5.774 95% CI: [-5.979,5.235] BF: 0.103 ROPE Bound: 0.1 In ROPE: 0.029 BF in ROPE: 9.965 RHat: 1.001 | Mode: -0.101 95% CI: [-0.584,0.14] BF: 0.228 ROPE Bound: 0.1 In ROPE: 0.232 BF in ROPE: 5.424 RHat: 1.001 | Mode: 0.598 95% CI: [-0.127,0.86] BF: 0.35 ROPE Bound: 0.1 In ROPE: 0.115 BF in ROPE: 3.181 RHat: 1 | Mode: 0.076 95% CI: [-0.451,0.487] BF: 0.117 ROPE Bound: 0.1 In ROPE: 0.344 BF in ROPE: 11.896 RHat: 1.001 |
| Family: gaussian Num. obs: 34 | Family: skew normal Num. obs: 34 | Family: gaussian Num. obs: 34 | Family: student Num. obs: 34 | Family: student Num. obs: 34 | Family: student Num. obs: 34 | Family: gaussian Num. obs: 34 | Family: gaussian Num. obs: 34 | Family: gaussian Num. obs: 34 |

##### 4.3.2. Control for features of feedback

*Table S31.* Separately only within systole or diastole trials (first and last 1/3 of the cardiac cycle), the single-trial associations between the continuous measure of P3b amplitude and the degree of change in the current interbeat interval (IBI) compared to the previous one, for the first and second heartbeats after feedback, controlling for outcome valence, prediction error (PE), or expectation. ROPE – region of practical equivalence, BF – Bayes Factor. All values are standardised.

| Parameter | Dependent variable: Change of IBI 1 only in systole | Dependent variable: Change of IBI 2 only in diastole | Dependent variable: Change of IBI 1 only in systole | Dependent variable: Change of IBI 2 only in diastole | Dependent variable: Change of IBI 1 only in systole | Dependent variable: Change of IBI 2 only in diastole |
| --- | --- | --- | --- | --- | --- | --- |
| Intercept | Mode: 0.128 95% CI: [0.036,0.164] BF: 0.993 ROPE Bound: 0.018 In ROPE: 0 BF in ROPE: 1.353 RHat: 1.001 | Mode: 0.176 95% CI: [0.116,0.216] BF: 5883 ROPE Bound: 0.019 In ROPE: 0 BF in ROPE: 0 RHat: 1 | Mode: 0.111 95% CI: [0.033,0.16] BF: 0.766 ROPE Bound: 0.018 In ROPE: 0 BF in ROPE: 1.588 RHat: 1.002 | Mode: 0.162 95% CI: [0.115,0.206] BF: 621 ROPE Bound: 0.019 In ROPE: 0 BF in ROPE: 0.002 RHat: 1 | Mode: 0.093 95% CI: [0.033,0.158] BF: 0.866 ROPE Bound: 0.018 In ROPE: 0 BF in ROPE: 1.492 RHat: 1.001 | Mode: 0.139 95% CI: [0.117,0.207] BF: 1256 ROPE Bound: 0.019 In ROPE: 0 BF in ROPE: 0.001 RHat: 1.001 |
| P3b | Mode: 0.012 95% CI: [0.012,0.069] BF: 13.73 ROPE Bound: 0.003 In ROPE: 0 BF in ROPE: 0.072 RHat: 1 | Mode: 0.061 95% CI: [0.027,0.083] BF: 407 ROPE Bound: 0.003 In ROPE: 0 BF in ROPE: 0.003 RHat: 1.001 | Mode: 0.061 95% CI: [0.012,0.068] BF: 16.421 ROPE Bound: 0.003 In ROPE: 0 BF in ROPE: 0.06 RHat: 1.001 | Mode: 0.052 95% CI: [0.029,0.082] BF: 456 ROPE Bound: 0.003 In ROPE: 0 BF in ROPE: 0.002 RHat: 1 | Mode: 0.049 95% CI: [0.012,0.067] BF: 12.468 ROPE Bound: 0.003 In ROPE: 0 BF in ROPE: 0.076 RHat: 1 | Mode: 0.051 95% CI: [0.026,0.081] BF: 241 ROPE Bound: 0.003 In ROPE: 0 BF in ROPE: 0.004 RHat: 1 |
| Outcome Lose | Mode: -0.022 95% CI: [-0.056,0.041] BF: 0.515 ROPE Bound: 0.018 In ROPE: 0.546 BF in ROPE: 2.771 RHat: 1.001 | Mode: -0.011 95% CI: [-0.062,0.033] BF: 0.58 ROPE Bound: 0.019 In ROPE: 0.507 BF in ROPE: 2.374 RHat: 1.001 |  |  |  |  |
| PE |  |  | Mode: 0.013 95% CI: [-0.014,0.039] BF: 0.43 ROPE Bound: 0.005 In ROPE: 0.177 BF in ROPE: 2.66 RHat: 1 | Mode: 0.014 95% CI: [-0.013,0.04] BF: 0.454 ROPE Bound: 0.005 In ROPE: 0.175 BF in ROPE: 2.371 RHat: 1 |  |  |
| Expectation |  |  |  |  | Mode: -0.011 95% CI: [-0.043,0.013] BF: 0.544 ROPE Bound: 0.005 In ROPE: 0.173 BF in ROPE: 2.071 RHat: 1 | Mode: -0.024 95% CI: [-0.039,0.017] BF: 0.36 ROPE Bound: 0.006 In ROPE: 0.238 BF in ROPE: 2.935 RHat: 1.001 |
|  | Family: student Num. obs: 3212 Num. groups: 34 Bayes R2: 0.026 | Family: student Num. obs: 3229 Num. groups: 34 Bayes R2: 0.015 | Family: student Num. obs: 3212 Num. groups: 34 Bayes R2: 0.027 | Family: student Num. obs: 3229 Num. groups: 34 Bayes R2: 0.015 | Family: student Num. obs: 3212 Num. groups: 34 Bayes R2: 0.027 | Family: student Num. obs: 3229 Num. groups: 34 Bayes R2: 0.015 |

##### 4.3.3. Baseline at Pz and average ECG amplitude

*Table S32.* Separately only within systole or diastole trials (first and last 1/3 of the cardiac cycle), the single-trial associations between the continuous measure of P3b amplitude and the degree of change in the current interbeat interval (IBI) compared to the previous one, for the first and second heartbeats after feedback, controlling for baseline amplitudes at Pz channel or for ECG amplitudes in the same time window. ROPE – region of practical equivalence, BF – Bayes Factor. All values are standardised.

| Parameter | Dependent variable: Change of IBI 1 only in systole | Dependent variable: Change of IBI 2 only in diastole | Dependent variable: Change of IBI 1 only in systole | Dependent variable: Change of IBI 2 only in diastole |
| --- | --- | --- | --- | --- |
| Intercept | Mode: 0.102 95% CI: [0.03,0.154] BF: 0.883 ROPE Bound: 0.018 In ROPE: 0 BF in ROPE: 1.546 RHat: 1.002 | Mode: 0.16 95% CI: [0.116,0.208] BF: 3835 ROPE Bound: 0.019 In ROPE: 0 BF in ROPE: 0 RHat: 1 | Mode: 0.016 95% CI: [-0.061,0.088] BF: 0.015 ROPE Bound: 0.018 In ROPE: 0.377 BF in ROPE: 108 RHat: 1.002 | Mode: 0.186 95% CI: [0.132,0.227] BF: 2223 ROPE Bound: 0.019 In ROPE: 0 BF in ROPE: 0.001 RHat: 1 |
| P3b | Mode: 0.036 95% CI: [0.021,0.08] BF: 79.875 ROPE Bound: 0.003 In ROPE: 0 BF in ROPE: 0.012 RHat: 1 | Mode: 0.046 95% CI: [0.031,0.086] BF: 553 ROPE Bound: 0.003 In ROPE: 0 BF in ROPE: 0.002 RHat: 1 | Mode: 0.064 95% CI: [0.01,0.067] BF: 10.863 ROPE Bound: 0.003 In ROPE: 0 BF in ROPE: 0.094 RHat: 1.001 | Mode: 0.052 95% CI: [0.029,0.083] BF: 275 ROPE Bound: 0.003 In ROPE: 0 BF in ROPE: 0.004 RHat: 1 |
| Pz baseline | Mode: -0.039 95% CI: [-0.069,-0.014] BF: 21.125 ROPE Bound: 0.001 In ROPE: 0 BF in ROPE: 0.048 RHat: 1.001 | Mode: -0.025 95% CI: [-0.048,0.008] BF: 0.715 ROPE Bound: 0.001 In ROPE: 0.028 BF in ROPE: 1.389 RHat: 1 |  |  |
| ECG |  |  | Mode: -0.221 95% CI: [-0.256,-0.117] BF: 31462 ROPE Bound: 0.002 In ROPE: 0 BF in ROPE: 0 RHat: 1.001 | Mode: -0.022 95% CI: [-0.052,-0.006] BF: 4.312 ROPE Bound: 0.001 In ROPE: 0 BF in ROPE: 0.235 RHat: 1.001 |
|  | Family: student Num. obs: 3212 Num. groups: 34 Bayes R2: 0.027 | Family: student Num. obs: 3229 Num. groups: 34 Bayes R2: 0.015 | Family: student Num. obs: 3212 Num. groups: 34 Bayes R2: 0.031 | Family: student Num. obs: 3229 Num. groups: 34 Bayes R2: 0.016 |

##### 4.3.4. Average IBI or RMSSD during rest

*Table S33.* Separately only within systole or diastole trials (first and last 1/3 of the cardiac cycle), the single-trial associations between the continuous measure of P3b amplitude and the degree of change in the current interbeat interval (IBI) compared to the previous one, for the first and second heartbeats after feedback, controlling for the resting average IBI or resting RMSSD. ROPE – region of practical equivalence, BF – Bayes Factor. All values are standardised.

| Parameter | Dependent variable: Change of IBI 1 only in systole | Dependent variable: Change of IBI 2 only in diastole | Dependent variable: Change of IBI 1 only in systole | Dependent variable: Change of IBI 2 only in diastole |
| --- | --- | --- | --- | --- |
| Intercept | Mode: 0.079 95% CI: [0.029,0.158] BF: 0.536 ROPE Bound: 0.018 In ROPE: 0 BF in ROPE: 2.315 RHat: 1.001 | Mode: 0.182 95% CI: [0.114,0.205] BF: 3204 ROPE Bound: 0.019 In ROPE: 0 BF in ROPE: 0.001 RHat: 1.001 | Mode: 0.11 95% CI: [0.027,0.155] BF: 0.598 ROPE Bound: 0.018 In ROPE: 0 BF in ROPE: 2.137 RHat: 1.002 | Mode: 0.144 95% CI: [0.11,0.202] BF: 1491 ROPE Bound: 0.019 In ROPE: 0 BF in ROPE: 0.001 RHat: 1 |
| P3b | Mode: 0.029 95% CI: [0.013,0.069] BF: 16.672 ROPE Bound: 0.003 In ROPE: 0 BF in ROPE: 0.059 RHat: 1 | Mode: 0.054 95% CI: [0.024,0.08] BF: 248 ROPE Bound: 0.003 In ROPE: 0 BF in ROPE: 0.004 RHat: 1.001 | Mode: 0.034 95% CI: [0.011,0.068] BF: 20.466 ROPE Bound: 0.003 In ROPE: 0 BF in ROPE: 0.05 RHat: 1.001 | Mode: 0.053 95% CI: [0.027,0.08] BF: 403 ROPE Bound: 0.003 In ROPE: 0 BF in ROPE: 0.002 RHat: 1 |
| Rest Average IBI | Mode: -0.041 95% CI: [-0.071,0.036] BF: 0.647 ROPE Bound: 0.004 In ROPE: 0.1 BF in ROPE: 1.631 RHat: 1 | Mode: -0.032 95% CI: [-0.069,0.017] BF: 0.826 ROPE Bound: 0.004 In ROPE: 0.085 BF in ROPE: 1.202 RHat: 1 |  |  |
| Rest RMSSD |  |  | Mode: 0.011 95% CI: [-0.06,0.051] BF: 0.557 ROPE Bound: 0.003 In ROPE: 0.084 BF in ROPE: 1.843 RHat: 1 | Mode: -0.024 95% CI: [-0.074,0.017] BF: 1.021 ROPE Bound: 0.003 In ROPE: 0.05 BF in ROPE: 0.998 RHat: 1 |
|  | Family: student Num. obs: 3212 Num. groups: 34 Bayes R2: 0.027 | Family: student Num. obs: 3229 Num. groups: 34 Bayes R2: 0.015 | Family: student Num. obs: 3212 Num. groups: 34 Bayes R2: 0.027 | Family: student Num. obs: 3229 Num. groups: 34 Bayes R2: 0.015 |

##### 4.3.5. Average IBI or RMSSD within a trial, before or after feedback

*Table S34.* Separately only within systole or diastole trials (first and last 1/3 of the cardiac cycle), the single-trial associations between the continuous measure of P3b amplitude and the degree of change in the current interbeat interval (IBI) compared to the previous one, for the first and second heartbeats after feedback, controlling for the average IBI or RMSSD before feedback. ROPE – region of practical equivalence, BF – Bayes Factor. All values are standardised.

| Parameter | Dependent variable: Change of IBI 1 only in systole | Dependent variable: Change of IBI 2 only in diastole | Dependent variable: Change of IBI 1 only in systole | Dependent variable: Change of IBI 2 only in diastole |
| --- | --- | --- | --- | --- |
| Intercept | Mode: 0.109 95% CI: [-0.007,0.175] BF: 0.093 ROPE Bound: 0.018 In ROPE: 0.049 BF in ROPE: 11.805 RHat: 1.002 | Mode: 0.151 95% CI: [0.079,0.213] BF: 19.785 ROPE Bound: 0.019 In ROPE: 0 BF in ROPE: 0.072 RHat: 1.002 | Mode: 0.105 95% CI: [0.03,0.156] BF: 0.617 ROPE Bound: 0.018 In ROPE: 0 BF in ROPE: 2.053 RHat: 1 | Mode: 0.171 95% CI: [0.114,0.204] BF: 5531 ROPE Bound: 0.019 In ROPE: 0 BF in ROPE: 0 RHat: 1 |
| P3b | Mode: 0.048 95% CI: [0.009,0.065] BF: 6.568 ROPE Bound: 0.003 In ROPE: 0 BF in ROPE: 0.15 RHat: 1 | Mode: 0.042 95% CI: [0.021,0.075] BF: 77.898 ROPE Bound: 0.003 In ROPE: 0 BF in ROPE: 0.013 RHat: 1 | Mode: 0.039 95% CI: [0.012,0.07] BF: 11.843 ROPE Bound: 0.003 In ROPE: 0 BF in ROPE: 0.082 RHat: 1 | Mode: 0.058 95% CI: [0.028,0.082] BF: 658 ROPE Bound: 0.003 In ROPE: 0 BF in ROPE: 0.001 RHat: 1 |
| Average IBI before feedback | Mode: -0.236 95% CI: [-0.286,-0.194] BF: 72541278372760 ROPE Bound: 0.001 In ROPE: 0 BF in ROPE: 0 RHat: 1.001 | Mode: -0.2 95% CI: [-0.27,-0.174] BF: 260740339 ROPE Bound: 0.001 In ROPE: 0 BF in ROPE: 0 RHat: 1 |  |  |
| RMSSD before feedback |  |  | Mode: -0.037 95% CI: [-0.089,0.02] BF: 1.157 ROPE Bound: 0 In ROPE: 0.004 BF in ROPE: 0.883 RHat: 1 | Mode: -0.014 95% CI: [-0.049,0.011] BF: 0.474 ROPE Bound: 0 In ROPE: 0.009 BF in ROPE: 2.061 RHat: 1 |
|  | Family: student Num. obs: 3212 Num. groups: 34 Bayes R2: 0.065 | Family: student Num. obs: 3229 Num. groups: 34 Bayes R2: 0.055 | Family: student Num. obs: 3212 Num. groups: 34 Bayes R2: 0.027 | Family: student Num. obs: 3229 Num. groups: 34 Bayes R2: 0.016 |

*Table S35.* Separately only within systole or diastole trials (first and last 1/3 of the cardiac cycle), the single-trial associations between the continuous measure of P3b amplitude and the degree of change in the current interbeat interval (IBI) compared to the previous one, for the first and second heartbeats after feedback, controlling for the average IBI or RMSSD after feedback. ROPE – region of practical equivalence, BF – Bayes Factor. All values are standardised.

| Parameter | Dependent variable: Change of IBI 1 only in systole | Dependent variable: Change of IBI 2 only in diastole | Dependent variable: Change of IBI 1 only in systole | Dependent variable: Change of IBI 2 only in diastole |
| --- | --- | --- | --- | --- |
| Intercept | Mode: 0.105 95% CI: [0.008,0.213] BF: 0.224 ROPE Bound: 0.018 In ROPE: 0.011 BF in ROPE: 4.923 RHat: 1.002 | Mode: 0.152 95% CI: [0.112,0.208] BF: 1872 ROPE Bound: 0.019 In ROPE: 0 BF in ROPE: 0.001 RHat: 1.001 | Mode: 0.113 95% CI: [0.043,0.187] BF: 1.015 ROPE Bound: 0.018 In ROPE: 0 BF in ROPE: 1.208 RHat: 1.002 | Mode: 0.199 95% CI: [0.141,0.255] BF: 3451 ROPE Bound: 0.019 In ROPE: 0 BF in ROPE: 0 RHat: 1.001 |
| P3b | Mode: 0.057 95% CI: [0.018,0.072] BF: 57.241 ROPE Bound: 0.003 In ROPE: 0 BF in ROPE: 0.017 RHat: 1.002 | Mode: 0.059 95% CI: [0.03,0.083] BF: 748 ROPE Bound: 0.003 In ROPE: 0 BF in ROPE: 0.001 RHat: 1.001 | Mode: 0.05 95% CI: [0.014,0.069] BF: 30.277 ROPE Bound: 0.003 In ROPE: 0 BF in ROPE: 0.033 RHat: 1 | Mode: 0.039 95% CI: [0.031,0.084] BF: 1163 ROPE Bound: 0.003 In ROPE: 0 BF in ROPE: 0.001 RHat: 1 |
| Average IBI after feedback | Mode: 0.267 95% CI: [0.221,0.315] BF: 69943229831 ROPE Bound: 0.002 In ROPE: 0 BF in ROPE: 0 RHat: 1.001 | Mode: 0.023 95% CI: [-0.007,0.071] BF: 1.353 ROPE Bound: 0.002 In ROPE: 0.022 BF in ROPE: 0.663 RHat: 1.001 |  |  |
| RMSSD after feedback |  |  | Mode: 0.207 95% CI: [0.179,0.26] BF: 16823747774 ROPE Bound: 0.001 In ROPE: 0 BF in ROPE: 0 RHat: 1 | Mode: 0.195 95% CI: [0.118,0.209] BF: 766835 ROPE Bound: 0.001 In ROPE: 0 BF in ROPE: 0 RHat: 1.002 |
|  | Family: student Num. obs: 3212 Num. groups: 34 Bayes R2: 0.061 | Family: student Num. obs: 3229 Num. groups: 34 Bayes R2: 0.016 | Family: student Num. obs: 3212 Num. groups: 34 Bayes R2: 0.059 | Family: student Num. obs: 3229 Num. groups: 34 Bayes R2: 0.035 |

##### 4.3.6. Number of a trial in a block and approach/avoid decisions

*Table S36.* Separately only within systole or diastole trials (first and last 1/3 of the cardiac cycle), the single-trial associations between the continuous measure of P3b amplitude and the degree of change in the current interbeat interval (IBI) compared to the previous one, for the first and second heartbeats after feedback, controlling for the number of trial in a block (learning effects) or approach/avoid decision trials. ROPE – region of practical equivalence, BF – Bayes Factor. All values are standardised.

| Parameter | Dependent variable: Change of IBI 1 only in systole | Dependent variable: Change of IBI 2 only in diastole | Dependent variable: Change of IBI 1 only in systole | Dependent variable: Change of IBI 2 only in diastole |
| --- | --- | --- | --- | --- |
| Intercept | Mode: 0.117 95% CI: [0.033,0.158] BF: 0.665 ROPE Bound: 0.018 In ROPE: 0 BF in ROPE: 2.099 RHat: 1.001 | Mode: 0.171 95% CI: [0.115,0.206] BF: 4843 ROPE Bound: 0.019 In ROPE: 0 BF in ROPE: 0 RHat: 1 | Mode: 0.101 95% CI: [0.017,0.152] BF: 0.244 ROPE Bound: 0.018 In ROPE: 0.006 BF in ROPE: 4.99 RHat: 1.002 | Mode: 0.124 95% CI: [0.097,0.199] BF: 792 ROPE Bound: 0.019 In ROPE: 0 BF in ROPE: 0.002 RHat: 1.001 |
| P3b | Mode: 0.044 95% CI: [0.012,0.069] BF: 14.817 ROPE Bound: 0.003 In ROPE: 0 BF in ROPE: 0.067 RHat: 1.001 | Mode: 0.066 95% CI: [0.029,0.082] BF: 392 ROPE Bound: 0.003 In ROPE: 0 BF in ROPE: 0.003 RHat: 1 | Mode: 0.052 95% CI: [0.012,0.069] BF: 17.979 ROPE Bound: 0.003 In ROPE: 0 BF in ROPE: 0.058 RHat: 1 | Mode: 0.05 95% CI: [0.027,0.082] BF: 332 ROPE Bound: 0.003 In ROPE: 0 BF in ROPE: 0.003 RHat: 1 |
| Trial in Block | Mode: -0.012 95% CI: [-0.035,0.019] BF: 0.33 ROPE Bound: 0.005 In ROPE: 0.268 BF in ROPE: 3.902 RHat: 1 | Mode: 0.014 95% CI: [-0.003,0.05] BF: 1.499 ROPE Bound: 0.005 In ROPE: 0.059 BF in ROPE: 0.694 RHat: 1.001 |  |  |
| Decision Avoid |  |  | Mode: 0.051 95% CI: [-0.019,0.077] BF: 0.897 ROPE Bound: 0.018 In ROPE: 0.342 BF in ROPE: 1.281 RHat: 1.001 | Mode: 0.021 95% CI: [-0.027,0.068] BF: 0.694 ROPE Bound: 0.019 In ROPE: 0.433 BF in ROPE: 1.672 RHat: 1.001 |
|  | Family: student Num. obs: 3212 Num. groups: 34 Bayes R2: 0.027 | Family: student Num. obs: 3229 Num. groups: 34 Bayes R2: 0.015 | Family: student Num. obs: 3212 Num. groups: 34 Bayes R2: 0.027 | Family: student Num. obs: 3229 Num. groups: 34 Bayes R2: 0.015 |

##### 4.3.7. Reaction times

*Table S37.* Separately only within systole or diastole trials (first and last 1/3 of the cardiac cycle) and only within approach or avoid trials, the single-trial associations between the continuous measure of P3b amplitude and the degree of change in the current interbeat interval (IBI) compared to the previous one, for the first and second heartbeats after feedback, controlling for the reaction time (RT). No RT control in avoid trials. ROPE – region of practical equivalence, BF – Bayes Factor. All values are standardised. Note that fitting the reaction time models only on systole or diastole trials and also only on approach trials affected power: 1789 and 1735 trials were included for R1 in systole and R2 in diastole instead of 3212 and 3229, which was already limited compared to the overall number of trials the main behavioural models are fitted on (9808).

| Parameter | Dependent variable: Change of IBI 1 only in systole only in approach trials | Dependent variable: Change of IBI 2 only in diastole only in approach trials | Dependent variable: Change of IBI 1 only in systole only in avoid trials | Dependent variable: Change of IBI 2 only in diastole only in avoid trials |
| --- | --- | --- | --- | --- |
| Intercept | Mode: 0.062 95% CI: [0.002,0.148] BF: 0.114 ROPE Bound: 0.018 In ROPE: 0.039 BF in ROPE: 10.42 RHat: 1 | Mode: 0.163 95% CI: [0.097,0.215] BF: 150 ROPE Bound: 0.018 In ROPE: 0 BF in ROPE: 0.01 RHat: 1.001 | Mode: 0.153 95% CI: [0.061,0.192] BF: 7.095 ROPE Bound: 0.018 In ROPE: 0 BF in ROPE: 0.193 RHat: 1.001 | Mode: 0.205 95% CI: [0.109,0.233] BF: 102 ROPE Bound: 0.019 In ROPE: 0 BF in ROPE: 0.012 RHat: 1 |
| P3b | Mode: 0.033 95% CI: [-0.005,0.068] BF: 1.491 ROPE Bound: 0.003 In ROPE: 0.031 BF in ROPE: 0.661 RHat: 1.002 | Mode: 0.047 95% CI: [0.013,0.083] BF: 11.939 ROPE Bound: 0.003 In ROPE: 0 BF in ROPE: 0.083 RHat: 1 | Mode: 0.042 95% CI: [0.013,0.089] BF: 10.241 ROPE Bound: 0.003 In ROPE: 0 BF in ROPE: 0.094 RHat: 1 | Mode: 0.078 95% CI: [0.022,0.098] BF: 28.132 ROPE Bound: 0.003 In ROPE: 0 BF in ROPE: 0.032 RHat: 1 |
| RT | Mode: -0.056 95% CI: [-0.103,-0.03] BF: 102 ROPE Bound: 0.004 In ROPE: 0 BF in ROPE: 0.01 RHat: 1 | Mode: -0.072 95% CI: [-0.118,-0.045] BF: 1303 ROPE Bound: 0.004 In ROPE: 0 BF in ROPE: 0.001 RHat: 1 |  |  |
|  | Family: student Num. obs: 1789 Num. groups: 34 Bayes R2: 0.037 | Family: student Num. obs: 1735 Num. groups: 34 Bayes R2: 0.022 | Family: student Num. obs: 1423 Num. groups: 34 Bayes R2: 0.02 | Family: student Num. obs: 1495 Num. groups: 34 Bayes R2: 0.022 |

##### 4.3.8. Prior sensitivity

*Table S38.* Separately only within systole or diastole trials (first and last 1/3 of the cardiac cycle), the single-trial associations between the continuous measure of P3b amplitude and the degree of change in the current interbeat interval (IBI) compared to the previous one, for the first and second heartbeats after feedback, with twice less (normal(0,0.1)) or twice more (normal(0,0.03)) informative priors. ROPE – region of practical equivalence, BF – Bayes Factor. All values are standardised.

| Parameter | Dependent variable: Change of IBI 1 only in systole  Prior: normal(0,0.1) | Dependent variable: Change of IBI 2 only in diastole  Prior: normal(0,0.1) | Dependent variable: Change of IBI 1 only in systole  Prior: normal(0,0.03) | Dependent variable: Change of IBI 2 only in diastole  Prior: normal(0,0.03) |
| --- | --- | --- | --- | --- |
| Intercept | Mode: 0.109 95% CI: [0.029,0.155] BF: 0.655 ROPE Bound: 0.018 In ROPE: 0 BF in ROPE: 2.064 RHat: 1.001 | Mode: 0.154 95% CI: [0.114,0.205] BF: 8092 ROPE Bound: 0.019 In ROPE: 0 BF in ROPE: 0 RHat: 1.002 | Mode: 0.121 95% CI: [0.032,0.157] BF: 0.551 ROPE Bound: 0.018 In ROPE: 0 BF in ROPE: 2.13 RHat: 1.002 | Mode: 0.155 95% CI: [0.112,0.205] BF: 2845 ROPE Bound: 0.019 In ROPE: 0 BF in ROPE: 0.001 RHat: 1 |
| P3b | Mode: 0.041 95% CI: [0.016,0.073] BF: 10.13 ROPE Bound: 0.003 In ROPE: 0 BF in ROPE: 0.098 RHat: 1 | Mode: 0.067 95% CI: [0.03,0.086] BF: 310 ROPE Bound: 0.003 In ROPE: 0 BF in ROPE: 0.003 RHat: 1 | Mode: 0.028 95% CI: [0.01,0.063] BF: 15.128 ROPE Bound: 0.003 In ROPE: 0 BF in ROPE: 0.06 RHat: 1 | Mode: 0.044 95% CI: [0.023,0.075] BF: 120 ROPE Bound: 0.003 In ROPE: 0 BF in ROPE: 0.008 RHat: 1 |
|  | Family: student Num. obs: 3212 Num. groups: 34 Bayes R2: 0.026 | Family: student Num. obs: 3229 Num. groups: 34 Bayes R2: 0.015 | Family: student Num. obs: 3212 Num. groups: 34 Bayes R2: 0.026 | Family: student Num. obs: 3229 Num. groups: 34 Bayes R2: 0.014 |

##### 4.3.9. HRV outlier removal

*Table S39.* Separately only within systole or diastole trials (first and last 1/3 of the cardiac cycle), the single-trial associations between the continuous measure of P3b amplitude and the degree of change in the current interbeat interval (IBI) compared to the previous one, for the first and second heartbeats after feedback, after removing RMSSD outliers. ROPE – region of practical equivalence, BF – Bayes Factor. All values are standardised.

| Parameter | Dependent variable: Change of IBI 1 only in systole | Dependent variable: Change of IBI 2 only in diastole |
| --- | --- | --- |
| Intercept | Mode: 0.102 95% CI: [0.031,0.167] BF: 0.813 ROPE Bound: 0.02 In ROPE: 0 BF in ROPE: 1.701 RHat: 1.001 | Mode: 0.175 95% CI: [0.114,0.217] BF: 496 ROPE Bound: 0.02 In ROPE: 0 BF in ROPE: 0.003 RHat: 1.001 |
| P3b | Mode: 0.046 95% CI: [0.013,0.073] BF: 14.649 ROPE Bound: 0.003 In ROPE: 0 BF in ROPE: 0.068 RHat: 1.002 | Mode: 0.056 95% CI: [0.03,0.089] BF: 297 ROPE Bound: 0.003 In ROPE: 0 BF in ROPE: 0.003 RHat: 1 |
|  | Family: student Num. obs: 3103 Num. groups: 33 Bayes R2: 0.03 | Family: student Num. obs: 3137 Num. groups: 33 Bayes R2: 0.017 |

#### 4.4. Results based on uncorrected IBI values

##### 4.4.1. Controlling for previous IBI

Here we present the same analysis as in Supplementary Material 2.2 but rerun on uncorrected, raw IBI values of each heartbeat surrounding feedback and only controlling for the previous one. To show the specificity of effects to the heartbeats we are focusing on (R -1, 2, 3), we show the results for the three heartbeats before and the four after feedback. You can see that the results are nearly identical in heartbeat associations and directions of associations as we show in the main analysis. R-1 and R3 scale with expectation (no interaction with cardiac phase, Table S40), R2 specifically shows scaling with outcome, PE, and unsigned PE (Tables S41, S42, S43) and an interaction with cardiac phase for PE (Table S42). Interaction of outcome valence with cardiac phase was not detected in these models (BF in ROPE 1.2), probably due to much higher complexity of these models compared to our main analysis (BF 3.2). No other findings emerged, which highlights that the IBI change approach captures well the IBI scaling with prediction, no matter the model specification. Note that in this section we have not checked if positive findings stand after Bayesian FDR correction.

*Table S40.* The single-trial associations between the continuous measure of expectation and the length of the current interbeat interval (IBI), for the three heartbeats before and the four heartbeats after feedback, controlling for the previous IBI. ROPE – region of practical equivalence, BF – Bayes Factor. Cardiac phase is a continuous measure of distance from the last heartbeat before feedback to feedback, scaled by the distance from the last heartbeat before to the first heartbeat after feedback (IBI 1). All values are standardised. Note that RHat is > 1.01 for Intercept (but not other coefficients) in R-2 and R4 models, indicating convergence issues.

| Parameter | Dependent variable: IBI -3 | Dependent variable: IBI -2 | Dependent variable: IBI -1 | Dependent variable: IBI 1 | Dependent variable: IBI 2 | Dependent variable: IBI 3 | Dependent variable: IBI 4 |
| --- | --- | --- | --- | --- | --- | --- | --- |
| Intercept | Mode: -0.031 95% CI: [-0.071,0.056] BF: 0.012 ROPE Bound: 0.008 In ROPE: 0.188 BF in ROPE: 104 RHat: 1.007 | Mode: -0.057 95% CI: [-0.095,0.045] BF: 0.017 ROPE Bound: 0.008 In ROPE: 0.145 BF in ROPE: 66.883 RHat: 1.011 | Mode: -0.01 95% CI: [-0.081,0.081] BF: 0.015 ROPE Bound: 0.008 In ROPE: 0.171 BF in ROPE: 78.683 RHat: 1.005 | Mode: -0.042 95% CI: [-0.066,0.04] BF: 0.011 ROPE Bound: 0.008 In ROPE: 0.225 BF in ROPE: 126 RHat: 1.01 | Mode: 0.013 95% CI: [-0.059,0.069] BF: 0.014 ROPE Bound: 0.009 In ROPE: 0.222 BF in ROPE: 102 RHat: 1.007 | Mode: 0.039 95% CI: [-0.068,0.098] BF: 0.016 ROPE Bound: 0.009 In ROPE: 0.175 BF in ROPE: 74.471 RHat: 1.006 | Mode: 0.038 95% CI: [-0.067,0.083] BF: 0.013 ROPE Bound: 0.009 In ROPE: 0.196 BF in ROPE: 89.488 RHat: 1.021 |
| Expectation | Mode: -0.004 95% CI: [-0.009,0.005] BF: 0.079 ROPE Bound: 0.002 In ROPE: 0.457 BF in ROPE: 20.223 RHat: 1.001 | Mode: -0.001 95% CI: [-0.005,0.009] BF: 0.087 ROPE Bound: 0.002 In ROPE: 0.429 BF in ROPE: 18.601 RHat: 1 | Mode: -0.011 95% CI: [-0.02,-0.006] BF: 28.357 ROPE Bound: 0.002 In ROPE: 0 BF in ROPE: 0.048 RHat: 1 | Mode: -0.014 95% CI: [-0.021,-0.006] BF: 31.6 ROPE Bound: 0.002 In ROPE: 0 BF in ROPE: 0.059 RHat: 1 | Mode: -0.001 95% CI: [-0.01,0.005] BF: 0.098 ROPE Bound: 0.003 In ROPE: 0.417 BF in ROPE: 15.787 RHat: 1 | Mode: 0.011 95% CI: [0.005,0.019] BF: 9.941 ROPE Bound: 0.003 In ROPE: 0 BF in ROPE: 0.166 RHat: 1 | Mode: 0.012 95% CI: [0,0.014] BF: 0.563 ROPE Bound: 0.003 In ROPE: 0.081 BF in ROPE: 2.436 RHat: 1.001 |
| Cardiac phase | Mode: -0.008 95% CI: [-0.011,0.002] BF: 0.156 ROPE Bound: 0.002 In ROPE: 0.244 BF in ROPE: 8.52 RHat: 1 | Mode: 0.003 95% CI: [-0.002,0.011] BF: 0.185 ROPE Bound: 0.002 In ROPE: 0.212 BF in ROPE: 7.036 RHat: 1.001 | Mode: 0.017 95% CI: [0.01,0.023] BF: 1445 ROPE Bound: 0.002 In ROPE: 0 BF in ROPE: 0.001 RHat: 1 | Mode: -0.032 95% CI: [-0.041,-0.027] BF: 9467462 ROPE Bound: 0.002 In ROPE: 0 BF in ROPE: 0 RHat: 1.001 | Mode: 0.052 95% CI: [0.048,0.062] BF: 1719584725064982 ROPE Bound: 0.002 In ROPE: 0 BF in ROPE: 0 RHat: 1.001 | Mode: 0.01 95% CI: [0.002,0.017] BF: 2.265 ROPE Bound: 0.003 In ROPE: 0.005 BF in ROPE: 0.685 RHat: 1.001 | Mode: 0.001 95% CI: [-0.007,0.007] BF: 0.071 ROPE Bound: 0.002 In ROPE: 0.564 BF in ROPE: 27.692 RHat: 1 |
| Expectation : Cardiac phase | Mode: -0.002 95% CI: [-0.006,0.008] BF: 0.069 ROPE Bound: 0.001 In ROPE: 0.164 BF in ROPE: 17.488 RHat: 1.001 | Mode: 0.005 95% CI: [-0.003,0.011] BF: 0.137 ROPE Bound: 0.001 In ROPE: 0.086 BF in ROPE: 7.845 RHat: 1.001 | Mode: -0.007 95% CI: [-0.008,0.005] BF: 0.075 ROPE Bound: 0.001 In ROPE: 0.153 BF in ROPE: 14.68 RHat: 1.001 | Mode: 0.004 95% CI: [-0.009,0.005] BF: 0.082 ROPE Bound: 0.001 In ROPE: 0.139 BF in ROPE: 13.656 RHat: 1 | Mode: -0.012 95% CI: [-0.014,0] BF: 0.475 ROPE Bound: 0.001 In ROPE: 0.02 BF in ROPE: 2.208 RHat: 1 | Mode: -0.002 95% CI: [-0.01,0.004] BF: 0.102 ROPE Bound: 0.001 In ROPE: 0.135 BF in ROPE: 11.244 RHat: 1 | Mode: 0.002 95% CI: [-0.007,0.006] BF: 0.07 ROPE Bound: 0.001 In ROPE: 0.166 BF in ROPE: 16.564 RHat: 1 |
| Previous IBI | Mode: 0.892 95% CI: [0.878,0.907] BF: 3.49055616955334e+170 ROPE Bound: 0 In ROPE: 0 BF in ROPE: 0 RHat: 1.001 | Mode: 0.766 95% CI: [0.748,0.775] BF: 2.32932904361928e+109 ROPE Bound: 0 In ROPE: 0 BF in ROPE: 0 RHat: 1.001 | Mode: 0.762 95% CI: [0.746,0.771] BF: 9.57164516126713e+136 ROPE Bound: 0.001 In ROPE: 0 BF in ROPE: 0 RHat: 1.001 | Mode: 0.787 95% CI: [0.782,0.808] BF: 1.01936528492779e+152 ROPE Bound: 0.001 In ROPE: 0 BF in ROPE: 0 RHat: 1.001 | Mode: 0.786 95% CI: [0.773,0.798] BF: 6.86827551349333e+133 ROPE Bound: 0.001 In ROPE: 0 BF in ROPE: 0 RHat: 1.001 | Mode: 0.753 95% CI: [0.74,0.765] BF: 1.97657705709558e+129 ROPE Bound: 0.001 In ROPE: 0 BF in ROPE: 0 RHat: 1 | Mode: 0.748 95% CI: [0.732,0.757] BF: 6.61653679636478e+126 ROPE Bound: 0.001 In ROPE: 0 BF in ROPE: 0 RHat: 1 |
| Expectation : Previous IBI | Mode: -0.012 95% CI: [-0.019,-0.001] BF: 1.036 ROPE Bound: 0 In ROPE: 0 BF in ROPE: NA RHat: 1 | Mode: 0.006 95% CI: [-0.006,0.01] BF: 0.093 ROPE Bound: 0 In ROPE: 0.018 BF in ROPE: 11.165 RHat: 1 | Mode: 0.004 95% CI: [-0.004,0.011] BF: 0.108 ROPE Bound: 0 In ROPE: 0.03 BF in ROPE: 9.657 RHat: 1.001 | Mode: -0.013 95% CI: [-0.018,-0.001] BF: 1.531 ROPE Bound: 0 In ROPE: 0 BF in ROPE: 0.634 RHat: 1 | Mode: -0.005 95% CI: [-0.011,0.005] BF: 0.102 ROPE Bound: 0 In ROPE: 0.051 BF in ROPE: 10.315 RHat: 1 | Mode: 0.011 95% CI: [-0.003,0.014] BF: 0.169 ROPE Bound: 0 In ROPE: 0.018 BF in ROPE: 5.873 RHat: 1.001 | Mode: 0.003 95% CI: [-0.005,0.011] BF: 0.111 ROPE Bound: 0 In ROPE: 0.031 BF in ROPE: 9.453 RHat: 1 |
| Cardiac phase : Previous IBI | Mode: 0.01 95% CI: [-0.005,0.013] BF: 0.115 ROPE Bound: 0 In ROPE: 0 BF in ROPE: NA RHat: 1 | Mode: -0.004 95% CI: [-0.009,0.006] BF: 0.083 ROPE Bound: 0 In ROPE: 0.018 BF in ROPE: 11.516 RHat: 1.001 | Mode: 0.01 95% CI: [0.001,0.016] BF: 1.036 ROPE Bound: 0 In ROPE: 0 BF in ROPE: 0.936 RHat: 1.001 | Mode: -0.014 95% CI: [-0.02,-0.005] BF: 12.61 ROPE Bound: 0 In ROPE: 0 BF in ROPE: 0.08 RHat: 1.001 | Mode: 0.026 95% CI: [0.019,0.034] BF: 100983 ROPE Bound: 0 In ROPE: 0 BF in ROPE: 0 RHat: 1.001 | Mode: -0.022 95% CI: [-0.031,-0.015] BF: 2317 ROPE Bound: 0 In ROPE: 0 BF in ROPE: 0 RHat: 1 | Mode: -0.012 95% CI: [-0.014,0.001] BF: 0.309 ROPE Bound: 0 In ROPE: 0.011 BF in ROPE: 3.205 RHat: 1 |
| Expectation : Cardiac phase : Previous IBI | Mode: -0.003 95% CI: [-0.009,0.008] BF: 0.083 ROPE Bound: 0 In ROPE: 0 BF in ROPE: NA RHat: 1 | Mode: -0.001 95% CI: [-0.005,0.011] BF: 0.107 ROPE Bound: 0 In ROPE: 0 BF in ROPE: NA RHat: 1 | Mode: -0.003 95% CI: [-0.006,0.009] BF: 0.083 ROPE Bound: 0 In ROPE: 0.024 BF in ROPE: 12.576 RHat: 1.001 | Mode: -0.002 95% CI: [-0.01,0.007] BF: 0.084 ROPE Bound: 0 In ROPE: 0.019 BF in ROPE: 12.051 RHat: 1.001 | Mode: -0.002 95% CI: [-0.009,0.007] BF: 0.085 ROPE Bound: 0 In ROPE: 0.02 BF in ROPE: 11.396 RHat: 1.001 | Mode: 0.003 95% CI: [-0.01,0.006] BF: 0.088 ROPE Bound: 0 In ROPE: 0.021 BF in ROPE: 11.774 RHat: 1.001 | Mode: -0.005 95% CI: [-0.009,0.006] BF: 0.083 ROPE Bound: 0 In ROPE: 0.021 BF in ROPE: 12.552 RHat: 1 |
|  | Family: student Num. obs: 9808 Num. groups: 34 Bayes R2: 0.668 | Family: student Num. obs: 9808 Num. groups: 34 Bayes R2: 0.736 | Family: student Num. obs: 9808 Num. groups: 34 Bayes R2: 0.849 | Family: student Num. obs: 9808 Num. groups: 34 Bayes R2: 0.818 | Family: student Num. obs: 9808 Num. groups: 34 Bayes R2: 0.83 | Family: student Num. obs: 9808 Num. groups: 34 Bayes R2: 0.837 | Family: student Num. obs: 9808 Num. groups: 34 Bayes R2: 0.785 |

*Table S41.* The single-trial associations between the valence of outcome (contrast: lose compared to win) and the length of the current interbeat interval (IBI), for the three heartbeats before and the four heartbeats after feedback, controlling for the previous IBI. ROPE – region of practical equivalence, BF – Bayes Factor. Cardiac phase is a continuous measure of distance from the last heartbeat before feedback to feedback, scaled by the distance from the last heartbeat before to the first heartbeat after feedback (IBI 1). All values are standardised. Note that RHat is > 1.01 for Intercept (but not other coefficients) in R-3, R-1, and R4 models, indicating convergence issues.

| Parameter | Dependent variable: IBI -3 | Dependent variable: IBI -2 | Dependent variable: IBI -1 | Dependent variable: IBI 1 | Dependent variable: IBI 2 | Dependent variable: IBI 3 | Dependent variable: IBI 4 |
| --- | --- | --- | --- | --- | --- | --- | --- |
| Intercept | Mode: -0.012 95% CI: [-0.081,0.05] BF: 0.012 ROPE Bound: 0.008 In ROPE: 0.188 BF in ROPE: 99.602 RHat: 1.022 | Mode: -0.072 95% CI: [-0.092,0.05] BF: 0.015 ROPE Bound: 0.008 In ROPE: 0.154 BF in ROPE: 70.269 RHat: 1.008 | Mode: 0.028 95% CI: [-0.086,0.076] BF: 0.013 ROPE Bound: 0.008 In ROPE: 0.182 BF in ROPE: 90.288 RHat: 1.013 | Mode: -0.022 95% CI: [-0.068,0.043] BF: 0.011 ROPE Bound: 0.008 In ROPE: 0.217 BF in ROPE: 112 RHat: 1.01 | Mode: -0.021 95% CI: [-0.069,0.056] BF: 0.011 ROPE Bound: 0.009 In ROPE: 0.229 BF in ROPE: 113 RHat: 1.004 | Mode: 0.032 95% CI: [-0.067,0.098] BF: 0.014 ROPE Bound: 0.009 In ROPE: 0.18 BF in ROPE: 80.447 RHat: 1.008 | Mode: 0.055 95% CI: [-0.068,0.085] BF: 0.014 ROPE Bound: 0.009 In ROPE: 0.185 BF in ROPE: 86.551 RHat: 1.015 |
| Outcome Lose | Mode: 0.01 95% CI: [-0.007,0.019] BF: 0.184 ROPE Bound: 0.008 In ROPE: 0.632 BF in ROPE: 11.788 RHat: 1.001 | Mode: 0.003 95% CI: [-0.018,0.008] BF: 0.165 ROPE Bound: 0.008 In ROPE: 0.701 BF in ROPE: 14.006 RHat: 1.001 | Mode: 0.012 95% CI: [-0.005,0.022] BF: 0.257 ROPE Bound: 0.008 In ROPE: 0.476 BF in ROPE: 6.232 RHat: 1.002 | Mode: -0.005 95% CI: [-0.016,0.012] BF: 0.147 ROPE Bound: 0.008 In ROPE: 0.773 BF in ROPE: 18.367 RHat: 1 | Mode: 0.032 95% CI: [0.016,0.044] BF: 116 ROPE Bound: 0.009 In ROPE: 0 BF in ROPE: 0.021 RHat: 1.001 | Mode: 0.014 95% CI: [-0.005,0.023] BF: 0.307 ROPE Bound: 0.009 In ROPE: 0.485 BF in ROPE: 5.544 RHat: 1.001 | Mode: -0.014 95% CI: [-0.026,0.001] BF: 0.71 ROPE Bound: 0.009 In ROPE: 0.276 BF in ROPE: 2.546 RHat: 1.001 |
| Cardiac phase | Mode: -0.006 95% CI: [-0.014,0.003] BF: 0.17 ROPE Bound: 0.002 In ROPE: 0.201 BF in ROPE: 6.538 RHat: 1.001 | Mode: 0.006 95% CI: [-0.005,0.013] BF: 0.114 ROPE Bound: 0.002 In ROPE: 0.304 BF in ROPE: 11.391 RHat: 1.001 | Mode: 0.015 95% CI: [0.009,0.027] BF: 76.091 ROPE Bound: 0.002 In ROPE: 0 BF in ROPE: 0.018 RHat: 1.001 | Mode: -0.036 95% CI: [-0.047,-0.028] BF: 2310445 ROPE Bound: 0.002 In ROPE: 0 BF in ROPE: 0 RHat: 1 | Mode: 0.063 95% CI: [0.051,0.07] BF: 903247900048097 ROPE Bound: 0.002 In ROPE: 0 BF in ROPE: 0 RHat: 1 | Mode: 0.009 95% CI: [-0.004,0.015] BF: 0.169 ROPE Bound: 0.003 In ROPE: 0.265 BF in ROPE: 8.009 RHat: 1.001 | Mode: 0.004 95% CI: [-0.005,0.012] BF: 0.121 ROPE Bound: 0.002 In ROPE: 0.322 BF in ROPE: 11.009 RHat: 1 |
| Outcome Lose : Cardiac phase | Mode: 0.013 95% CI: [-0.01,0.015] BF: 0.14 ROPE Bound: 0.002 In ROPE: 0.26 BF in ROPE: 8.904 RHat: 1.001 | Mode: 0.007 95% CI: [-0.01,0.017] BF: 0.151 ROPE Bound: 0.002 In ROPE: 0.238 BF in ROPE: 7.96 RHat: 1 | Mode: -0.012 95% CI: [-0.015,0.012] BF: 0.135 ROPE Bound: 0.002 In ROPE: 0.267 BF in ROPE: 9.493 RHat: 1.001 | Mode: 0.011 95% CI: [-0.007,0.02] BF: 0.223 ROPE Bound: 0.002 In ROPE: 0.186 BF in ROPE: 5.215 RHat: 1 | Mode: -0.013 95% CI: [-0.027,0.001] BF: 0.896 ROPE Bound: 0.002 In ROPE: 0.036 BF in ROPE: 1.215 RHat: 1.001 | Mode: 0.019 95% CI: [-0.003,0.025] BF: 0.447 ROPE Bound: 0.003 In ROPE: 0.1 BF in ROPE: 2.391 RHat: 1.001 | Mode: -0.011 95% CI: [-0.021,0.005] BF: 0.308 ROPE Bound: 0.002 In ROPE: 0.139 BF in ROPE: 3.683 RHat: 1 |
| Previous IBI | Mode: 0.886 95% CI: [0.872,0.904] BF: 2.07247450093026e+147 ROPE Bound: 0 In ROPE: 0 BF in ROPE: 0 RHat: 1 | Mode: 0.757 95% CI: [0.747,0.778] BF: 7.9575551255225e+99 ROPE Bound: 0 In ROPE: 0 BF in ROPE: 0 RHat: 1.001 | Mode: 0.759 95% CI: [0.74,0.768] BF: 8.11025649911952e+143 ROPE Bound: 0.001 In ROPE: 0 BF in ROPE: 0 RHat: 1 | Mode: 0.791 95% CI: [0.774,0.804] BF: 1.59716949991852e+146 ROPE Bound: 0.001 In ROPE: 0 BF in ROPE: 0 RHat: 1.001 | Mode: 0.784 95% CI: [0.772,0.8] BF: 3.4813035340118e+142 ROPE Bound: 0.001 In ROPE: 0 BF in ROPE: 0 RHat: 1 | Mode: 0.755 95% CI: [0.734,0.762] BF: 1.30798200058794e+130 ROPE Bound: 0.001 In ROPE: 0 BF in ROPE: 0 RHat: 1.001 | Mode: 0.737 95% CI: [0.724,0.752] BF: 2.22996929127345e+119 ROPE Bound: 0.001 In ROPE: 0 BF in ROPE: 0 RHat: 1.001 |
| Outcome Lose : Previous IBI | Mode: 0.026 95% CI: [-0.005,0.028] BF: 0.445 ROPE Bound: 0 In ROPE: 0.004 BF in ROPE: 2.216 RHat: 1 | Mode: -0.011 95% CI: [-0.016,0.015] BF: 0.163 ROPE Bound: 0 In ROPE: 0.031 BF in ROPE: 6.304 RHat: 1 | Mode: 0.013 95% CI: [-0.002,0.028] BF: 0.649 ROPE Bound: 0.001 In ROPE: 0.019 BF in ROPE: 1.535 RHat: 1 | Mode: 0.028 95% CI: [0,0.032] BF: 1.159 ROPE Bound: 0.001 In ROPE: 0.003 BF in ROPE: 0.874 RHat: 1 | Mode: 0.01 95% CI: [-0.017,0.013] BF: 0.16 ROPE Bound: 0.001 In ROPE: 0.096 BF in ROPE: 6.776 RHat: 1 | Mode: 0.012 95% CI: [-0.007,0.024] BF: 0.315 ROPE Bound: 0.001 In ROPE: 0.044 BF in ROPE: 3.481 RHat: 1 | Mode: 0.017 95% CI: [0,0.03] BF: 1.397 ROPE Bound: 0.001 In ROPE: 0 BF in ROPE: 0.682 RHat: 1 |
| Cardiac phase : Previous IBI | Mode: 0.014 95% CI: [-0.002,0.02] BF: 0.382 ROPE Bound: 0 In ROPE: 0 BF in ROPE: NA RHat: 1.001 | Mode: -0.004 95% CI: [-0.016,0.005] BF: 0.162 ROPE Bound: 0 In ROPE: 0.009 BF in ROPE: 6.196 RHat: 1 | Mode: 0.008 95% CI: [-0.003,0.016] BF: 0.219 ROPE Bound: 0 In ROPE: 0.015 BF in ROPE: 4.568 RHat: 1.001 | Mode: -0.014 95% CI: [-0.028,-0.007] BF: 22.443 ROPE Bound: 0 In ROPE: 0 BF in ROPE: 0.044 RHat: 1.001 | Mode: 0.014 95% CI: [0.009,0.03] BF: 27.724 ROPE Bound: 0 In ROPE: 0 BF in ROPE: 0.036 RHat: 1 | Mode: -0.028 95% CI: [-0.035,-0.015] BF: 650 ROPE Bound: 0 In ROPE: 0 BF in ROPE: 0.002 RHat: 1.001 | Mode: -0.009 95% CI: [-0.016,0.003] BF: 0.215 ROPE Bound: 0 In ROPE: 0.014 BF in ROPE: 4.845 RHat: 1.001 |
| Outcome Lose : Cardiac phase : Previous IBI | Mode: -0.011 95% CI: [-0.031,0.003] BF: 0.586 ROPE Bound: 0 In ROPE: 0 BF in ROPE: NA RHat: 1 | Mode: 0.004 95% CI: [-0.007,0.024] BF: 0.283 ROPE Bound: 0 In ROPE: 0.006 BF in ROPE: 3.571 RHat: 1 | Mode: 0.013 95% CI: [-0.009,0.02] BF: 0.199 ROPE Bound: 0 In ROPE: 0.016 BF in ROPE: 5.098 RHat: 1.001 | Mode: 0.015 95% CI: [-0.005,0.025] BF: 0.339 ROPE Bound: 0 In ROPE: 0.009 BF in ROPE: 2.914 RHat: 1.002 | Mode: 0.014 95% CI: [0.001,0.032] BF: 1.353 ROPE Bound: 0 In ROPE: 0 BF in ROPE: 0.773 RHat: 1 | Mode: 0.015 95% CI: [-0.01,0.021] BF: 0.203 ROPE Bound: 0 In ROPE: 0.016 BF in ROPE: 4.8 RHat: 1.001 | Mode: 0.001 95% CI: [-0.015,0.014] BF: 0.15 ROPE Bound: 0 In ROPE: 0.023 BF in ROPE: 6.842 RHat: 1.001 |
|  | Family: student Num. obs: 9808 Num. groups: 34 Bayes R2: 0.667 | Family: student Num. obs: 9808 Num. groups: 34 Bayes R2: 0.737 | Family: student Num. obs: 9808 Num. groups: 34 Bayes R2: 0.849 | Family: student Num. obs: 9808 Num. groups: 34 Bayes R2: 0.818 | Family: student Num. obs: 9808 Num. groups: 34 Bayes R2: 0.831 | Family: student Num. obs: 9808 Num. groups: 34 Bayes R2: 0.837 | Family: student Num. obs: 9808 Num. groups: 34 Bayes R2: 0.785 |

*Table S42.* The single-trial associations between the continuous measure of prediction error (PE) and the length of the current interbeat interval (IBI), for the three heartbeats before and the four heartbeats after feedback, controlling for the previous IBI. ROPE – region of practical equivalence, BF – Bayes Factor. Cardiac phase is a continuous measure of distance from the last heartbeat before feedback to feedback, scaled by the distance from the last heartbeat before to the first heartbeat after feedback (IBI 1). All values are standardised. Note that RHat is > 1.01 for Intercept (but not other coefficients) in R-1 and R4 models, indicating convergence issues.

| Parameter | Dependent variable: IBI -3 | Dependent variable: IBI -2 | Dependent variable: IBI -1 | Dependent variable: IBI 1 | Dependent variable: IBI 2 | Dependent variable: IBI 3 | Dependent variable: IBI 4 |
| --- | --- | --- | --- | --- | --- | --- | --- |
| Intercept | Mode: -0.04 95% CI: [-0.07,0.057] BF: 0.012 ROPE Bound: 0.008 In ROPE: 0.191 BF in ROPE: 97.847 RHat: 1.004 | Mode: -0.045 95% CI: [-0.089,0.052] BF: 0.016 ROPE Bound: 0.008 In ROPE: 0.156 BF in ROPE: 74.527 RHat: 1.005 | Mode: 0.012 95% CI: [-0.077,0.08] BF: 0.015 ROPE Bound: 0.008 In ROPE: 0.158 BF in ROPE: 75.443 RHat: 1.011 | Mode: -0.013 95% CI: [-0.069,0.042] BF: 0.011 ROPE Bound: 0.008 In ROPE: 0.23 BF in ROPE: 115 RHat: 1.009 | Mode: -0.001 95% CI: [-0.059,0.073] BF: 0.012 ROPE Bound: 0.009 In ROPE: 0.223 BF in ROPE: 90.635 RHat: 1.005 | Mode: 0.012 95% CI: [-0.066,0.098] BF: 0.015 ROPE Bound: 0.009 In ROPE: 0.182 BF in ROPE: 77.74 RHat: 1.007 | Mode: 0.012 95% CI: [-0.074,0.076] BF: 0.014 ROPE Bound: 0.009 In ROPE: 0.194 BF in ROPE: 94.454 RHat: 1.011 |
| PE | Mode: -0.005 95% CI: [-0.008,0.005] BF: 0.076 ROPE Bound: 0.002 In ROPE: 0.425 BF in ROPE: 20.777 RHat: 1 | Mode: -0.002 95% CI: [-0.005,0.008] BF: 0.075 ROPE Bound: 0.002 In ROPE: 0.42 BF in ROPE: 20.235 RHat: 1 | Mode: 0.002 95% CI: [-0.003,0.01] BF: 0.117 ROPE Bound: 0.002 In ROPE: 0.311 BF in ROPE: 11.801 RHat: 1 | Mode: 0.012 95% CI: [0.001,0.015] BF: 0.992 ROPE Bound: 0.002 In ROPE: 0.028 BF in ROPE: 1.355 RHat: 1 | Mode: -0.01 95% CI: [-0.02,-0.006] BF: 29.439 ROPE Bound: 0.002 In ROPE: 0 BF in ROPE: 0.053 RHat: 1 | Mode: -0.013 95% CI: [-0.018,-0.003] BF: 3.902 ROPE Bound: 0.002 In ROPE: 0 BF in ROPE: 0.337 RHat: 1 | Mode: 0.002 95% CI: [-0.004,0.01] BF: 0.086 ROPE Bound: 0.002 In ROPE: 0.382 BF in ROPE: 15.129 RHat: 1.001 |
| Cardiac phase | Mode: -0.007 95% CI: [-0.01,0.003] BF: 0.154 ROPE Bound: 0.002 In ROPE: 0.261 BF in ROPE: 9.061 RHat: 1 | Mode: 0.003 95% CI: [-0.001,0.011] BF: 0.2 ROPE Bound: 0.002 In ROPE: 0.188 BF in ROPE: 6.319 RHat: 1 | Mode: 0.015 95% CI: [0.01,0.024] BF: 388 ROPE Bound: 0.002 In ROPE: 0 BF in ROPE: 0.004 RHat: 1.001 | Mode: -0.031 95% CI: [-0.042,-0.028] BF: 190553832 ROPE Bound: 0.002 In ROPE: 0 BF in ROPE: 0 RHat: 1.001 | Mode: 0.053 95% CI: [0.047,0.061] BF: 87071922101768 ROPE Bound: 0.002 In ROPE: 0 BF in ROPE: 0 RHat: 1.001 | Mode: 0.012 95% CI: [0.003,0.017] BF: 2.625 ROPE Bound: 0.003 In ROPE: 0.001 BF in ROPE: 0.576 RHat: 1.001 | Mode: 0.004 95% CI: [-0.007,0.007] BF: 0.066 ROPE Bound: 0.002 In ROPE: 0.55 BF in ROPE: 26.785 RHat: 1 |
| PE : Cardiac phase | Mode: -0.002 95% CI: [-0.008,0.005] BF: 0.08 ROPE Bound: 0.001 In ROPE: 0.12 BF in ROPE: 14.26 RHat: 1.001 | Mode: -0.002 95% CI: [-0.01,0.003] BF: 0.119 ROPE Bound: 0.001 In ROPE: 0.081 BF in ROPE: 8.993 RHat: 1.002 | Mode: 0 95% CI: [-0.006,0.008] BF: 0.072 ROPE Bound: 0.001 In ROPE: 0.133 BF in ROPE: 15.737 RHat: 1.001 | Mode: 0.001 95% CI: [-0.009,0.005] BF: 0.093 ROPE Bound: 0.001 In ROPE: 0.106 BF in ROPE: 11.609 RHat: 1 | Mode: 0.01 95% CI: [0.004,0.018] BF: 6.797 ROPE Bound: 0.001 In ROPE: 0 BF in ROPE: 0.154 RHat: 1 | Mode: -0.003 95% CI: [-0.011,0.003] BF: 0.119 ROPE Bound: 0.001 In ROPE: 0.101 BF in ROPE: 9.441 RHat: 1 | Mode: 0.01 95% CI: [-0.003,0.011] BF: 0.141 ROPE Bound: 0.001 In ROPE: 0.073 BF in ROPE: 7.57 RHat: 1.002 |
| Previous IBI | Mode: 0.894 95% CI: [0.879,0.908] BF: 2.04558619911376e+157 ROPE Bound: 0 In ROPE: 0 BF in ROPE: 0 RHat: 1.001 | Mode: 0.76 95% CI: [0.748,0.775] BF: 3.03853221523206e+117 ROPE Bound: 0 In ROPE: 0 BF in ROPE: 0 RHat: 1 | Mode: 0.759 95% CI: [0.747,0.772] BF: 5.33461851557994e+149 ROPE Bound: 0.001 In ROPE: 0 BF in ROPE: 0 RHat: 1.002 | Mode: 0.794 95% CI: [0.783,0.809] BF: 4.9812173394832e+129 ROPE Bound: 0.001 In ROPE: 0 BF in ROPE: 0 RHat: 1 | Mode: 0.794 95% CI: [0.774,0.799] BF: 1.4045634348616e+186 ROPE Bound: 0.001 In ROPE: 0 BF in ROPE: 0 RHat: 1 | Mode: 0.749 95% CI: [0.739,0.764] BF: 1.14185686764085e+172 ROPE Bound: 0.001 In ROPE: 0 BF in ROPE: 0 RHat: 1 | Mode: 0.747 95% CI: [0.732,0.757] BF: 2.58458472292238e+160 ROPE Bound: 0.001 In ROPE: 0 BF in ROPE: 0 RHat: 1.001 |
| PE : Previous IBI | Mode: 0.006 95% CI: [-0.004,0.014] BF: 0.176 ROPE Bound: 0 In ROPE: 0 BF in ROPE: NA RHat: 1 | Mode: 0.001 95% CI: [-0.004,0.011] BF: 0.111 ROPE Bound: 0 In ROPE: 0.016 BF in ROPE: 9.549 RHat: 1 | Mode: -0.006 95% CI: [-0.013,0.003] BF: 0.177 ROPE Bound: 0 In ROPE: 0.019 BF in ROPE: 5.854 RHat: 1 | Mode: 0.002 95% CI: [-0.007,0.009] BF: 0.08 ROPE Bound: 0 In ROPE: 0.041 BF in ROPE: 13.269 RHat: 1 | Mode: 0.01 95% CI: [-0.002,0.014] BF: 0.272 ROPE Bound: 0 In ROPE: 0.011 BF in ROPE: 3.775 RHat: 1.001 | Mode: -0.001 95% CI: [-0.011,0.005] BF: 0.124 ROPE Bound: 0 In ROPE: 0.034 BF in ROPE: 8.883 RHat: 1 | Mode: -0.006 95% CI: [-0.013,0.002] BF: 0.219 ROPE Bound: 0 In ROPE: 0.016 BF in ROPE: 4.61 RHat: 1 |
| Cardiac phase : Previous IBI | Mode: 0.006 95% CI: [-0.006,0.011] BF: 0.103 ROPE Bound: 0 In ROPE: 0 BF in ROPE: NA RHat: 1 | Mode: -0.005 95% CI: [-0.009,0.007] BF: 0.082 ROPE Bound: 0 In ROPE: 0.021 BF in ROPE: 12.455 RHat: 1.001 | Mode: 0.015 95% CI: [0.001,0.016] BF: 1.113 ROPE Bound: 0 In ROPE: 0 BF in ROPE: 0.846 RHat: 1 | Mode: -0.012 95% CI: [-0.021,-0.006] BF: 14.134 ROPE Bound: 0 In ROPE: 0 BF in ROPE: 0.068 RHat: 1.001 | Mode: 0.026 95% CI: [0.019,0.034] BF: 108467 ROPE Bound: 0 In ROPE: 0 BF in ROPE: 0 RHat: 1 | Mode: -0.023 95% CI: [-0.03,-0.015] BF: 109113 ROPE Bound: 0 In ROPE: 0 BF in ROPE: 0 RHat: 1 | Mode: -0.011 95% CI: [-0.013,0.001] BF: 0.326 ROPE Bound: 0 In ROPE: 0.011 BF in ROPE: 2.898 RHat: 1.001 |
| PE : Cardiac phase : Previous IBI | Mode: 0.014 95% CI: [-0.002,0.015] BF: 0.288 ROPE Bound: 0 In ROPE: 0 BF in ROPE: NA RHat: 1 | Mode: -0.011 95% CI: [-0.013,0.002] BF: 0.194 ROPE Bound: 0 In ROPE: 0 BF in ROPE: NA RHat: 1.001 | Mode: -0.004 95% CI: [-0.011,0.004] BF: 0.114 ROPE Bound: 0 In ROPE: 0.013 BF in ROPE: 8.763 RHat: 1.001 | Mode: -0.001 95% CI: [-0.012,0.003] BF: 0.14 ROPE Bound: 0 In ROPE: 0.012 BF in ROPE: 7.206 RHat: 1 | Mode: -0.01 95% CI: [-0.015,0.001] BF: 0.317 ROPE Bound: 0 In ROPE: 0.004 BF in ROPE: 3.11 RHat: 1 | Mode: -0.001 95% CI: [-0.01,0.005] BF: 0.094 ROPE Bound: 0 In ROPE: 0.017 BF in ROPE: 10.957 RHat: 1 | Mode: 0.008 95% CI: [-0.006,0.009] BF: 0.084 ROPE Bound: 0 In ROPE: 0.018 BF in ROPE: 13.013 RHat: 1 |
|  | Family: student Num. obs: 9808 Num. groups: 34 Bayes R2: 0.669 | Family: student Num. obs: 9808 Num. groups: 34 Bayes R2: 0.737 | Family: student Num. obs: 9808 Num. groups: 34 Bayes R2: 0.849 | Family: student Num. obs: 9808 Num. groups: 34 Bayes R2: 0.818 | Family: student Num. obs: 9808 Num. groups: 34 Bayes R2: 0.831 | Family: student Num. obs: 9808 Num. groups: 34 Bayes R2: 0.837 | Family: student Num. obs: 9808 Num. groups: 34 Bayes R2: 0.785 |

*Table S43.* The single-trial associations between the continuous measure of unsigned prediction error (unsigned PE) and the length of the current interbeat interval (IBI), for the three heartbeats before and the four heartbeats after feedback, controlling for the previous IBI. ROPE – region of practical equivalence, BF – Bayes Factor. Cardiac phase is a continuous measure of distance from the last heartbeat before feedback to feedback, scaled by the distance from the last heartbeat before to the first heartbeat after feedback (IBI 1). All values are standardised. Note that RHat is > 1.01 for Intercept (but not other coefficients) in R-3, R-2, R1, R2, and R3 models, indicating convergence issues.

| Parameter | Dependent variable: IBI -3 | Dependent variable: IBI -2 | Dependent variable: IBI -1 | Dependent variable: IBI 1 | Dependent variable: IBI 2 | Dependent variable: IBI 3 | Dependent variable: IBI 4 |
| --- | --- | --- | --- | --- | --- | --- | --- |
| Intercept | Mode: -0.016 95% CI: [-0.075,0.063] BF: 0.014 ROPE Bound: 0.008 In ROPE: 0.18 BF in ROPE: 95.21 RHat: 1.011 | Mode: -0.013 95% CI: [-0.086,0.043] BF: 0.015 ROPE Bound: 0.008 In ROPE: 0.145 BF in ROPE: 76.462 RHat: 1.02 | Mode: 0.029 95% CI: [-0.076,0.087] BF: 0.017 ROPE Bound: 0.008 In ROPE: 0.148 BF in ROPE: 68.593 RHat: 1.005 | Mode: -0.058 95% CI: [-0.068,0.042] BF: 0.011 ROPE Bound: 0.008 In ROPE: 0.226 BF in ROPE: 119 RHat: 1.012 | Mode: 0.021 95% CI: [-0.056,0.068] BF: 0.012 ROPE Bound: 0.009 In ROPE: 0.209 BF in ROPE: 106 RHat: 1.013 | Mode: 0.012 95% CI: [-0.063,0.093] BF: 0.015 ROPE Bound: 0.009 In ROPE: 0.184 BF in ROPE: 80.723 RHat: 1.012 | Mode: 0.031 95% CI: [-0.071,0.082] BF: 0.014 ROPE Bound: 0.009 In ROPE: 0.193 BF in ROPE: 85.599 RHat: 1.003 |
| Unsigned PE | Mode: -0.007 95% CI: [-0.01,0.003] BF: 0.115 ROPE Bound: 0.003 In ROPE: 0.371 BF in ROPE: 12.383 RHat: 1 | Mode: 0.002 95% CI: [-0.007,0.006] BF: 0.072 ROPE Bound: 0.003 In ROPE: 0.583 BF in ROPE: 27.931 RHat: 1.001 | Mode: 0.008 95% CI: [-0.002,0.012] BF: 0.195 ROPE Bound: 0.003 In ROPE: 0.246 BF in ROPE: 7.105 RHat: 1 | Mode: 0.002 95% CI: [-0.005,0.009] BF: 0.083 ROPE Bound: 0.003 In ROPE: 0.531 BF in ROPE: 20.998 RHat: 1.001 | Mode: 0.018 95% CI: [0.008,0.023] BF: 152 ROPE Bound: 0.003 In ROPE: 0 BF in ROPE: 0.013 RHat: 1.001 | Mode: 0.011 95% CI: [0.003,0.018] BF: 3.313 ROPE Bound: 0.003 In ROPE: 0.003 BF in ROPE: 0.528 RHat: 1 | Mode: -0.004 95% CI: [-0.01,0.003] BF: 0.107 ROPE Bound: 0.003 In ROPE: 0.458 BF in ROPE: 15.589 RHat: 1 |
| Cardiac phase | Mode: -0.003 95% CI: [-0.011,0.002] BF: 0.166 ROPE Bound: 0.002 In ROPE: 0.239 BF in ROPE: 7.901 RHat: 1 | Mode: 0.008 95% CI: [-0.001,0.011] BF: 0.204 ROPE Bound: 0.002 In ROPE: 0.188 BF in ROPE: 6.334 RHat: 1.001 | Mode: 0.017 95% CI: [0.01,0.024] BF: 785 ROPE Bound: 0.002 In ROPE: 0 BF in ROPE: 0.002 RHat: 1 | Mode: -0.034 95% CI: [-0.041,-0.027] BF: 370725090 ROPE Bound: 0.002 In ROPE: 0 BF in ROPE: 0 RHat: 1.001 | Mode: 0.055 95% CI: [0.047,0.062] BF: 784894501698 ROPE Bound: 0.002 In ROPE: 0 BF in ROPE: 0 RHat: 1 | Mode: 0.011 95% CI: [0.002,0.017] BF: 2.546 ROPE Bound: 0.003 In ROPE: 0.001 BF in ROPE: 0.588 RHat: 1.001 | Mode: 0.001 95% CI: [-0.006,0.007] BF: 0.07 ROPE Bound: 0.002 In ROPE: 0.554 BF in ROPE: 27.667 RHat: 1 |
| Unsigned PE : Cardiac phase | Mode: -0.005 95% CI: [-0.011,0.002] BF: 0.152 ROPE Bound: 0.001 In ROPE: 0.084 BF in ROPE: 7.197 RHat: 1.001 | Mode: -0.001 95% CI: [-0.009,0.005] BF: 0.076 ROPE Bound: 0.001 In ROPE: 0.177 BF in ROPE: 15.033 RHat: 1 | Mode: -0.002 95% CI: [-0.007,0.007] BF: 0.068 ROPE Bound: 0.001 In ROPE: 0.198 BF in ROPE: 18.224 RHat: 1 | Mode: 0.002 95% CI: [-0.005,0.009] BF: 0.087 ROPE Bound: 0.001 In ROPE: 0.154 BF in ROPE: 12.923 RHat: 1 | Mode: -0.011 95% CI: [-0.013,0.001] BF: 0.447 ROPE Bound: 0.001 In ROPE: 0.028 BF in ROPE: 2.259 RHat: 1 | Mode: 0.012 95% CI: [0,0.015] BF: 0.567 ROPE Bound: 0.001 In ROPE: 0.015 BF in ROPE: 1.799 RHat: 1 | Mode: -0.002 95% CI: [-0.01,0.004] BF: 0.099 ROPE Bound: 0.001 In ROPE: 0.148 BF in ROPE: 10.947 RHat: 1 |
| Previous IBI | Mode: 0.889 95% CI: [0.879,0.907] BF: 8.88957296009768e+188 ROPE Bound: 0 In ROPE: 0 BF in ROPE: 0 RHat: 1.001 | Mode: 0.764 95% CI: [0.748,0.775] BF: 6.53068206264776e+167 ROPE Bound: 0 In ROPE: 0 BF in ROPE: 0 RHat: 1.001 | Mode: 0.754 95% CI: [0.747,0.771] BF: 7.08159596417893e+133 ROPE Bound: 0.001 In ROPE: 0 BF in ROPE: 0 RHat: 1.002 | Mode: 0.797 95% CI: [0.784,0.809] BF: 5.10192651405375e+150 ROPE Bound: 0.001 In ROPE: 0 BF in ROPE: 0 RHat: 1 | Mode: 0.788 95% CI: [0.772,0.798] BF: 3.19571558694692e+158 ROPE Bound: 0.001 In ROPE: 0 BF in ROPE: 0 RHat: 1.001 | Mode: 0.751 95% CI: [0.739,0.764] BF: 6.03225328973009e+143 ROPE Bound: 0.001 In ROPE: 0 BF in ROPE: 0 RHat: 1.001 | Mode: 0.74 95% CI: [0.732,0.756] BF: 2.57125053284088e+135 ROPE Bound: 0.001 In ROPE: 0 BF in ROPE: 0 RHat: 1.001 |
| Unsigned PE : Previous IBI | Mode: -0.01 95% CI: [-0.012,0.005] BF: 0.134 ROPE Bound: 0 In ROPE: 0 BF in ROPE: NA RHat: 1 | Mode: -0.001 95% CI: [-0.01,0.005] BF: 0.099 ROPE Bound: 0 In ROPE: 0.018 BF in ROPE: 10.316 RHat: 1 | Mode: -0.002 95% CI: [-0.007,0.008] BF: 0.078 ROPE Bound: 0 In ROPE: 0.039 BF in ROPE: 12.841 RHat: 1.001 | Mode: 0.001 95% CI: [-0.003,0.013] BF: 0.145 ROPE Bound: 0 In ROPE: 0.032 BF in ROPE: 7.003 RHat: 1 | Mode: 0.002 95% CI: [-0.004,0.012] BF: 0.147 ROPE Bound: 0 In ROPE: 0.034 BF in ROPE: 7.107 RHat: 1.002 | Mode: 0.003 95% CI: [-0.006,0.01] BF: 0.09 ROPE Bound: 0 In ROPE: 0.062 BF in ROPE: 11.851 RHat: 1.001 | Mode: 0.001 95% CI: [-0.007,0.008] BF: 0.075 ROPE Bound: 0 In ROPE: 0.066 BF in ROPE: 14.244 RHat: 1.002 |
| Cardiac phase : Previous IBI | Mode: -0.001 95% CI: [-0.005,0.012] BF: 0.114 ROPE Bound: 0 In ROPE: 0 BF in ROPE: NA RHat: 1 | Mode: 0.004 95% CI: [-0.009,0.006] BF: 0.083 ROPE Bound: 0 In ROPE: 0.02 BF in ROPE: 12.21 RHat: 1.001 | Mode: 0.01 95% CI: [0.001,0.016] BF: 1.147 ROPE Bound: 0 In ROPE: 0 BF in ROPE: 0.874 RHat: 1.001 | Mode: -0.017 95% CI: [-0.021,-0.006] BF: 18.277 ROPE Bound: 0 In ROPE: 0 BF in ROPE: 0.055 RHat: 1 | Mode: 0.027 95% CI: [0.019,0.034] BF: 17899 ROPE Bound: 0 In ROPE: 0 BF in ROPE: 0 RHat: 1 | Mode: -0.024 95% CI: [-0.03,-0.015] BF: 6762 ROPE Bound: 0 In ROPE: 0 BF in ROPE: 0 RHat: 1.001 | Mode: -0.014 95% CI: [-0.014,0.001] BF: 0.284 ROPE Bound: 0 In ROPE: 0.012 BF in ROPE: 3.56 RHat: 1 |
| Unsigned PE : Cardiac phase : Previous IBI | Mode: -0.005 95% CI: [-0.011,0.006] BF: 0.099 ROPE Bound: 0 In ROPE: 0 BF in ROPE: NA RHat: 1 | Mode: 0.002 95% CI: [-0.007,0.009] BF: 0.082 ROPE Bound: 0 In ROPE: 0 BF in ROPE: NA RHat: 1.001 | Mode: 0.002 95% CI: [-0.004,0.011] BF: 0.106 ROPE Bound: 0 In ROPE: 0.015 BF in ROPE: 8.919 RHat: 1.001 | Mode: 0.002 95% CI: [-0.005,0.01] BF: 0.098 ROPE Bound: 0 In ROPE: 0.016 BF in ROPE: 10.357 RHat: 1 | Mode: -0.002 95% CI: [-0.007,0.009] BF: 0.083 ROPE Bound: 0 In ROPE: 0.021 BF in ROPE: 12.032 RHat: 1 | Mode: 0.004 95% CI: [-0.007,0.009] BF: 0.088 ROPE Bound: 0 In ROPE: 0.02 BF in ROPE: 11.749 RHat: 1 | Mode: 0 95% CI: [-0.007,0.008] BF: 0.081 ROPE Bound: 0 In ROPE: 0.025 BF in ROPE: 12.083 RHat: 1.001 |
|  | Family: student Num. obs: 9808 Num. groups: 34 Bayes R2: 0.668 | Family: student Num. obs: 9808 Num. groups: 34 Bayes R2: 0.737 | Family: student Num. obs: 9808 Num. groups: 34 Bayes R2: 0.849 | Family: student Num. obs: 9808 Num. groups: 34 Bayes R2: 0.818 | Family: student Num. obs: 9808 Num. groups: 34 Bayes R2: 0.831 | Family: student Num. obs: 9808 Num. groups: 34 Bayes R2: 0.837 | Family: student Num. obs: 9808 Num. groups: 34 Bayes R2: 0.785 |

##### 4.4.2. Without controlling for previous IBI

Without controlling for previous IBI, it is difficult to distinguish immediate IBI change from the correlations with neighbouring IBI which makes the observed results less certain and more distributed across several heartbeats, complicating interpretation and reducing the strength of inference. Associations with expectation were detected for R-1 (as before), R1, R2, but not R3 (Table S44). From our beat-to-beat interpretation, it is because R-1 reflects expectation, R1 and R2 share some continuing correlation but not many new effects, and R3 change shows the return to initial dynamics, in proportion to how it was modulated prior to feedback at R-1. Effects of outcome valence were detected at R2 (as before) and R3 (probably due to correlation with R2), but no interaction with cardiac phase was observed, which is mechanistically unlikely (Table S45). For PE, we detected only moderate evidence (BF 5) at R3 (Table S46), which is very different from our main findings where we observe strong evidence for PE scaling at R2, as well as interactions with cardiac phase. For unsigned PE, we observed, similarly to the main findings, modulation at R2, but also, unlike in the other findings, at R3, also probably due to correlation with R2 (Table S47). These findings do not provide additional mechanistic insight because as soon as we control for previous IBI length, as we did in the previous section, we observe the same pattern of findings as in our main models. The results in the previous section also show how strongly the previous IBI predicts the current one (Tables S40-S43). Therefore, we show that the IBI change approach (“beat-to-beat”) may be more precise and interpretable (we always only consider immediate parasympathetic influences on the pacemaker) and results in stronger inference. Note that in this section we have not checked if positive findings stand after Bayesian FDR correction.

*Table S44.* The single-trial associations between the continuous measure of expectation and the length of the current interbeat interval (IBI), for the three heartbeats before and the four heartbeats after feedback. ROPE – region of practical equivalence, BF – Bayes Factor. Cardiac phase is a continuous measure of distance from the last heartbeat before feedback to feedback, scaled by the distance from the last heartbeat before to the first heartbeat after feedback (IBI 1). All values are standardised. Note that RHat is > 1.01 for Intercept (but not other coefficients) in R-1, R2 and R4 models, indicating convergence issues.

| Parameter | Dependent variable: IBI -3 | Dependent variable: IBI -2 | Dependent variable: IBI -1 | Dependent variable: IBI 1 | Dependent variable: IBI 2 | Dependent variable: IBI 3 | Dependent variable: IBI 4 |
| --- | --- | --- | --- | --- | --- | --- | --- |
| Intercept | Mode: -0.171 95% CI: [-0.266,0.272] BF: 0.054 ROPE Bound: 0.008 In ROPE: 0.042 BF in ROPE: 18.658 RHat: 1.007 | Mode: 0.313 95% CI: [-0.276,0.365] BF: 0.058 ROPE Bound: 0.008 In ROPE: 0.042 BF in ROPE: 16.427 RHat: 1.007 | Mode: 0.133 95% CI: [-0.25,0.396] BF: 0.067 ROPE Bound: 0.008 In ROPE: 0.038 BF in ROPE: 15.528 RHat: 1.011 | Mode: -0.156 95% CI: [-0.229,0.372] BF: 0.06 ROPE Bound: 0.008 In ROPE: 0.042 BF in ROPE: 16.772 RHat: 1.008 | Mode: 0.116 95% CI: [-0.271,0.368] BF: 0.061 ROPE Bound: 0.009 In ROPE: 0.043 BF in ROPE: 16.887 RHat: 1.026 | Mode: 0.238 95% CI: [-0.242,0.329] BF: 0.06 ROPE Bound: 0.009 In ROPE: 0.05 BF in ROPE: 17.036 RHat: 1.004 | Mode: 0.135 95% CI: [-0.223,0.332] BF: 0.051 ROPE Bound: 0.009 In ROPE: 0.051 BF in ROPE: 20.304 RHat: 1.015 |
| Expectation | Mode: 0.004 95% CI: [-0.007,0.014] BF: 0.135 ROPE Bound: 0.002 In ROPE: 0.29 BF in ROPE: 10.469 RHat: 1.002 | Mode: -0.003 95% CI: [-0.01,0.011] BF: 0.116 ROPE Bound: 0.002 In ROPE: 0.334 BF in ROPE: 12.131 RHat: 1.001 | Mode: -0.012 95% CI: [-0.026,-0.005] BF: 6.589 ROPE Bound: 0.002 In ROPE: 0 BF in ROPE: 0.184 RHat: 1.003 | Mode: -0.029 95% CI: [-0.034,-0.012] BF: 95.512 ROPE Bound: 0.002 In ROPE: 0 BF in ROPE: 0.011 RHat: 1.001 | Mode: -0.018 95% CI: [-0.029,-0.006] BF: 5.358 ROPE Bound: 0.003 In ROPE: 0 BF in ROPE: 0.212 RHat: 1.001 | Mode: -0.004 95% CI: [-0.012,0.011] BF: 0.117 ROPE Bound: 0.003 In ROPE: 0.359 BF in ROPE: 11.219 RHat: 1.002 | Mode: 0.005 95% CI: [-0.004,0.018] BF: 0.234 ROPE Bound: 0.003 In ROPE: 0.174 BF in ROPE: 4.698 RHat: 1.001 |
| Cardiac phase | Mode: -0.013 95% CI: [-0.027,-0.006] BF: 10.183 ROPE Bound: 0.002 In ROPE: 0 BF in ROPE: 0.116 RHat: 1.002 | Mode: -0.011 95% CI: [-0.016,0.004] BF: 0.191 ROPE Bound: 0.002 In ROPE: 0.208 BF in ROPE: 6.608 RHat: 1.002 | Mode: 0.015 95% CI: [0.002,0.023] BF: 1.625 ROPE Bound: 0.002 In ROPE: 0.004 BF in ROPE: 0.72 RHat: 1.003 | Mode: -0.026 95% CI: [-0.033,-0.012] BF: 331 ROPE Bound: 0.002 In ROPE: 0 BF in ROPE: 0.003 RHat: 1.002 | Mode: 0.034 95% CI: [0.025,0.048] BF: 1469364 ROPE Bound: 0.002 In ROPE: 0 BF in ROPE: 0 RHat: 1.001 | Mode: 0.033 95% CI: [0.027,0.047] BF: 290442 ROPE Bound: 0.003 In ROPE: 0 BF in ROPE: 0 RHat: 1.002 | Mode: 0.029 95% CI: [0.016,0.037] BF: 11934 ROPE Bound: 0.002 In ROPE: 0 BF in ROPE: 0 RHat: 1.002 |
| Expectation : Cardiac phase | Mode: 0.002 95% CI: [-0.013,0.008] BF: 0.119 ROPE Bound: 0.001 In ROPE: 0.102 BF in ROPE: 8.907 RHat: 1.002 | Mode: -0.004 95% CI: [-0.011,0.01] BF: 0.104 ROPE Bound: 0.001 In ROPE: 0.11 BF in ROPE: 10.673 RHat: 1.001 | Mode: 0.002 95% CI: [-0.01,0.01] BF: 0.112 ROPE Bound: 0.001 In ROPE: 0.115 BF in ROPE: 10.71 RHat: 1.003 | Mode: 0.001 95% CI: [-0.01,0.011] BF: 0.104 ROPE Bound: 0.001 In ROPE: 0.11 BF in ROPE: 10.703 RHat: 1.001 | Mode: -0.014 95% CI: [-0.016,0.006] BF: 0.174 ROPE Bound: 0.001 In ROPE: 0.079 BF in ROPE: 6.437 RHat: 1.001 | Mode: -0.003 95% CI: [-0.014,0.008] BF: 0.117 ROPE Bound: 0.001 In ROPE: 0.113 BF in ROPE: 9.724 RHat: 1.003 | Mode: -0.01 95% CI: [-0.013,0.008] BF: 0.119 ROPE Bound: 0.001 In ROPE: 0.098 BF in ROPE: 9.147 RHat: 1.001 |
|  | Family: student Num. obs: 9808 Num. groups: 34 Bayes R2: 0.535 | Family: student Num. obs: 9808 Num. groups: 34 Bayes R2: 0.615 | Family: student Num. obs: 9808 Num. groups: 34 Bayes R2: 0.636 | Family: student Num. obs: 9808 Num. groups: 34 Bayes R2: 0.62 | Family: student Num. obs: 9808 Num. groups: 34 Bayes R2: 0.61 | Family: student Num. obs: 9808 Num. groups: 34 Bayes R2: 0.616 | Family: student Num. obs: 9808 Num. groups: 34 Bayes R2: 0.59 |

*Table S45.* The single-trial associations between the valence of outcome (contrast: lose compared to win) and the length of the current interbeat interval (IBI), for the three heartbeats before and the four heartbeats after feedback. ROPE – region of practical equivalence, BF – Bayes Factor. Cardiac phase is a continuous measure of distance from the last heartbeat before feedback to feedback, scaled by the distance from the last heartbeat before to the first heartbeat after feedback (IBI 1). All values are standardised. Note that RHat is > 1.01 for Intercept (but not other coefficients) in R-3, R-1, and R1 models, indicating convergence issues.

| Parameter | Dependent variable: IBI -3 | Dependent variable: IBI -2 | Dependent variable: IBI -1 | Dependent variable: IBI 1 | Dependent variable: IBI 2 | Dependent variable: IBI 3 | Dependent variable: IBI 4 |
| --- | --- | --- | --- | --- | --- | --- | --- |
| Intercept | Mode: 0.182 95% CI: [-0.224,0.336] BF: 0.053 ROPE Bound: 0.008 In ROPE: 0.047 BF in ROPE: 20.267 RHat: 1.016 | Mode: 0.143 95% CI: [-0.256,0.348] BF: 0.059 ROPE Bound: 0.008 In ROPE: 0.037 BF in ROPE: 17.711 RHat: 1.008 | Mode: 0.187 95% CI: [-0.284,0.353] BF: 0.062 ROPE Bound: 0.008 In ROPE: 0.038 BF in ROPE: 15.416 RHat: 1.029 | Mode: 0.024 95% CI: [-0.275,0.384] BF: 0.066 ROPE Bound: 0.008 In ROPE: 0.037 BF in ROPE: 15.392 RHat: 1.025 | Mode: -0.111 95% CI: [-0.237,0.33] BF: 0.053 ROPE Bound: 0.009 In ROPE: 0.051 BF in ROPE: 20.364 RHat: 1.009 | Mode: 0.114 95% CI: [-0.258,0.327] BF: 0.046 ROPE Bound: 0.009 In ROPE: 0.063 BF in ROPE: 22.177 RHat: 1.008 | Mode: 0.16 95% CI: [-0.256,0.351] BF: 0.058 ROPE Bound: 0.009 In ROPE: 0.051 BF in ROPE: 17.857 RHat: 1.01 |
| Outcome Lose | Mode: 0.013 95% CI: [-0.014,0.027] BF: 0.252 ROPE Bound: 0.008 In ROPE: 0.478 BF in ROPE: 5.817 RHat: 1.002 | Mode: -0.01 95% CI: [-0.025,0.016] BF: 0.234 ROPE Bound: 0.008 In ROPE: 0.526 BF in ROPE: 7.063 RHat: 1.001 | Mode: 0.022 95% CI: [-0.005,0.037] BF: 0.596 ROPE Bound: 0.008 In ROPE: 0.238 BF in ROPE: 2.084 RHat: 1.003 | Mode: -0.013 95% CI: [-0.017,0.025] BF: 0.255 ROPE Bound: 0.008 In ROPE: 0.518 BF in ROPE: 6.475 RHat: 1.004 | Mode: 0.041 95% CI: [0.011,0.057] BF: 11.047 ROPE Bound: 0.009 In ROPE: 0 BF in ROPE: 0.123 RHat: 1.002 | Mode: 0.032 95% CI: [0.011,0.053] BF: 13.391 ROPE Bound: 0.009 In ROPE: 0 BF in ROPE: 0.106 RHat: 1.002 | Mode: 0.019 95% CI: [-0.015,0.028] BF: 0.266 ROPE Bound: 0.009 In ROPE: 0.53 BF in ROPE: 6.375 RHat: 1 |
| Cardiac phase | Mode: -0.021 95% CI: [-0.033,-0.006] BF: 6.612 ROPE Bound: 0.002 In ROPE: 0 BF in ROPE: 0.163 RHat: 1.002 | Mode: -0.013 95% CI: [-0.022,0.005] BF: 0.29 ROPE Bound: 0.002 In ROPE: 0.132 BF in ROPE: 3.822 RHat: 1.003 | Mode: 0.011 95% CI: [-0.004,0.023] BF: 0.4 ROPE Bound: 0.002 In ROPE: 0.109 BF in ROPE: 2.79 RHat: 1.002 | Mode: -0.022 95% CI: [-0.042,-0.014] BF: 109 ROPE Bound: 0.002 In ROPE: 0 BF in ROPE: 0.01 RHat: 1.001 | Mode: 0.04 95% CI: [0.026,0.055] BF: 2303 ROPE Bound: 0.002 In ROPE: 0 BF in ROPE: 0 RHat: 1.001 | Mode: 0.035 95% CI: [0.022,0.05] BF: 5032 ROPE Bound: 0.003 In ROPE: 0 BF in ROPE: 0 RHat: 1.003 | Mode: 0.023 95% CI: [0.014,0.04] BF: 215 ROPE Bound: 0.002 In ROPE: 0 BF in ROPE: 0.005 RHat: 1.001 |
| Outcome Lose : Cardiac phase | Mode: 0.014 95% CI: [-0.013,0.028] BF: 0.247 ROPE Bound: 0.002 In ROPE: 0.155 BF in ROPE: 4.798 RHat: 1.001 | Mode: 0.014 95% CI: [-0.014,0.027] BF: 0.254 ROPE Bound: 0.002 In ROPE: 0.133 BF in ROPE: 4.406 RHat: 1.003 | Mode: 0.016 95% CI: [-0.016,0.024] BF: 0.235 ROPE Bound: 0.002 In ROPE: 0.17 BF in ROPE: 4.866 RHat: 1.003 | Mode: 0.012 95% CI: [-0.011,0.031] BF: 0.316 ROPE Bound: 0.002 In ROPE: 0.122 BF in ROPE: 3.395 RHat: 1 | Mode: -0.018 95% CI: [-0.028,0.015] BF: 0.254 ROPE Bound: 0.002 In ROPE: 0.169 BF in ROPE: 4.54 RHat: 1.002 | Mode: 0.011 95% CI: [-0.018,0.025] BF: 0.22 ROPE Bound: 0.003 In ROPE: 0.188 BF in ROPE: 5.109 RHat: 1.003 | Mode: 0.014 95% CI: [-0.024,0.016] BF: 0.23 ROPE Bound: 0.002 In ROPE: 0.19 BF in ROPE: 5.202 RHat: 1.001 |
|  | Family: student Num. obs: 9808 Num. groups: 34 Bayes R2: 0.535 | Family: student Num. obs: 9808 Num. groups: 34 Bayes R2: 0.615 | Family: student Num. obs: 9808 Num. groups: 34 Bayes R2: 0.636 | Family: student Num. obs: 9808 Num. groups: 34 Bayes R2: 0.619 | Family: student Num. obs: 9808 Num. groups: 34 Bayes R2: 0.61 | Family: student Num. obs: 9808 Num. groups: 34 Bayes R2: 0.617 | Family: student Num. obs: 9808 Num. groups: 34 Bayes R2: 0.59 |

*Table S46.* The single-trial associations between the continuous measure of prediction error (PE) and the length of the current interbeat interval (IBI), for the three heartbeats before and the four heartbeats after feedback. ROPE – region of practical equivalence, BF – Bayes Factor. Cardiac phase is a continuous measure of distance from the last heartbeat before feedback to feedback, scaled by the distance from the last heartbeat before to the first heartbeat after feedback (IBI 1). All values are standardised. Note that RHat is > 1.01 for Intercept (but not other coefficients) in R-1 and R1 models, indicating convergence issues.

| Parameter | Dependent variable: IBI -3 | Dependent variable: IBI -2 | Dependent variable: IBI -1 | Dependent variable: IBI 1 | Dependent variable: IBI 2 | Dependent variable: IBI 3 | Dependent variable: IBI 4 |
| --- | --- | --- | --- | --- | --- | --- | --- |
| Intercept | Mode: 0.203 95% CI: [-0.214,0.338] BF: 0.048 ROPE Bound: 0.008 In ROPE: 0.044 BF in ROPE: 19.368 RHat: 1.006 | Mode: -0.168 95% CI: [-0.243,0.354] BF: 0.056 ROPE Bound: 0.008 In ROPE: 0.041 BF in ROPE: 18.45 RHat: 1.008 | Mode: 0.141 95% CI: [-0.282,0.352] BF: 0.066 ROPE Bound: 0.008 In ROPE: 0.04 BF in ROPE: 16.556 RHat: 1.011 | Mode: 0.171 95% CI: [-0.219,0.357] BF: 0.058 ROPE Bound: 0.008 In ROPE: 0.041 BF in ROPE: 18.68 RHat: 1.014 | Mode: 0.127 95% CI: [-0.254,0.351] BF: 0.067 ROPE Bound: 0.009 In ROPE: 0.041 BF in ROPE: 15.465 RHat: 1.02 | Mode: 0.223 95% CI: [-0.235,0.334] BF: 0.061 ROPE Bound: 0.009 In ROPE: 0.042 BF in ROPE: 18.243 RHat: 1.007 | Mode: 0.11 95% CI: [-0.246,0.317] BF: 0.056 ROPE Bound: 0.009 In ROPE: 0.047 BF in ROPE: 19.487 RHat: 1.007 |
| PE | Mode: 0.004 95% CI: [-0.015,0.006] BF: 0.171 ROPE Bound: 0.002 In ROPE: 0.201 BF in ROPE: 7.115 RHat: 1.002 | Mode: -0.004 95% CI: [-0.007,0.013] BF: 0.106 ROPE Bound: 0.002 In ROPE: 0.296 BF in ROPE: 12.711 RHat: 1.003 | Mode: -0.007 95% CI: [-0.009,0.012] BF: 0.111 ROPE Bound: 0.002 In ROPE: 0.32 BF in ROPE: 12.469 RHat: 1.001 | Mode: 0.011 95% CI: [-0.001,0.021] BF: 0.713 ROPE Bound: 0.002 In ROPE: 0.049 BF in ROPE: 1.693 RHat: 1.001 | Mode: -0.015 95% CI: [-0.019,0.004] BF: 0.305 ROPE Bound: 0.002 In ROPE: 0.128 BF in ROPE: 3.672 RHat: 1.002 | Mode: -0.022 95% CI: [-0.027,-0.005] BF: 5.44 ROPE Bound: 0.002 In ROPE: 0 BF in ROPE: 0.207 RHat: 1.003 | Mode: -0.013 95% CI: [-0.017,0.004] BF: 0.249 ROPE Bound: 0.002 In ROPE: 0.152 BF in ROPE: 4.598 RHat: 1.001 |
| Cardiac phase | Mode: -0.014 95% CI: [-0.027,-0.006] BF: 14.199 ROPE Bound: 0.002 In ROPE: 0 BF in ROPE: 0.088 RHat: 1.002 | Mode: -0.014 95% CI: [-0.016,0.004] BF: 0.198 ROPE Bound: 0.002 In ROPE: 0.192 BF in ROPE: 6.238 RHat: 1.002 | Mode: 0.012 95% CI: [0.001,0.022] BF: 1.059 ROPE Bound: 0.002 In ROPE: 0.013 BF in ROPE: 0.951 RHat: 1.002 | Mode: -0.02 95% CI: [-0.034,-0.012] BF: 532 ROPE Bound: 0.002 In ROPE: 0 BF in ROPE: 0.002 RHat: 1.001 | Mode: 0.034 95% CI: [0.026,0.047] BF: 21325 ROPE Bound: 0.002 In ROPE: 0 BF in ROPE: 0 RHat: 1.001 | Mode: 0.031 95% CI: [0.026,0.047] BF: 126735 ROPE Bound: 0.003 In ROPE: 0 BF in ROPE: 0 RHat: 1.001 | Mode: 0.02 95% CI: [0.016,0.036] BF: 2595 ROPE Bound: 0.002 In ROPE: 0 BF in ROPE: 0 RHat: 1.001 |
| PE : Cardiac phase | Mode: -0.007 95% CI: [-0.012,0.009] BF: 0.118 ROPE Bound: 0.001 In ROPE: 0.092 BF in ROPE: 9.79 RHat: 1.001 | Mode: -0.01 95% CI: [-0.014,0.006] BF: 0.141 ROPE Bound: 0.001 In ROPE: 0.073 BF in ROPE: 7.264 RHat: 1.001 | Mode: -0.007 95% CI: [-0.014,0.007] BF: 0.131 ROPE Bound: 0.001 In ROPE: 0.077 BF in ROPE: 8.176 RHat: 1.001 | Mode: -0.01 95% CI: [-0.017,0.005] BF: 0.212 ROPE Bound: 0.001 In ROPE: 0.043 BF in ROPE: 4.927 RHat: 1.002 | Mode: 0.008 95% CI: [-0.005,0.018] BF: 0.208 ROPE Bound: 0.001 In ROPE: 0.056 BF in ROPE: 4.909 RHat: 1.001 | Mode: 0.002 95% CI: [-0.01,0.011] BF: 0.11 ROPE Bound: 0.001 In ROPE: 0.108 BF in ROPE: 10.07 RHat: 1.003 | Mode: 0.002 95% CI: [-0.006,0.015] BF: 0.154 ROPE Bound: 0.001 In ROPE: 0.063 BF in ROPE: 7.31 RHat: 1.002 |
|  | Family: student Num. obs: 9808 Num. groups: 34 Bayes R2: 0.535 | Family: student Num. obs: 9808 Num. groups: 34 Bayes R2: 0.615 | Family: student Num. obs: 9808 Num. groups: 34 Bayes R2: 0.636 | Family: student Num. obs: 9808 Num. groups: 34 Bayes R2: 0.62 | Family: student Num. obs: 9808 Num. groups: 34 Bayes R2: 0.61 | Family: student Num. obs: 9808 Num. groups: 34 Bayes R2: 0.616 | Family: student Num. obs: 9808 Num. groups: 34 Bayes R2: 0.59 |

*Table S47.* The single-trial associations between the continuous measure of unsigned prediction error (unsigned PE) and the length of the current interbeat interval (IBI), for the three heartbeats before and the four heartbeats after feedback. ROPE – region of practical equivalence, BF – Bayes Factor. Cardiac phase is a continuous measure of distance from the last heartbeat before feedback to feedback, scaled by the distance from the last heartbeat before to the first heartbeat after feedback (IBI 1). All values are standardised. Note that RHat is > 1.01 for Intercept (but not other coefficients) in R-3, R-2, R1 and R2 models, indicating convergence issues.

| Parameter | Dependent variable: IBI -3 | Dependent variable: IBI -2 | Dependent variable: IBI -1 | Dependent variable: IBI 1 | Dependent variable: IBI 2 | Dependent variable: IBI 3 | Dependent variable: IBI 4 |
| --- | --- | --- | --- | --- | --- | --- | --- |
| Intercept | Mode: 0.115 95% CI: [-0.23,0.317] BF: 0.05 ROPE Bound: 0.008 In ROPE: 0.047 BF in ROPE: 20.135 RHat: 1.021 | Mode: -0.105 95% CI: [-0.227,0.414] BF: 0.064 ROPE Bound: 0.008 In ROPE: 0.037 BF in ROPE: 16.806 RHat: 1.017 | Mode: -0.134 95% CI: [-0.27,0.367] BF: 0.059 ROPE Bound: 0.008 In ROPE: 0.04 BF in ROPE: 17.477 RHat: 1.009 | Mode: -0.131 95% CI: [-0.246,0.392] BF: 0.067 ROPE Bound: 0.008 In ROPE: 0.041 BF in ROPE: 14.749 RHat: 1.022 | Mode: 0.221 95% CI: [-0.275,0.342] BF: 0.066 ROPE Bound: 0.009 In ROPE: 0.042 BF in ROPE: 15.812 RHat: 1.011 | Mode: -0.153 95% CI: [-0.241,0.317] BF: 0.051 ROPE Bound: 0.009 In ROPE: 0.048 BF in ROPE: 19.091 RHat: 1.006 | Mode: 0.108 95% CI: [-0.219,0.326] BF: 0.057 ROPE Bound: 0.009 In ROPE: 0.047 BF in ROPE: 18.513 RHat: 1.009 |
| Unsigned PE | Mode: -0.01 95% CI: [-0.015,0.006] BF: 0.133 ROPE Bound: 0.003 In ROPE: 0.316 BF in ROPE: 10.239 RHat: 1.003 | Mode: -0.011 95% CI: [-0.019,0.001] BF: 0.407 ROPE Bound: 0.003 In ROPE: 0.111 BF in ROPE: 2.717 RHat: 1.002 | Mode: -0.002 95% CI: [-0.01,0.01] BF: 0.11 ROPE Bound: 0.003 In ROPE: 0.418 BF in ROPE: 13.821 RHat: 1.001 | Mode: -0.01 95% CI: [-0.011,0.011] BF: 0.109 ROPE Bound: 0.003 In ROPE: 0.438 BF in ROPE: 14.71 RHat: 1.003 | Mode: 0.023 95% CI: [0.007,0.029] BF: 14.324 ROPE Bound: 0.003 In ROPE: 0 BF in ROPE: 0.084 RHat: 1.002 | Mode: 0.028 95% CI: [0.011,0.033] BF: 120 ROPE Bound: 0.003 In ROPE: 0 BF in ROPE: 0.012 RHat: 1.001 | Mode: 0.014 95% CI: [0,0.021] BF: 0.946 ROPE Bound: 0.003 In ROPE: 0.043 BF in ROPE: 1.327 RHat: 1.001 |
| Cardiac phase | Mode: -0.022 95% CI: [-0.027,-0.006] BF: 8.979 ROPE Bound: 0.002 In ROPE: 0 BF in ROPE: 0.126 RHat: 1.001 | Mode: -0.004 95% CI: [-0.016,0.005] BF: 0.188 ROPE Bound: 0.002 In ROPE: 0.203 BF in ROPE: 6.635 RHat: 1.001 | Mode: 0.013 95% CI: [0.002,0.022] BF: 1.46 ROPE Bound: 0.002 In ROPE: 0.008 BF in ROPE: 0.752 RHat: 1.002 | Mode: -0.018 95% CI: [-0.035,-0.013] BF: 395 ROPE Bound: 0.002 In ROPE: 0 BF in ROPE: 0.003 RHat: 1.002 | Mode: 0.041 95% CI: [0.026,0.048] BF: 200112 ROPE Bound: 0.002 In ROPE: 0 BF in ROPE: 0 RHat: 1.001 | Mode: 0.032 95% CI: [0.026,0.047] BF: 118513 ROPE Bound: 0.003 In ROPE: 0 BF in ROPE: 0 RHat: 1.002 | Mode: 0.023 95% CI: [0.015,0.036] BF: 2602 ROPE Bound: 0.002 In ROPE: 0 BF in ROPE: 0 RHat: 1.001 |
| Unsigned PE : Cardiac phase | Mode: -0.002 95% CI: [-0.016,0.005] BF: 0.152 ROPE Bound: 0.001 In ROPE: 0.096 BF in ROPE: 6.989 RHat: 1.002 | Mode: -0.003 95% CI: [-0.012,0.009] BF: 0.117 ROPE Bound: 0.001 In ROPE: 0.117 BF in ROPE: 9.691 RHat: 1.001 | Mode: 0.004 95% CI: [-0.011,0.01] BF: 0.102 ROPE Bound: 0.001 In ROPE: 0.126 BF in ROPE: 10.592 RHat: 1.001 | Mode: -0.002 95% CI: [-0.009,0.013] BF: 0.12 ROPE Bound: 0.001 In ROPE: 0.108 BF in ROPE: 9.263 RHat: 1.001 | Mode: -0.011 95% CI: [-0.019,0.003] BF: 0.373 ROPE Bound: 0.001 In ROPE: 0.041 BF in ROPE: 2.764 RHat: 1.001 | Mode: 0.003 95% CI: [-0.01,0.011] BF: 0.12 ROPE Bound: 0.001 In ROPE: 0.14 BF in ROPE: 10.118 RHat: 1.001 | Mode: -0.011 95% CI: [-0.014,0.008] BF: 0.128 ROPE Bound: 0.001 In ROPE: 0.128 BF in ROPE: 8.888 RHat: 1.001 |
|  | Family: student Num. obs: 9808 Num. groups: 34 Bayes R2: 0.535 | Family: student Num. obs: 9808 Num. groups: 34 Bayes R2: 0.615 | Family: student Num. obs: 9808 Num. groups: 34 Bayes R2: 0.636 | Family: student Num. obs: 9808 Num. groups: 34 Bayes R2: 0.619 | Family: student Num. obs: 9808 Num. groups: 34 Bayes R2: 0.61 | Family: student Num. obs: 9808 Num. groups: 34 Bayes R2: 0.617 | Family: student Num. obs: 9808 Num. groups: 34 Bayes R2: 0.59 |

#### 4.5. Models with and without random slopes

Here we present the same analyses as for the main findings but with random slopes and intercepts instead of intercepts-only. The findings and directions of associations are the same as in the main analyses, only the strength of evidence is reduced due to lower degrees of freedom (Tables S48, S49, S50). We compared the model fits with and without random slopes, and while for some models the fits were slightly better with random slopes, these differences were not strong (Table S51).

##### 4.5.1. Model results with random slopes

*Table S48.* The single-trial associations between the continuous measure of expectation and the degree of change of the current interbeat interval (IBI) compared to the previous one, for the last heartbeat before and the third heartbeat after feedback. ROPE – region of practical equivalence, BF – Bayes Factor. Cardiac phase is a continuous measure of distance from the last heartbeat before feedback to feedback, scaled by the distance from the last heartbeat before to the first heartbeat after feedback (IBI 1). All values are standardised. Models include random slopes.

| Parameter | Dependent variable: Change of IBI -1 | Dependent variable: Change of IBI 3 |
| --- | --- | --- |
| Intercept | Mode: -0.028 95% CI: [-0.045,0.063] BF: 0.011 ROPE Bound: 0.018 In ROPE: 0.499 BF in ROPE: 165 | Mode: 0.037 95% CI: [-0.057,0.119] BF: 0.02 ROPE Bound: 0.02 In ROPE: 0.295 BF in ROPE: 67.496 |
| Expectation | Mode: -0.023 95% CI: [-0.05,-0.012] BF: 16.281 ROPE Bound: 0.006 In ROPE: 0 BF in ROPE: 0.069 | Mode: 0.032 95% CI: [0.015,0.053] BF: 56.168 ROPE Bound: 0.006 In ROPE: 0 BF in ROPE: 0.022 |
| Cardiac phase | Mode: 0.033 95% CI: [0.019,0.062] BF: 68.956 ROPE Bound: 0.005 In ROPE: 0 BF in ROPE: 0.017 | Mode: 0.02 95% CI: [-0.018,0.031] BF: 0.277 ROPE Bound: 0.006 In ROPE: 0.335 BF in ROPE: 4.714 |
| Expectation : Cardiac phase | Mode: -0.004 95% CI: [-0.02,0.013] BF: 0.176 ROPE Bound: 0.002 In ROPE: 0.146 BF in ROPE: 6.25 | Mode: 0.007 95% CI: [-0.019,0.016] BF: 0.173 ROPE Bound: 0.002 In ROPE: 0.158 BF in ROPE: 6.375 |
|  | Family: student Num. obs: 9808 Num. groups: 34 Bayes R2: 0.024 | Family: student Num. obs: 9808 Num. groups: 34 Bayes R2: 0.057 |

*Table S49.* The single-trial associations between the valence of outcome (contrast: lose compared to win), continuous measures of prediction error (PE) or unsigned PE and the degree of change of the current interbeat interval (IBI) compared to the previous one, for the second heartbeat after feedback. ROPE – region of practical equivalence, BF – Bayes Factor. Cardiac phase is a continuous measure of distance from the last heartbeat before feedback to feedback, scaled by the distance from the last heartbeat before to the first heartbeat after feedback (IBI 1). All values are standardised. Models include random slopes.

| Parameter | Dependent variable: Change of IBI 2 | Dependent variable: Change of IBI 2 | Dependent variable: Change of IBI 2 |
| --- | --- | --- | --- |
| Intercept | Mode: -0.025 95% CI: [-0.047,0.078] BF: 0.012 ROPE Bound: 0.019 In ROPE: 0.428 BF in ROPE: 125 | Mode: 0.08 95% CI: [-0.026,0.093] BF: 0.021 ROPE Bound: 0.019 In ROPE: 0.291 BF in ROPE: 66.591 | Mode: 0.032 95% CI: [-0.028,0.096] BF: 0.021 ROPE Bound: 0.019 In ROPE: 0.301 BF in ROPE: 74.014 |
| Outcome Lose | Mode: 0.045 95% CI: [0.003,0.077] BF: 4.502 ROPE Bound: 0.019 In ROPE: 0.085 BF in ROPE: 0.27 |  |  |
| Cardiac phase | Mode: 0.131 95% CI: [0.083,0.161] BF: 45754 ROPE Bound: 0.005 In ROPE: 0 BF in ROPE: 0 | Mode: 0.1 95% CI: [0.072,0.146] BF: 3730 ROPE Bound: 0.005 In ROPE: 0 BF in ROPE: 0 | Mode: 0.12 95% CI: [0.071,0.146] BF: 4824 ROPE Bound: 0.005 In ROPE: 0 BF in ROPE: 0 |
| Outcome Lose : Cardiac phase | Mode: -0.017 95% CI: [-0.063,-0.002] BF: 3.519 ROPE Bound: 0.005 In ROPE: 0.007 BF in ROPE: 0.287 |  |  |
| PE |  | Mode: -0.021 95% CI: [-0.041,-0.006] BF: 8.186 ROPE Bound: 0.005 In ROPE: 0 BF in ROPE: 0.155 |  |
| PE : Cardiac phase |  | Mode: 0.035 95% CI: [0.011,0.042] BF: 27.378 ROPE Bound: 0.001 In ROPE: 0 BF in ROPE: 0.037 |  |
| Unsigned PE |  |  | Mode: 0.025 95% CI: [0.008,0.047] BF: 11.615 ROPE Bound: 0.007 In ROPE: 0 BF in ROPE: 0.125 |
| Unsigned PE : Cardiac phase |  |  | Mode: -0.021 95% CI: [-0.031,0.001] BF: 0.806 ROPE Bound: 0.002 In ROPE: 0.039 BF in ROPE: 1.283 |
|  | Family: student Num. obs: 9808 Num. groups: 34 Bayes R2: 0.048 | Family: student Num. obs: 9808 Num. groups: 34 Bayes R2: 0.048 | Family: student Num. obs: 9808 Num. groups: 34 Bayes R2: 0.049 |

*Table S50.* Separately only within systole or diastole trials (first and last 1/3 of the cardiac cycle), the single-trial associations between the continuous measure of P3b amplitude and the degree of change in the current interbeat interval (IBI) compared to the previous one, for the first and second heartbeats after feedback. ROPE – region of practical equivalence, BF – Bayes Factor. All values are standardised. Models include random slopes.

| Parameter | Dependent variable: Change of IBI 1 only in systole | Dependent variable: Change of IBI 2 only in diastole |
| --- | --- | --- |
| Intercept | Mode: 0.104 95% CI: [0.033,0.159] BF: 0.666 ROPE Bound: 0.018 In ROPE: 0 BF in ROPE: 1.756 | Mode: 0.137 95% CI: [0.111,0.204] BF: 786 ROPE Bound: 0.019 In ROPE: 0 BF in ROPE: 0.002 |
| P3b | Mode: 0.028 95% CI: [0.012,0.071] BF: 15.737 ROPE Bound: 0.003 In ROPE: 0 BF in ROPE: 0.066 | Mode: 0.072 95% CI: [0.024,0.082] BF: 52.588 ROPE Bound: 0.003 In ROPE: 0 BF in ROPE: 0.017 |
|  | Family: student Num. obs: 3212 Num. groups: 34 Bayes R2: 0.027 | Family: student Num. obs: 3229 Num. groups: 34 Bayes R2: 0.015 |

##### 4.5.2. Model comparisons

*Table S51.* We compared the model fits with and without random slopes using leave-one-out cross-validation. For each Bayesian model, we extracted leave-one-out information criterion (LOOIC), effective number of parameters (pLOO), and pointwise expected log predictive density (ELPD). Model differences were evaluated based on the total ELPD difference between models as well as the standard error of these differences (SE diff). Positive ELPD difference between models with and without random slopes favours the random-slope model, indicating better out-of-sample predictive performance. However, small ELPD values may reflect estimation uncertainty, so we interpret only the differences that exceed three SE as meaningful. For all models, we found no strong differences. However, two models (marked with * in Conclusion) showed the largest differences that could be meaningful with a threshold of two SE.

| Model | LOOIC with slopes | LOOIC without slopes | pLOO with | pLOO without | LOOIC diff (without - with) | ELPD diff (with - without) | SE diff | Conclusion |
| --- | --- | --- | --- | --- | --- | --- | --- | --- |
| R-1 and Expectation | 26,451.46 | 26,451.14 | 62.92 | 42.03 | -0.32 | -0.16 | 2.76 | No meaningful difference |
| R3 and Expectation | 26,471.97 | 26,481.25 | 71.54 | 46.73 | 9.27 | 4.64 | 4.36 | No meaningful difference |
| R2 and Outcome | 25,670.04 | 25,742.49 | 88.13 | 43.49 | 72.45 | 36.22 | 12.37 | No meaningful difference* |
| R2 and PE | 25,667.84 | 25,736.94 | 82.19 | 43.77 | 69.10 | 34.55 | 12.03 | No meaningful difference* |
| R1 and P3b in systole | 8,558.97 | 8,557.14 | 38.84 | 34.68 | -1.83 | -0.92 | 0.62 | No meaningful difference |
| R2 and P3b in Diastole | 8,371.37 | 8,369.68 | 30.53 | 26.27 | -1.70 | -0.85 | 0.62 | No meaningful difference |

### 5. Computational modelling results

#### 5.1. Model fit

*Figure S4.* (A, B, C, D) Estimated parameters in the study. Alpha, alpha_neg, and alpha_pos are learning rates, fitted in the range from 0.01 to 0.99. Beta is the inverse temperature representing the degree of exploration and fitted in the range from 0.01 to 10. For RW, alpha was fitted at upper bound for 5 participants. For AC, alpha was fitted at upper and lower bounds for 1 participant, and beta at lower bound for 1 participant. For CB, alpha_pos was fitted at upper bound for 4 participants, alpha_neg at upper for 1 and at lower for 4 participants. Maximum beta across all models was 4.5 for RW model. (E) The number of participants assigned the best-fitting model based on BIC across all five models, including two control models. CB leads with 14 participants. Random responding was not assigned to any participant, outperformed by other models. BIC - Bayesian Information Criterion, rw - Rescorla-Wagner model, ac_rw - anticorrelated Rescorla-Wagner model, cb - Confirmation Bias model, wsls - noisy Win-Stay-Lose-Shift.

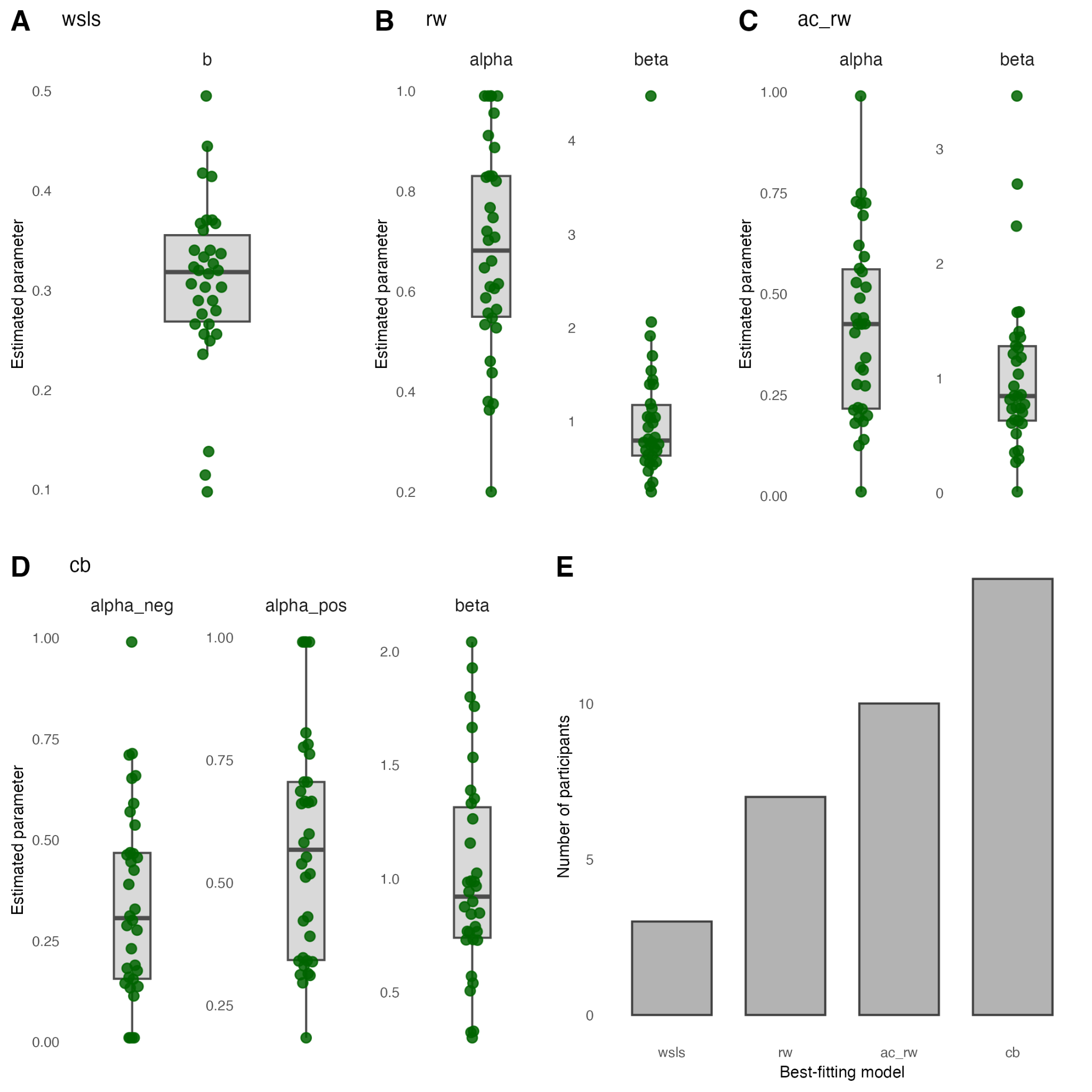

#### 5.2. Recovery and confusion

Parameter recovery was evaluated by simulating datasets across a grid of parameter values with 10 equally-spaced options per parameter, resulting in 100 simulations for RW and AC models, and 1000 simulations for the CB model. Then we fitted the generating model back to each simulated dataset using maximum-likelihood estimation and compared the generating and recovered parameters (Figure S11). We observed satisfactory recovery performance based on mean unsigned errors (MAE) and Pearson correlations for RW (alpha MAE = 0.15, alpha cor = 0.66, beta MAE = 1.82, beta cor = 0.71), AC (alpha MAE = 0.14, alpha cor = 0.79, beta MAE = 1.85, beta cor = 0.75), and CB (alpha_pos MAE = 0.23, alpha_pos cor = 0.6, alpha_neg MAE = 0.13, alpha_neg cor = 0.8, beta MAE = 1.93, beta cor = 0.75) models.

We then utilised the same simulations that cover the full grid of parameter values to construct a confusion matrix (Figure S11D). The confusion matrix reflects, for each combination of two models, the proportion across all simulations when the model that generated the data was chosen as the best fitting model based on Bayesian Information Criterion (BIC) or when another model was chosen as the best fitting one.

RW, CB, and WSLS are the best-fitting models when they were used for data generation. AC is the second best model, closely following WSLS. There is also a moderate degree of confusion between CB and AC or WSLS, as all of these models produce analogous simultaneous propensity for switching behavior in response to negative outcomes and stickiness after positive outcomes.

*Figure S5.* Simulation-based quality checks for the three main learning models (RW, AC, CB) and two control heuristic models (WSLS and random responding). (A, B, C) Parameter recovery: simulated parameter values against recovered ones, for a grid of parameter values (10 options per parameter). Alpha, alpha_neg, and alpha_pos are learning rates. Beta is the inverse temperature that represents the degree of exploration. Dashed lines mark perfect recovery. (D) Model confusion matrix that reflects how often data generating models (rows) were assigned as best-fitting models (columns) based on BIC. BIC - Bayesian Information Criterion, rw - Rescorla-Wagner model, ac_rw - anticorrelated Rescorla-Wagner model, cb - Confirmation Bias model, wsls - noisy Win-Stay-Lose-Shift, rand_resp - random responding.

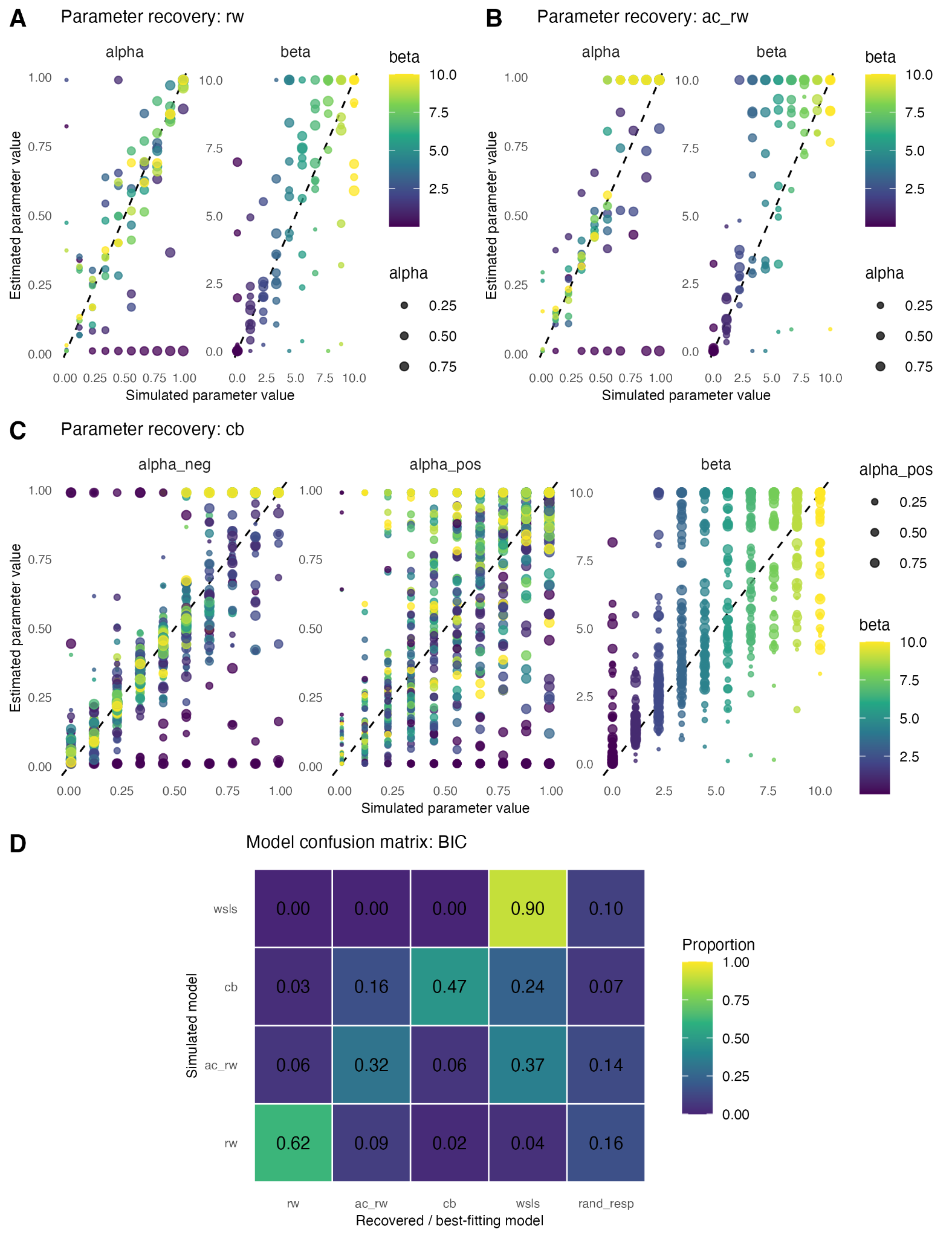

### 6. Distributional diagnostics

For Bayesian model fitting, we chose a likelihood family objectively based on the distribution of dependent variables and a series of statistical tests. Student-t is the default. First, approximate uniformity is defined as no significant deviation from a uniform distribution in either the Kolmogorov-Smirnov (KS) or Cramer-von Mises (CvM) tests, p-value > 0.05, together with unsigned skewness < 0.10 and kurtosis between 1.5 and 2.1. If the variable met uniformity criteria, we used beta (if bounded between [0,1]) or the default family. Second, we conducted normality tests using Shapiro-Wilk (if number of observations ≤ 5000) or Anderson-Darling (otherwise), alongside a KS test for normality. If both tests failed to reject normality, the Gaussian was selected. If normality was rejected and kurtosis exceeded 3, we compared Gaussian and Student-t models via Akaike Information Criterion (AIC). If Student-t provided a better fit, we classified the variable as heavy-tailed and used a Student-t distribution. If heavy tails were absent, we performed skewness testing using a skew-normal fit. If skewness was statistically significant (p-value < 0.05) and unsigned skewness exceeded 0.5, we selected the skew-normal likelihood. Otherwise, we used Student-t.
